# An in vivo and in vitro spatiotemporal atlas of human midbrain development

**DOI:** 10.1101/2024.09.25.613908

**Authors:** Dimitri Budinger, Pau Puigdevall Costa, George T. Hall, Charlotte Roth, Theodoros Xenakis, Elena Marrosu, Julie Jerber, Alessandro Di Domenico, Helena Kilpinen, Sergi Castellano, Serena Barral, Manju A. Kurian

## Abstract

The dopaminergic system has key roles in human physiology and is implicated in a broad range of neurological and neuropsychiatric conditions that are increasingly investigated using induced pluripotent stem cell-derived midbrain models. To determine the similarity of such models to human systems, we undertook single cell and spatial profiling of first and second trimester fetal midbrain and compared it to in vitro midbrain models. Our initial histological analysis of second trimester fetal midbrain revealed structural complexity already similar to that of adult tissue, although this similarity did not fully extend to transcriptional activity. Moreover, we show that in vitro models recapitulate the transcriptional activity of late first trimester fetal midbrain, while 3D models replicate the spatial organization and cellular microenvironments of first and second trimester fetal midbrain. Understanding the extent of human tissue recapitulation in midbrain laboratory models is essential to justify their use as biological proxies.

## Introduction

The midbrain is a complex brain region comprising nuclei, nerves and tracts involved in vision, sensory processing, cognitive function and motor control (Parraga et al. 2016). In this composite system, dopaminergic nuclei, located ventrally to the cerebral aqueduct, play an important role in reward, cognition and control of voluntary movement, throughout the ventral tegmental area (VTA) and substantia nigra (SN) (Garritsen et al. 2023). Dysfunction of the midbrain dopaminergic system has been associated with a wide spectrum of neurological and neuropsychiatric conditions, including Parkinson’s disease (PD), attention deficit hyperactivity disorder, autism, drug addiction, and childhood neurological diseases (Kurian et al. 2011, Bissonette and Roesch 2016).

Despite conservation of several midbrain characteristics between species, the human midbrain has unique dopaminergic populations and developmental characteristics (Nelander et al. 2009, La Manno et al. 2016, Birtele et al. 2022, Zagare et al. 2022). Recent advancements in single cell RNA sequencing (scRNA-seq) and spatial transcriptomics have enabled the generation of comprehensive anatomical atlases for both the developing and adult human brain (Eze et al. 2021, Yu et al. 2021, Braun et al. 2023, Li et al. 2023, Siletti et al. 2023). These human atlases have offered a detailed map of the cellular diversity and molecular properties in both fetal and adult midbrain. Despite these advances, a significant gap in knowledge remains, particularly regarding cellular diversity, spatiotemporal distribution, and the developmental trajectory in late embryonic to postnatal midbrain development.

In parallel, the advent of in vitro 2D models derived from induced pluripotent stem cells (iPSCs) has provided valuable insights into the human midbrain (Barral and Kurian 2016). These models are also relevant for studying neurological disorders associated with midbrain dysfunction (Gantner et al. 2020, Carola et al. 2021). However, they do not capture the complete spectrum of cellular interactions and 3D spatial organization that is characteristic of the human midbrain. In contrast, 3D midbrain-like organoids (MLO) more faithfully recapitulate the complexity of the human midbrain, where development of multiple cell types and structural organization more closely resembles the tissue organization observed during embryonic development (Jo et al. 2016, Nickels et al. 2020, Fiorenzano et al. 2021). Nevertheless, fundamental questions remain unanswered: at what stage do these iPSC-derived models deviate from the natural course of development and if there is deviation, how effectively can they model neurodevelopmental and late-onset neurological disorders?

To explore the cellular diversity and developmental trajectory of the human midbrain, we employed a multimodal approach to profile midbrain tissue from human fetal samples at 6-22 post conceptional weeks (PCW), using immunohistochemistry, scRNA-seq and spatial transcriptomics. To provide a comprehensive overview of the cellular landscape during this developmental window, we compared our findings to previously profiled human fetal and postmortem adult samples. Furthermore, we explored how this aligned with 2D and 3D iPSC-derived midbrain models, offering a valuable perspective on the relevance and fidelity of these in vitro systems to study human fetal midbrain development and model neurological disorders.

## Results

### Anatomical changes of the developing human ventral midbrain across the first and second gestational trimester

To investigate the spatiotemporal organization of the ventral midbrain during the first half of fetal human development, we conducted immunohistochemistry analysis on midbrain tissues from 6 to 22 PCW (n=5 donors) **(Table S1),** focusing our analysis on dopaminergic neurons. All images are available at high-resolution in the MRC-Wellcome Trust Human Developmental Biology Resource (HDBR) Atlas (www.hdbr.org).

The first gestational trimester (up to 12 PCW) represents the neurogenesis stage of mDA neurons (Almqvist et al. 1996, Nelander et al. 2009, Marklund et al. 2014, Tiklova et al. 2019, Birtele et al. 2022). Following regional specification of the neural tube, mDA progenitors express the transcription factors FOXA2 (Ferri et al. 2007) and LMX1A (Andersson et al. 2006), both required for mDA neuron development (**Supp Fig 1A,B**). At 6 PCW, proliferating mDA progenitors in the ventricular zone (VZ) express SOX2, Ki67 and OTX2 (**Supp Fig 1C-E**) (Nelander et al. 2009), while postmitotic mDA neurons express tyrosine hydroxylase (TH) and the microtubule-associated protein 2 (MAP2), in neurons located at the intermediate zone (IZ) (La Manno et al. 2016, Asgrimsdottir and Arenas 2020) (**Supp Fig 1F**). To further understand the spatiotemporal emergence of the SN and VTA, we investigated changes of the human midbrain during the second gestational trimester. We analyzed midbrain tissue from 13, 15, 19 and 22 PCW and detected a progressive increase in morphological complexity associated with the mDA neuron distribution over time (**Fig 1A; Supp Fig 2**). At 13 PCW the human midbrain is characterized by expansion of the ventral area (mantel zone) (**Fig 1B; Supp Fig 2A**). From 15 PCW, we could identify the emerging red nucleus (RN), lateral to the ventral midline, and the primordium of the SN and VTA within the mantel zone (MZ) of the basal plate, where TH+ mDA neurons start to position (**Supp Fig 2B**). During late stages of the second trimester, we observed increased complexity of the ventral midbrain and mDA nuclei organization. Further morphological specification of the SN and VTA was evident at 19 and 22 PCW. At 22 PCW, the ventral midbrain resembled the complex architecture of the adult tissue (Waldvogel et al. 2006) (**Fig 1C,D; Supp Fig 2C,D**). From 15 to 22 PCW, we observed synaptophysin (SYP) expression in the TH+ projections located in the SN and VTA (Garritsen et al. 2023) (**Supp Fig 3A**). We also observed the presence of mDA neurons expressing the neurotransmitter GABA, in line with recent transcriptomic observations (**Supp Fig 3A**) (Siletti et al. 2023).

**Fig. 1.**
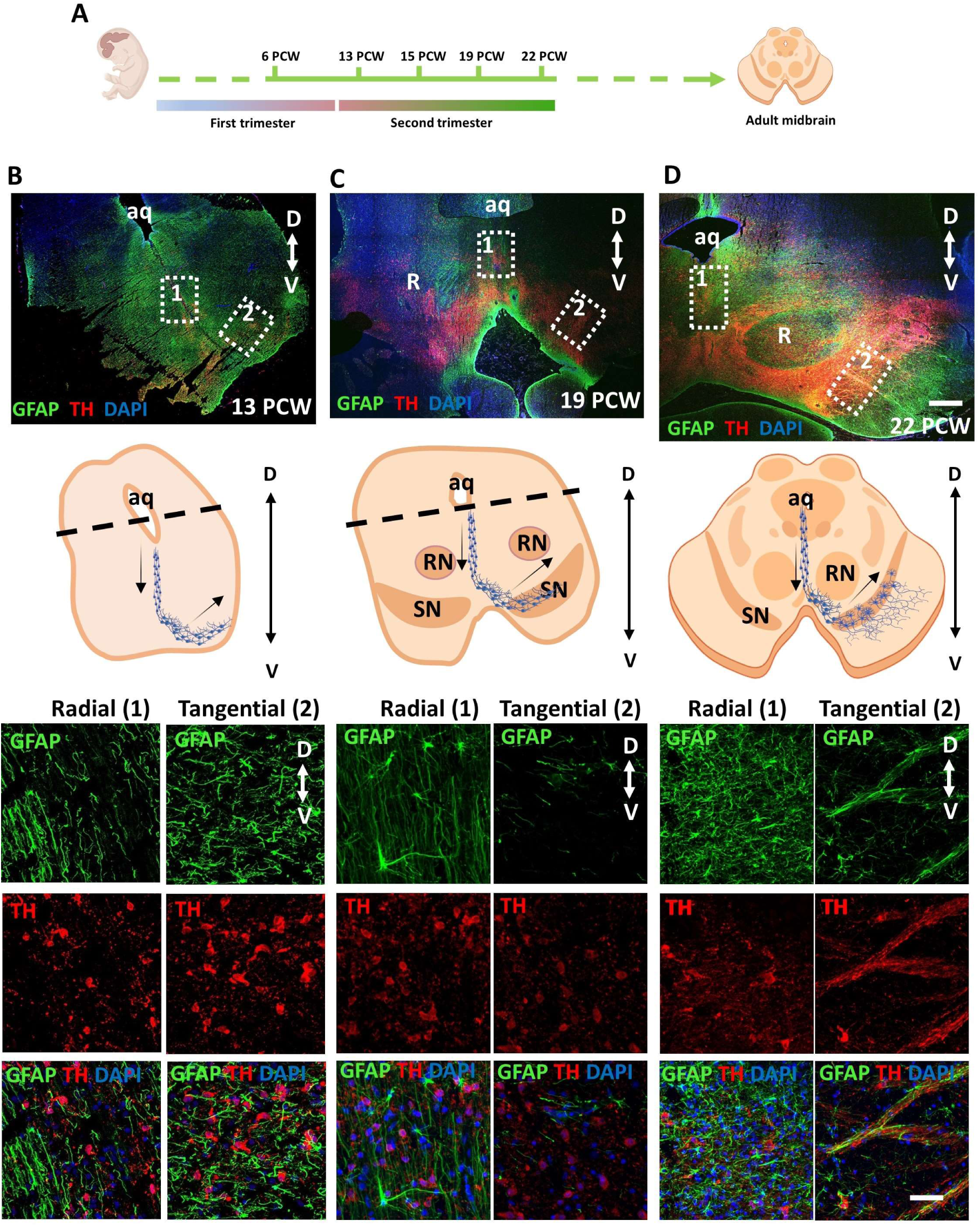
Morphological changes of the ventral human midbrain during the second trimester. **A** experimental design indicating age of fetal samples analysed. **B-C**, and **D**, immunofluorescence analysis of fetal midbrain tissue coronal sections collected at 13 (a), 19 (b) and 22 (c) pcw for TH and GFAP (*upper panel*). Nuclei are stained with DAPI. Scale bar= 500 μm. Graphical representation of the ventral midbrain coronal section and TH positive cells radial and tangential migration (*middle panel*). Higher magnification of radial and tangential migrating TH+ and GFAP+ neural cells at 13 (a), 19 (b) and 22 (c) pcw. Dotted lines demarcate zoomed radial (1) and tangential (2) neuronal migration regions. Cerebral aqueduct (aq), dorsal (D), ventral (V), red nucleus (RN), substantia nigra (SN).

**Fig 2.**
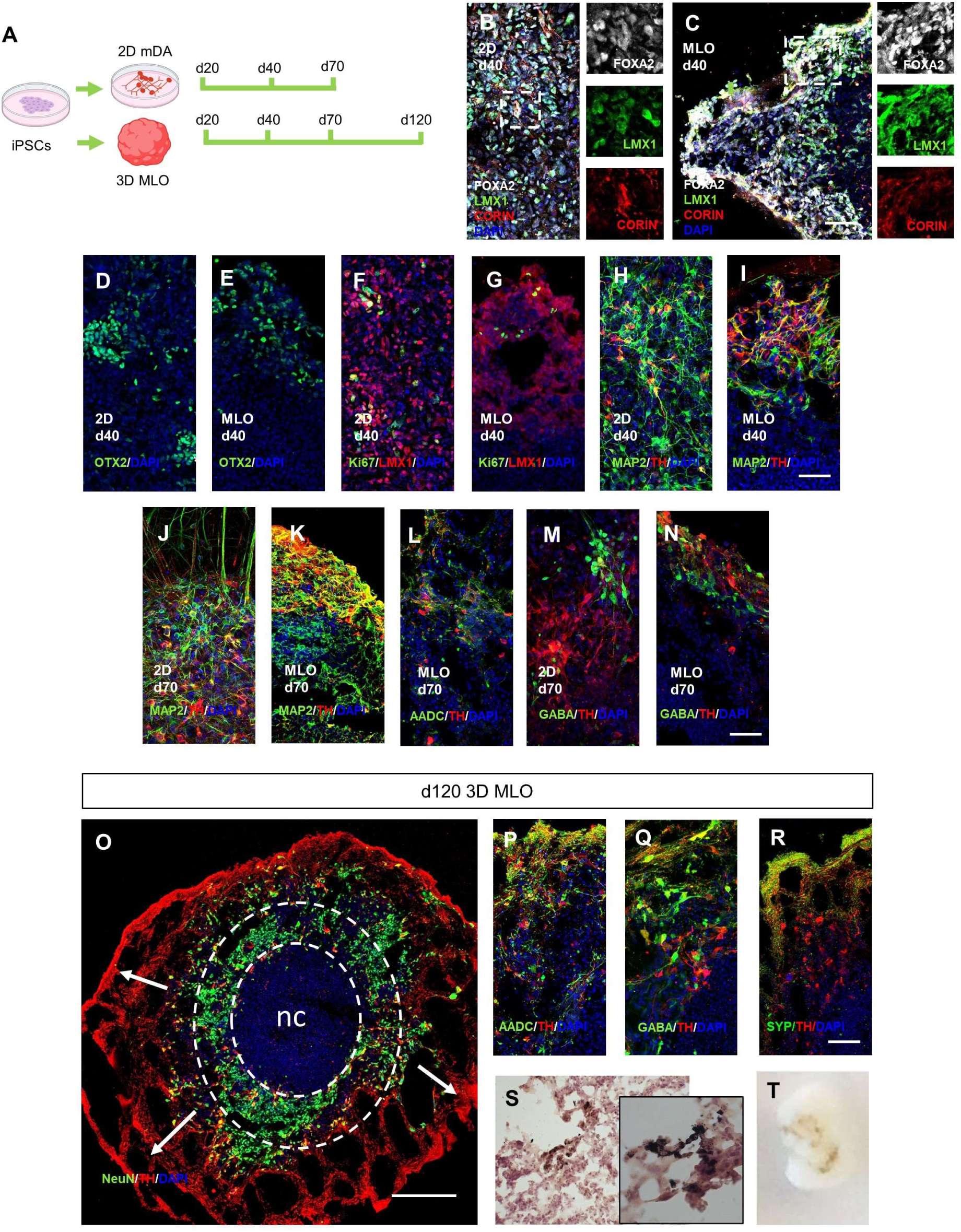
In vitro derived 2D mDA neurons and 3D MLO models cellular composition. **A**, experimental design for the *in vitro* system analysis. **B-I**, representative images of 2D mDA neuronal culture and MLO at 40. Cultures where stained for the midbrain-related and neuronal proteins FOXA2, LMX1, CORIN, OTX2, TH, MAP2, and proliferative marker Ki67. **J-N**, representative images of 2D mDA neuronal culture and MLOs at 70 days of differentiation showing expression of TH, MAP2, AADC and GABA. **O-R** Immunofluorescence analysis of MLOs at 120 days of differentiations. MLOs were stained for, TH, NEUN, AADC, GABA, and synaptophysin (SYP). Nuclei are staining with DAPI. Scale bar= 50 μm. **S**, Fontana-Masson staining and haematoxylin and eosin (H&E) stain of MLO at 120 days of differentiation showing melanin precipitation. **T**, Bright field image visualizing melanin aggregates.

**Fig. 3.**
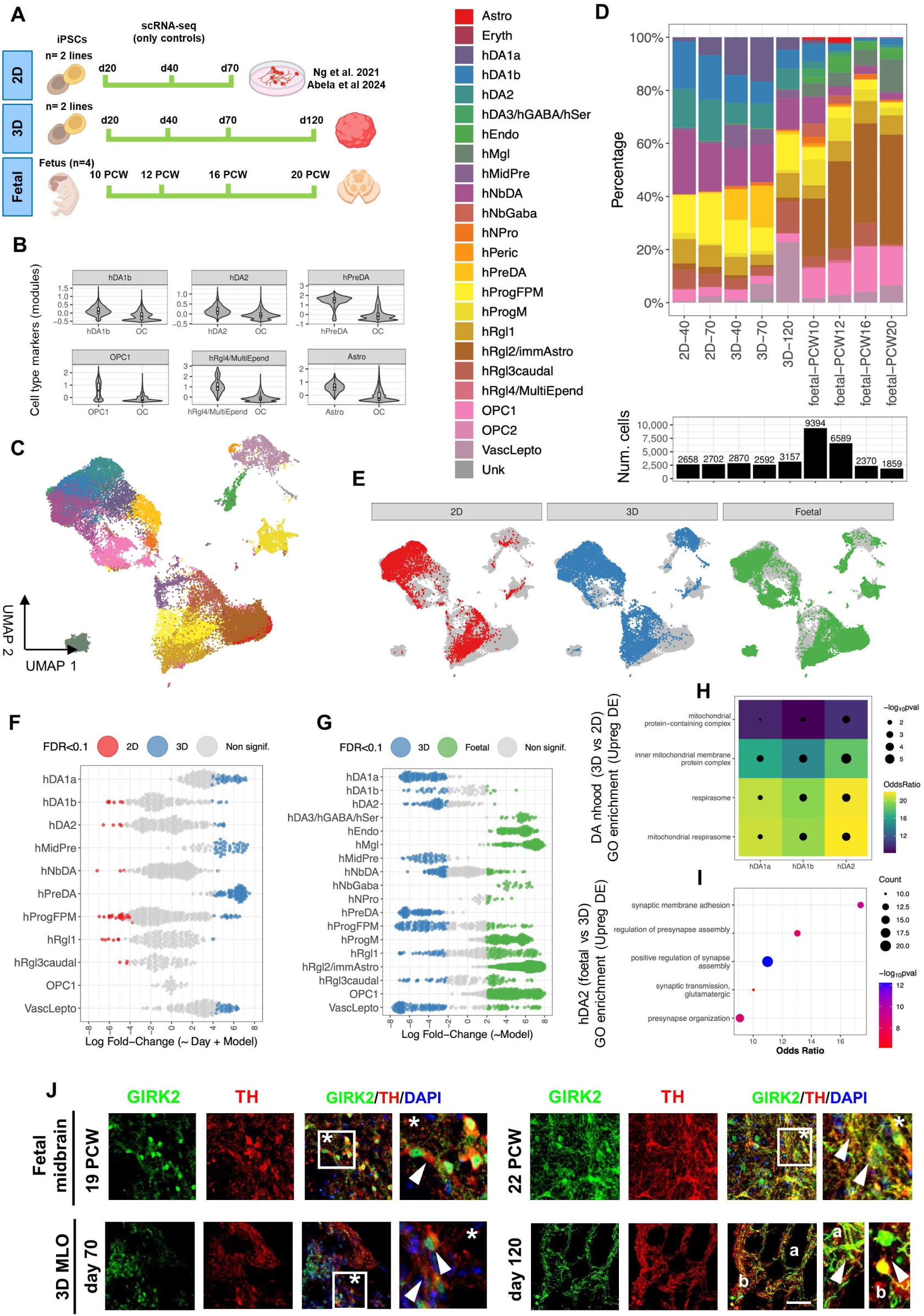
Single cell RNA-seq profiling of in vitro and in vivo foetal samples. **A**, Scheme of the in vitro (2D, 3D) and foetal samples profiled with droplet-based scRNA-seq. Only control samples were considered for all models: 2 iPSC lines from pediatric controls were differentiated to a midbrain dopaminergic fate, either as a 2D culture (days 40, 70) (Ng et al. 2021; Abela et al. 2024) or 3D organoids (days 40, 70, 120) (Li et al. 2016). As for fetal samples, the midbrain section of four donors were profiled at single timepoints: 10 and 12 PCW for the first trimester, and 16 and 20 PCW for the second trimester. **B**, The violin plots display the distribution of activation score for cell-type specific marker modules (Table S2), comparing the cell type of interest versus other cell types (OC). **C**, UMAP visualization based on gene expression with cells colored by the 24 annotated cell types. The largest cell clusters correspond either to neuronal cell population (top-left) or radial glia/floor-plate progenitors (bottom-right). **D**, Upper: cell type composition per model and time point. Lower: number of cells profiled in each corresponding group to estimate cell type composition. **E**, Same UMAP visualization (as in b), colored by in-vitro or in-vivo model. **F-G**, Differential abundance (DA) analysis comparing cell types between 3D vs 2D in-vitro models (f), and between fetal samples and the 3D model (g). Each point represents a neighborhood (group of cells connected by an edge in the KNN graph), with colored points indicating significant DA (SpatialFDR<0.1). Only in (f) is the sampling timepoint included as a covariate. **H**, The mitochondrial respirasome is commonly upregulated in 3D versus 2D models across neuronal populations (hDA1a, hDA1b and hDA2) indicating a prominent role of mitochondria metabolism in the maturation of dopaminergic neurons. Differential gene expression between 3D and 2D cells was calculated within the same cell type (neighborhood group), accounting for timepoints as a covariate (adj P.val<0.05, FC>1.75). **i,** Fetal hDA2 cells show an upregulation of synaptic-related pathways when compared with the 3D hDA2 cells (only the top-5 biological processes of the gene ontology enrichment are shown, pval<0.05, minSize=25). **J**, Immunofluorescence analysis of fetal midbrain at 19 and 22 PCW and MLOs at 70 and 120 days of differentiation for TH and the potassium channel GIRK2. Arrowheads indicate colocalization of GIRK2 and TH either in the soma or in the neurites. Nuclei are staining with DAPI. Scale bar= 50 μm.

Throughout the second trimester of gestation, we still observed mDA neurons located along the ventral midline migrating radially and not yet in their final location (**Fig 1B,C; Supp Fig 3B**) (Hanaway et al. 1971, Kawano et al. 1995, Hegarty et al. 2013, Arenas et al. 2015, Asgrimsdottir and Arenas 2020). We also detected GFAP+ cells exhibiting an elongated morphology and oriented in a dorsal-ventral direction, suggesting the presence of cells with radial glial or astrocytic identity. At 22 PCW, GFAP+ cells in the ventral midline lost their oriented morphology (**Fig 1D**). Both at 19 and 22 PCW, we observed mDA neurons undergoing tangential migration in the SN and VTA (Hegarty et al. 2013, Arenas et al. 2015, Asgrimsdottir and Arenas 2020) (**Fig 1C,D**). At 22 PCW, the tangential migration of mDA neurons was associated with GFAP+ cells. Radial and tangential migration were dependent on the maturation stage of mDA neurons. While at 19 PCW radial migrating mDA neurons did not express NEUN, a marker for neuronal maturity, it was expressed in radial migrating mDA neurons at 22 PCW, indicating increased neuronal maturity of these cells. In contrast, tangentially migrating mDA neurons expressed NEUN both at 19 and 22 PCW (**Supp Fig 3B**). Together, these observations indicate that across the first and second trimesters of gestation, the human midbrain undergoes major morphological changes, with increased complexity and neuronal maturity between 19 and 22 PCW, at which stage the fetal midbrain closely resembles the adult tissue (Johns 2014, Gupta 2017).

### Cellular and morphological complexity of iPSCs-derived midbrain models

We next investigated the alignment of in vitro human midbrain models to fetal development. 2D mDA neuronal cultures (Ng et al. 2021) and 3D midbrain-like organoids (MLO) (Kirkeby et al. 2012, Jo et al. 2016) were derived from two previously characterized human iPSC lines (Ng et al. 2021, Abela et al. 2024) (**Fig 2A**). At 20 days of differentiation, we confirmed efficient neuralization and ventral midbrain identity, with mDA progenitors expressing both FOXA2 and LMX1A. MLOs had an expected central necrotic core where no FOXA2*/*LMX1A positive cells were detected (Lancaster et al. 2013, Monzel et al. 2017). LMX1A+ cells also expressed CORIN, SOX2 and the cell cycle marker Ki67, further confirming progenitor midbrain identity. At 20 days of differentiation, we also observed MAP2+/TH+ cells in both 2D and 3D culture, suggesting that a fraction of mDA neurons have already reached post-mitotic maturation stage (**Supp Fig 4; Supp Fig 5**). At 40 days of differentiation, FOXA2, LMX1A, OTX2 and SOX2 expression was maintained in mDA progenitors in both 2D and 3D models (**Fig 2B-E; Supp Fig 4; Supp Fig 5**), while we observed fewer Ki67+ cells (**Fig 2F,G; Supp Fig 4; Supp Fig 5**). MAP2+/TH+ cells were present in both 2D and MLOs (**Fig 2H,I; Supp Fig 4; Supp Fig 5**). At 70 days, co-expression of MAP2, AADC and TH, NEUN, indicated the presence of post-mitotic mature mDA neurons (**Fig 2J-L; Supp Fig 4; Supp Fig 5**). We also observed the presence of neurons expressing the GABAergic neuronal marker GABA, and cells expressing GFAP (**Fig 2M,N; Supp Fig 4; Supp Fig 5**). 2D mDA neurons were maintained for a maximum of 70 days, whereas MLO differentiation allowed for an extended culture of up to 120 days. At this later stage of MLO differentiation, we observed the presence of NEUN+ and TH+ mDA neurons, which projected neurites outwards of the organoids (**Fig 2O**). We also observed expression of mature midbrain markers including AADC, GABA and SYP (**Fig 2P-R; Supp Fig 6**). Gene expression of *LMX1B*, *EN1*, *EN2*, *OTX2*, *NURR1*, *TH*, *PITX3*, *DAT* and *SNCA* also confirmed midbrain progenitor identity and mDA maturation over time and across both 2D mDA neurons and MLO cultures (**Supp Fig 6K**). Fontana-Masson staining revealed the presence of intracellular and extracellular neuromelanin, visible as a dark granular pigmentation, suggesting the specification and mature profile of mDA neurons of the SN pars compacta in these organoids (**Figure 2S,T**). Thus, our iPSC-derived mDA neuronal 2D and 3D systems recapitulate the major developmental steps of human midbrain development and can be harnessed to model these processes.

**Fig. 4.**
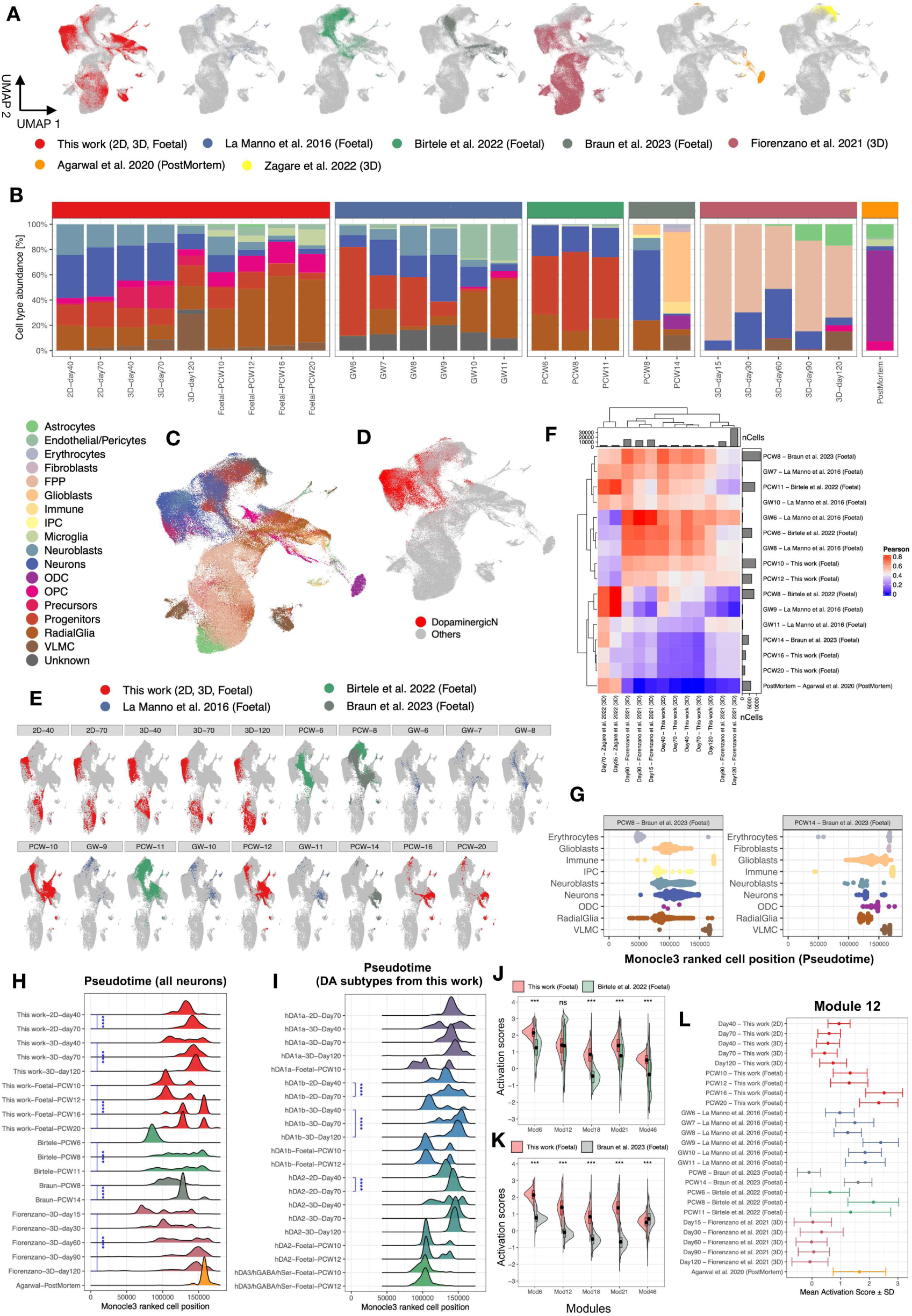
Meta-integration of multiple dopaminergic scRNA-seq datasets and analysis of pseudotemporal dynamics. **A**, UMAP visualization based on the integrated gene expression (MNN correction) of our dataset with in-vitro 3D dopaminergic models (Fiorenzano et al. 2021, Zagare et al. 2022), in-vivo fetal data (La Manno et al. 2016, Birtele et al. 2022 and Braun et al. 2023), and adult post-mortem samples (Zagare et al. 2022). The final integrated dataset contains 179,428 cells. **B**, Cell type composition per dataset and time point, expressed in percentage. For comparability between datasets, a unified cell type annotation (18 identities) has been defined (legend on 3^rd^ row). Abbrev: floor plate progenitors (FPP), intermediate progenitor cells (IPC), oligodendrocytes (ODC), oligodendrocyte progenitor cells (OPC) and vascular leptomeningeal cells (VLMC). **C-D**, UMAP visualization from (a) colored by either cell types (unified annotation) or just dopaminergic neurons, respectively. **E**, UMAP visualization from (a) colored by dataset and timepoint. Only cells collected from this work and foetal samples are highlighted. **F**, Transcriptomic similarity (Pearson’s correlation of MNN corrected gene expression, n=2,000 features) between fetal gestational timepoints and in vitro 3D dopaminergic differentiation timepoints. Abbrev: gestational week (GW), post-conceptional week (PCW). **G**, Violin scatter plots depicting the inferred Monocle3 pseudotime distribution per cell type (Braun et al. 2023) for post-conceptional weeks 8 and 14. The x axis indicates the ranked cell position across the integrated dataset as a surrogate for pseudotime. **H**, Ridge density plot of the pseudotime distribution for neurons, colored per dataset and displayed per timepoint (each row represents a different timepoint). Significance level: p<10^-4^ (****, Wilcoxon test). **I**, Ridge density plot of the pseudotime distribution for dopaminergic neurons subtypes annotated in this study. The plot is colored by subtype and displayed per timepoint (each row represents a different timepoint), shown only when enough cells (>50) are available per combination. Significance level: p<10^-4^ (****, Wilcoxon test). **J-K**, Split violin plots showing activation scores (aggregated gene expression) for the five neuron-specific pseudotime-dependent gene modules, comparing fetal data from this study with either Birtele et al. 2022 (f) or with Braun et al. 2023 (k), respectively. Significance level: p<10^-3^ (***, One-way permutation test). **L**, Activation score distribution for the neuron-specific module 12, colored by dataset and displayed per timepoint (each row represents a different timepoint). The plot includes mean values with standard deviation error bars.

**Fig. 5.**
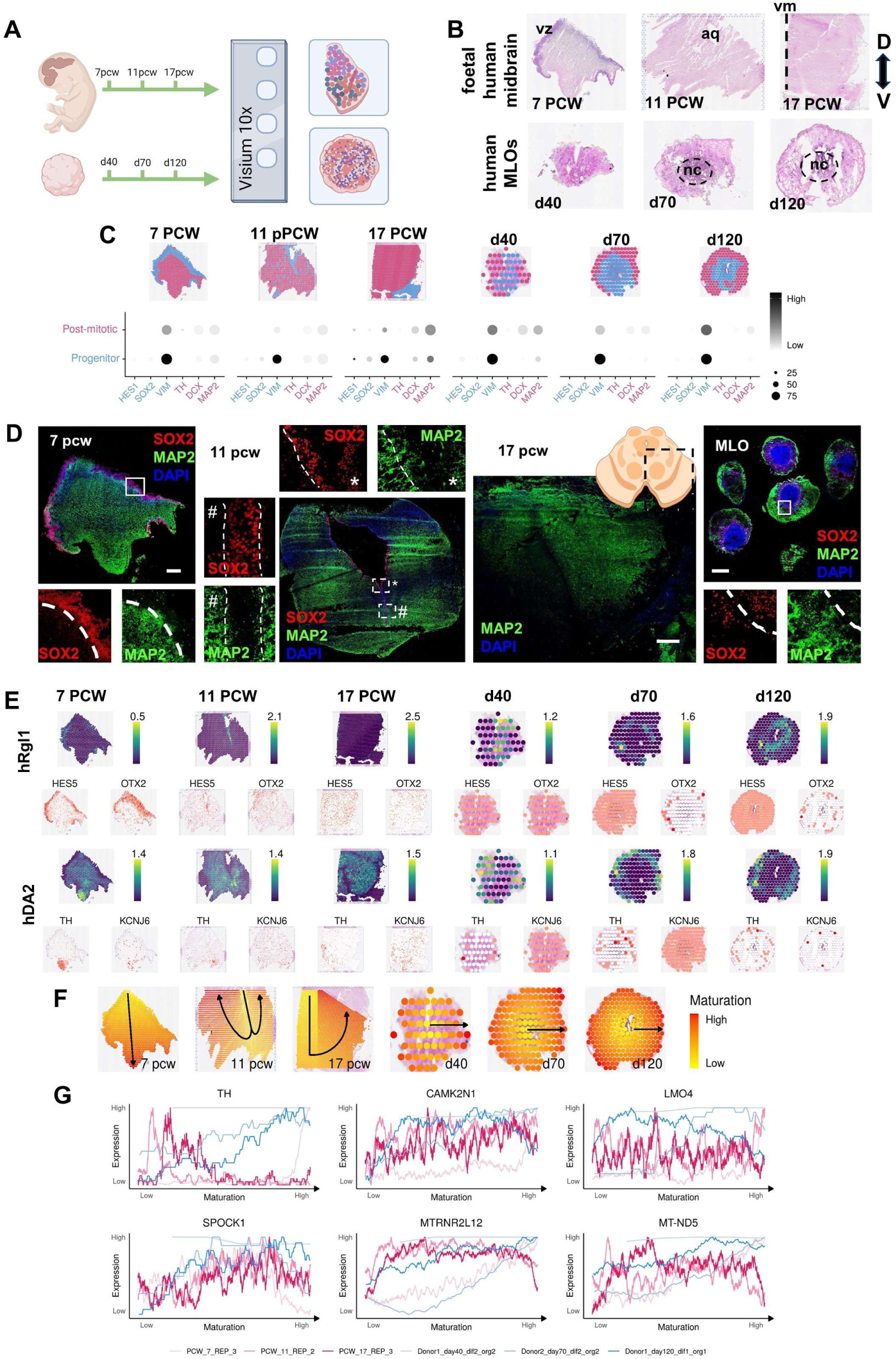
Spatial distribution of midbrain cell types across development. **A**, experimental design for the spatial transcriptomics. **B**, haematoxylin and eosin (H&E) stain of human fetal samples at 7, 11 and 17 PCW and MLO samples at 40, 70 and 120 days of differentiation. Ventricular zone (VZ), cerebral aqueduct (aq), dorsal (D), ventral (V). **C**, progenitor and post-mitotic cell populations in tissues and organoids, defined by clusters annotated according to anatomical features and marker genes (displayed below). **D**, immunofluorescence analysis of fetal and MLO samples for SOX2 and MAP2. Nuclei are stained with DAPI. Scale bar = 500 µm. **E**, Abundance of hRgl1 and hDA2 cells in tissues and organoids, as estimated by the Cell2Location deconvolution algorithm. Expression of two marker genes for each cell type are displayed below (white = low, red = high). **F**, manually annotated maturation paths on tissues and organoids. **G**, moving averages of the expression of genes that are differentially expressed with respect to maturation paths, calculated over a window of 50 spots along the path of maturation.

**Fig. 6.**
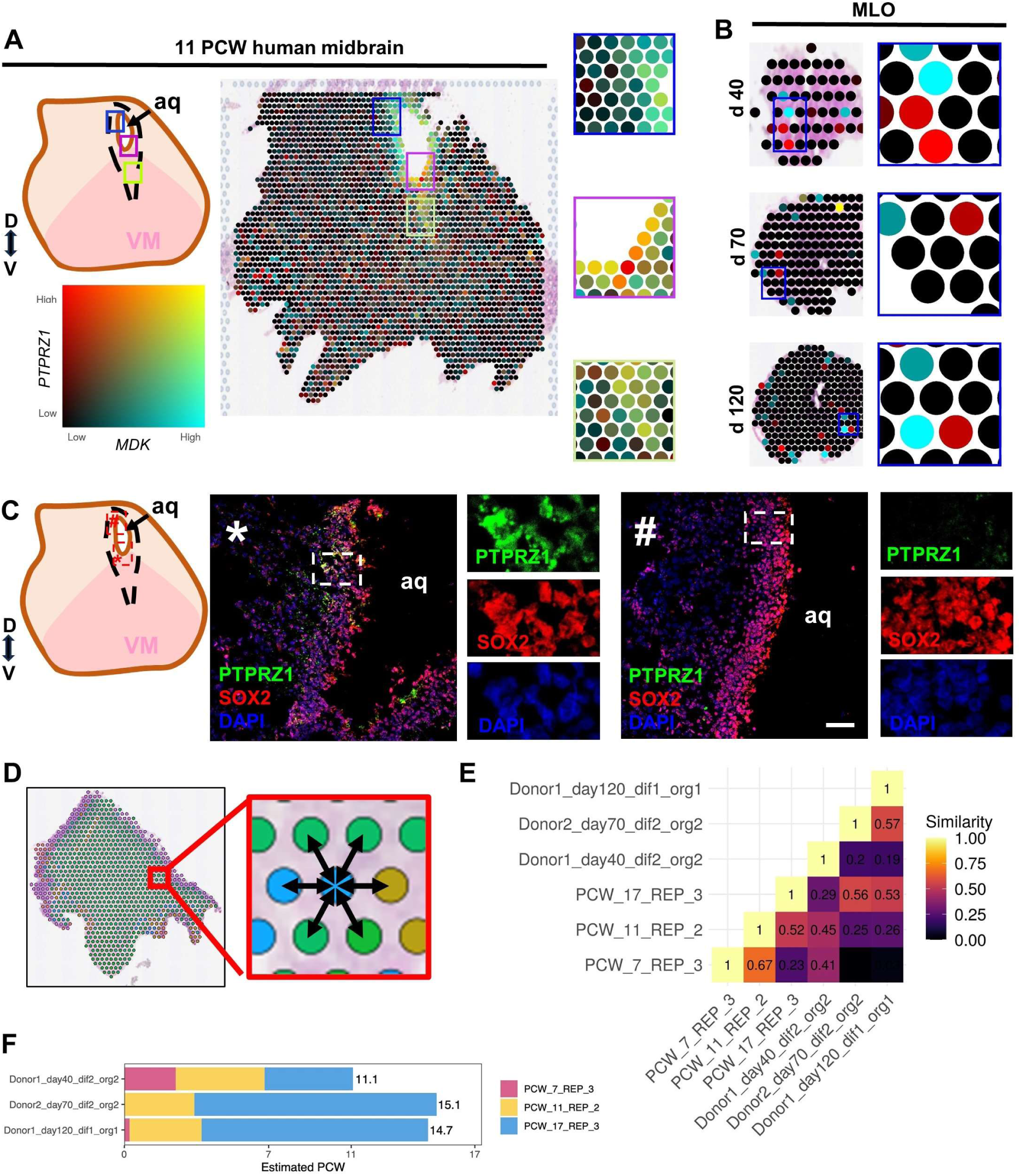
Fetal and MLO midbrain systems share spatial cell-cell communication pathways and spatial characteristics. **A**, Expression in 11 PCW tissue of the PTN-PTPRZ1 ligand-receptor pair, identified by COMMOT to be highly active and shown by Moran’s I to be spatially correlated. **B**, Expression in organoids of the PTN-PTPRZ1 ligand receptor pair. **C**, immunofluorescence analysis of 11 PCW human fetal coronal section for PTPRZ1 and SOX2. Nuclei are stained with DAPI. Scale bar = 50 µm. Cerebral aqueduct (aq). **D**, Schematic of similarity quantification approach: cell types of neighboring spots are counted and used to compute similarity between samples. **E**, Quantification of spatial similarity between tissues and organoids. **F**, Temporal alignment of organoids and tissues, where the timepoints of the tissues are averaged and weighted according to the organoid’s spatial similarity to each one.

### Single cell atlas of the developing human midbrain

To further understand both the cellular and molecular complexity of the developing human midbrain, we performed droplet-based scRNA-seq of the human fetal midbrain covering first-trimester (10 and 12 PCW) and second-trimester gestational ages (16 and 20 PCW), jointly with iPSC-derived 2D mDA neurons (days 40 and 70) and MLOs (days 40, 70, and 120). Overall, we profiled a total of 34,191 cells from six donors that were pooled in 13 samples (10X libraries prepared from a single channel) (**Fig 3A; Table S2**). Donor identity per cell was deconvoluted using demuxlet (Kang et al. 2018), samples were merged and normalized with Seurat (Hao et al. 2024) and integrated with Harmony (Korsunsky et al. 2019). We identified 24 cell types based on established markers (La Manno et al. 2016, Fiorenzano et al. 2021) (**Fig 3B-D; Supp Fig 7A,B; Table S3**). Four of the cell types corresponded to either vascular identities, including endothelial (hEndo), pericytes (hPeric)(La Manno et al. 2016), erythrocytes (Eryth)(Billett 1990), and vascular leptomeningeal (vascLepto)(Fiorenzano et al. 2021); or to glial cell types, including microglia (hMgl), astrocytes (Astro) and two oligodendrocyte progenitor cells (OPC1 and OPC2). We distinguished OPC1 and OPC2 populations by the higher expression of *SOX10* in the OPC1 cluster (**Supp Fig 7A**). We also identified four radial glia-like clusters (hRgl1, hRgl2/immAstro, hRgl3caudal, hRgl4/multiEpend) based on their specific transcriptomic signature: hRgl1 showed midbrain floor plate identity, expressing *FOXA2*, *CORIN*, *SOX6*, *PBX1*. Interestingly, hRgl4/multiEpend cells upregulated the expression of *CCNO* (Funk et al. 2015) related to cilium organization/assembly and axoneme assembly (ciliary core). We further identified three progenitor clusters: progenitor medial floorplate (hProgFPM), expressing *FOXA2*, progenitor midline (hProgM) and neuronal progenitors (hNPro); and two precursor clusters: midbrain precursors (hMidPre) and dopaminergic precursors (hPreDA). Two populations of neuroblasts were also identified, either with dopaminergic commitment hNbDA, expressing *TH, LMO3*, *DDC, CALB1* and *PBX1;* or with GABergic commitment (hNbGABA). Our clustering also segregated mDA neurons into four subpopulations: hDA1a, hDA1b, hDA2 and hDA3/hGABA/h. hDA1a and hDA1b were annotated based on their relative expressions (low in hDA1a, high in hDA1b) of *TH*, *KCNJ6* and *DCX* genes, while hDA2 expressed *TH*, *KCNJ6*, *CALB1* and *DDC*. The hDA3/hGABA/hSert was a mixed cluster population, expressing increased levels of *LMO3, PBX1* and *KCNJ6,* jointly with high levels of *GAD1*, *GAD2* and *MEIS2* (GABAergic-related genes) and *GATA3* (involved in serotonergic neurons differentiation). Finally, we identified a cluster of cells of unknown origin, which did not categorize in any of the previous cell identities. This cluster is abundant in genes related to collagens (*COL2A1*, *COL9A3*, *COL9A1*, *COL9A2*, *COL11A1*) which may represent connective tissue cells (Ricard-Blum 2011).

**Fig 7.**
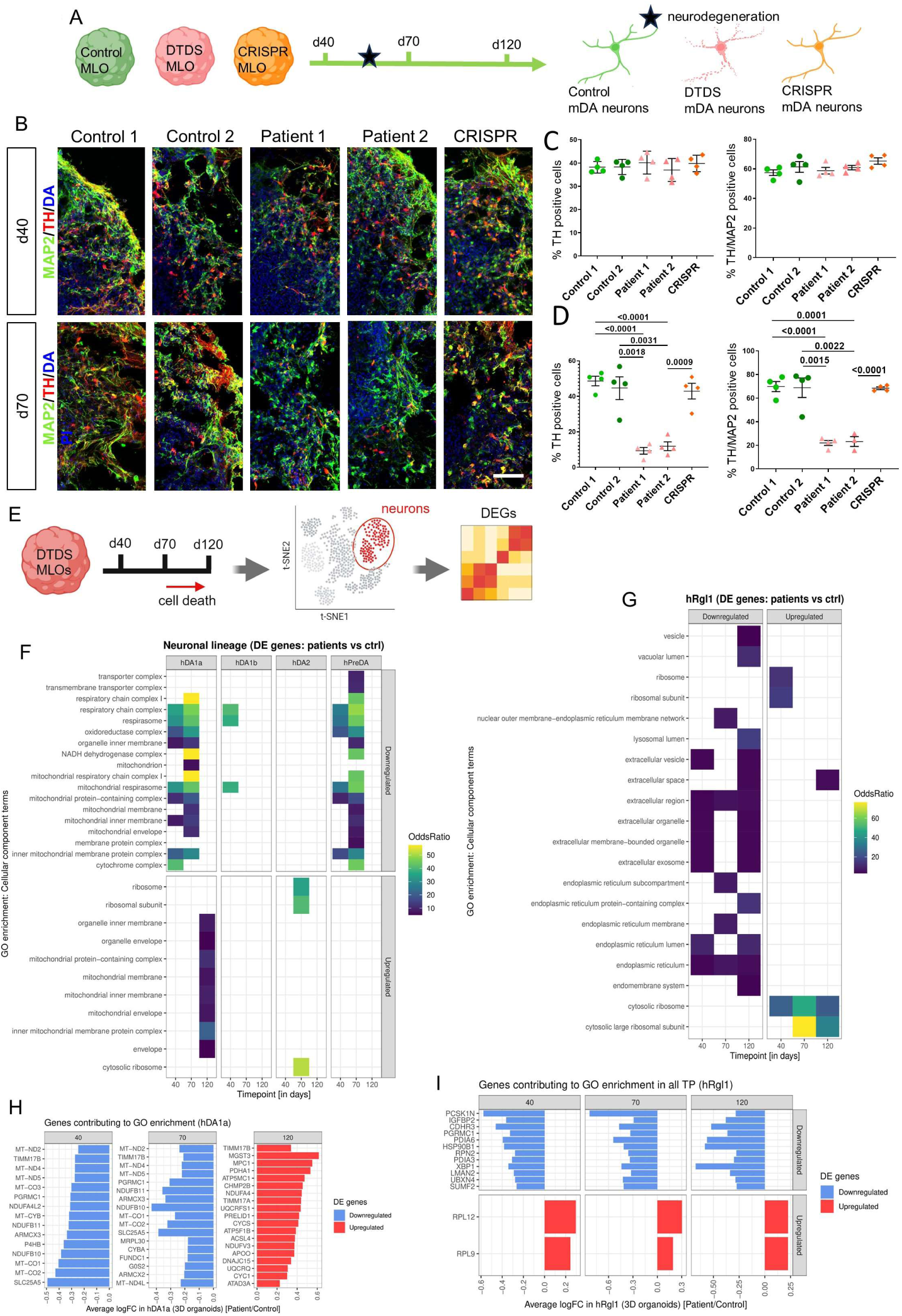
MLO derived from DTDS iPSC allow investigation of disease progression in dopaminergic neurons. **A**, experimental design for the analysis of DTDS derived MLOs at different maturation time points (40, 70, 120 days of differentiation) to investigate progressive neurodegeneration. **B**, immunofluorescence analysis for the neuronal markers TH and MAP2 at d40 and d70 of differentiation in Control 1, Control 2, Patient 1, Patient 2 and CRISPR isogenic-derived MLOs. Nuclei were counterstained with DAPI. Scale bar= 50µm. **C-D**, quantification of TH positive and TH/MAP2 double positive neurons for controls and patient lines (n=4 for every line). **E**, experimental design for the differentially expressed genes (DEGs) analysis in mDA clusters. **F-G**, Gene ontology enrichment analysis (GO) on cell compartments based on the DEGs in neuronal lineage and hRgl1 clusters respectively, in patients compared to controls. **H-I**, List of genes contributing to GO enrichment in hDA1 and hRgl1 clusters respectively. Error bars indicate SEM. DTDS lines were independently compared to controls using two-tailed Student’s *t*-test for all analyses. Statistically significant differences indicate comparison of Control 1 or 2 to Patient 1 or 2 and Patient 2 to CRISPR.

After cell type annotation, we estimated cell type proportions for each model and time point, with the most accurate estimates coming from first-trimester fetal samples, which had at least twice as many cells (**Fig 3D**). When comparing in vivo with in vitro models, we detected a larger proportion of cells with dopaminergic fate in vitro, either neuroblasts or neurons, due to the more directed differentiation towards the midbrain lineage. Conversely, microglia and radial glia with immature astrocyte identity (hRgl2/immAstro) were more abundant in fetal samples. Since our cells followed a continuous trajectory rather than clustering discretely, we used the MiloR framework (Dann et al., 2022) to test for differential abundance of cell types between the three models (**Fig 3E-G**).

By comparing 3D models against 2D ones, and considering the differentiation day as a covariate, we observed an enrichment of the dopaminergic population hDA1a and some of the precursors (hMidPre, hPreDA) in MLO (**Fig 3F**). On the other hand, some of the other dopaminergic neurons (hDA1b, hDA2) had subsets enriched in both models, indicating substantial cell state differences. Despite this, when running differential gene expression between 3D and 2D in the three neuronal populations, they all converged in the upregulation of the mitochondrial respirasome (Pval<0.05, FC>1.75), suggesting a more prominent role of mitochondria metabolism in the dopaminergic neurons from MLO (**Fig 3H, Table S4**). We then compared fetal samples with 3D MLOs, observing a strong enrichment of the mixed hDA3 population and a depletion of the other neuronal subtypes during in vivo development (**Fig 3G**). Differential gene expression analysis between fetal samples and MLOs revealed an upregulation of synaptic-related pathways in hDA2 fetal cells (synaptic membrane adhesion, presynaptic organization) (odds ratio, OR>10) (**Fig 3I; Supp Fig 7C, Table S5**). Conversely, hDA2 fetal cells showed downregulation of transcription and translation-related pathways, usually involving the Golgi vesicle-transport (**Supp Fig 7D**). As previously observed, fetal samples exhibited a higher abundance of microglia, radial glia and oligodendrocytes progenitors, highlighting greater differences between fetal samples and 3D organoids than differences between the two in vitro models (3D vs 2D). We also tested for differential abundance between first and second-trimester gestational ages and found microglia enriched in late fetal, while neuroblasts were enriched in early fetal stages (**Supp Fig 7E**).

To further investigate the transcriptomic differences observed between fetal and in vitro systems, we used immunofluorescence to analyze the spatial expression of tyrosine hydroxylase (TH) and the G-protein-regulated inward-rectifier potassium channel 2 (GIRK2), pivotal for synaptic maturation in mDA neurons of the VTA and SN (Reyes et al. 2012). Comparing SN samples at 19 and 22 PCW, we observed GIRK2 primarily in the soma of TH+ neurons at 19 PCW, shifting to cytoplasmatic and dendritic localization at 22 PCW (**Fig 3J**). In our 2D in vitro system, GIRK2 expression remained confined to the cellular soma, whereas in d120 MLOs, it localized to neurites (**Supp Fig 7F**), closely resembling the expression pattern seen in the 22 PCW fetal sample. These findings indicate that while there are differences in cell type composition and transcriptomic profiles between in vivo and in vitro systems, there are also similarities between the 3D and fetal samples that suggest comparable degree of maturity between these models.

### Transcriptional dynamics of integrated atlases of developing human midbrain in vivo and in vitro

To comprehensively assess the strengths of our scRNA-seq results, we integrated our dataset with previously published ones covering fetal development (La Manno et al. 2016, Birtele et al. 2022, Braun et al. 2023), in vitro dopaminergic models (Fiorenzano et al. 2021, Zagare et al. 2022) and adult post-mortem samples (Agarwal et al. 2020). In total, we integrated 179,428 cells using the mutual nearest neighbours implementation (fastMNN) from Seurat v5. The integration showed alignment of cells upon their origin (2D, 3D MLO, fetal), except for the Zagare dataset (Zagare et al. 2022) and, as expected, for the post-mortem adult brains (Agarwal et al. 2020) (**Fig 4A**).

To compare cell type composition differences, we established a unified cell type annotation based on the published annotations of the datasets (**Fig 4B-D, Table S6**). Neuron abundance varied significantly between datasets, with more stable proportions in vitro (min-max: 34.1%-39% in 2D; 6.1%-38.9% in 3D) compared to fetal samples (min-max: 1.2%-55.3%). Generally, neuron proportions decreased at later stages of fetal development, with a simultaneous increase in glial cells, microglia and oligodendrocyte cell lineages. Unsurprisingly, the proportion of neurons in adult post-mortem sample was only 3.3%. Fetal datasets exhibited large variability in cell type composition which became more pronounced at later gestational stages, potentially due to a combination of both biological and technical factors. Specifically, we found an increase in radial glia and a decrease in progenitors in only this and La Manno studies (La Manno et al. 2016), but not in the Birtele or Braun studies (Birtele et al. 2022, Braun et al. 2023) (**Fig 4B-D**).

We identified a developmental temporal axis among samples that correlates with the first component of the UMAP projection (**Fig 4E**). In this sequence, in vitro models precede first-trimester fetal samples, followed by second-trimester samples. Thus, cells from the oldest sample (20 PCW) are positioned at the right end of this developmental axis. Based on this observation, we compared the transcriptional alignment of iPSC-derived models to fetal samples using pseudobulk batch-corrected values. Our findings reveal that iPSC-derived models begin to diverge from fetal development around 10 PCW, consistently in this and the Fiorenzano studies (Fiorenzano et al. 2021) (**Fig 4F**). Based on transcriptional similarity, days 15-70 in MLO strongly correlated with gestational weeks (GW6-GW8) fetuses from La Manno (La Manno et al. 2016) (average R=0.68, Pearson’s correlation), while days 90-120 in MLO correlated with 12 PCW fetus from this study (average R=0.47). In contrast, one group has reported organoids (days 35-70) that exhibit similarities with adult post-mortem brains (Zagare et al. 2022), a feature not shared by any of the other considered organoid models.

Given the presence of this potential developmental axis, we inferred a pseudotime trajectory on the integrated dataset using three methods: Slingshot, Destiny and Monocle3 (Trapnell et al. 2014, Angerer et al. 2016, Street et al. 2018) (**Table S7)**. Among these, Destiny and Monocle3 showed a significant alignment on the final trajectory (R^2^=0.737). Consequently, we chose the Monocle3 output for downstream analysis, with the root for the start of the trajectory at floor plate progenitors (**Supp Fig 8A-B**). The trajectory followed the expected progression of cell type identities along the developmental axis, moving from radial glia, progenitors to precursors, intermediate progenitor cells, neuroblasts, and neurons. The sampling timepoint also influenced the pseudotime inference, even within the same cell types (**Fig 4G**). For instance, neurons in Braun samples (Braun et al. 2023), were, on average, associated with a larger pseudotime at PCW14 compared to PCW8. Likewise, progenitors at PCW10 appeared older than those from d120 3D organoids. Interestingly, this progression was not maintained when comparing neurons from these two samples (**Supp Fig 8C-D**). To analyze the neuronal developmental axis specifically, we compared the pseudotime distribution of neurons across the different datasets and timepoints. In each dataset, older samples exhibited a skewed distribution toward larger predicted pseudotime values, both in differentiation and in development (**Fig 4H**). This pattern was observed in 3D-organoids from day 120 and in late fetal samples (i.e. PCW20 in this study and PCW14 in Braun dataset). Among dopaminergic neuron subtypes from our study, the pseudotime values again showed some association with developmental time, especially for the hDA1b subtype in iPSC-derived models (**Fig 4I**).

To further understand the transcriptional dynamics of the dopaminergic system, we identified 50 gene modules of co-regulated genes whose expression changed with pseudotime (**Table S8**). We then computed activation scores for each cell to identify modules specific to cell type identities (**Supp Fig 8E**). While not all modules were cell type-specific, some showed high activation in certain cell types, such as module 1 in ODC and module 38 in IPC. We focused on the five most highly activated modules for neurons (modules 6, 12, 18, 21 and 46), which were also activated in neuroblasts and other cell types (**Supp Fig 8F**). For instance, module 12 was highly expressed in progenitors, while module 18 was highly expressed in precursors. To identify the biological processes behind these neuron-enriched gene modules, we performed gene ontology enrichment analysis with *gProfiler2 (Kolberg et al. 2023)*. We found significant enrichment in cellular components (p<0.05) for three of the modules: synapse in module 6, voltage-gated calcium channel complex in module 12, and axon terminus in module 18 (**Supp Fig 8G**).

The neuron-enriched modules showed activation differences across fetal cells from different publications (**Fig 4J-K; Supp Fig 8H**). In our study, all modules were significantly upregulated when compared to other datasets, except for module 12, which showed no difference with the Birtele study (Birtele et al. 2022) and was downregulated when compared to La Manno samples (La Manno et al. 2016) (*Fisher-Pitman permutation test*). This overall upregulation in our study can be partially attributed to the inclusion of second-trimester fetal samples. In line with this, module 12 is upregulated with developmental time, underscoring the relevance of calcium metabolism in neuron maturation (**Fig 4I**). Conversely, module 46 is downregulated with developmental time, explaining the lower activation scores in our fetal cells compared to the Braun dataset (Braun et al. 2023) (**Supp Fig 8K**). Comparing module activation scores within our dataset between in vitro models and fetal cells (**Supp Fig 8I**), we observed higher activation in vitro in those modules whose scores anticorrelate with developmental time (modules 18 and 46), and lower activation for those that actually correlate (modules 6 and 12). Using the 3D organoid cells from Fiorenzano (Fiorenzano et al. 2021) as a control comparison, we observed higher activation of the five neuron-enriched modules in our in vitro models (**Supp Fig 8J**), particularly in module 18, highlighting a more prominent role of axon terminus, necessary to release neurotransmitters. However, those differences are limited by the low number of neurons at late fetal stages resulting in more uncertain activation scores, as well using uncorrected (log-normalised) expression values to include all available genes. Overall, our results highlight consistent developmental patterns of in vitro models across studies, showing that they most closely resemble those of the first gestational trimester.

### Spatial cellular complexity of the developing in human midbrain and in vitro models

Whilst we have identified significant changes in the transcriptional dynamics of dopaminergic neurons differentiation at single cell level, incorporating spatial resolution is necessary to better understand the role of dopaminergic subtypes during human midbrain development. To decode the spatiotemporal developmental events of the ventral midbrain, we performed spatial transcriptomic analysis using the 10X Genomics Visium platform. We collected 7 PCW sagittal sections, along with 11 and 17 PCW coronal sections of the ventral human midbrain (n=3 donors) (**Fig 5A,B; Supp Fig 9A,B; Table S9**). Additionally, we analyzed MLOs at 40, 70 and 120 days of differentiation in groups of six per capture area on the Visium slide (**Fig 5A-B; Supp Fig 9C**). In fetal midbrain sections, we observed distinct spatial distribution of ‘progenitor’ cells in the VZ, expressing *OTX2*, *SOX2* and *VIM* (the area around the cerebral aqueduct was excluded in the 17 PCW midbrain sectioning due to the 6.5mm x 6.5mm constraint of the capture area). More mature (‘post-mitotic’) cells populate the IZ and MZ of the ventral midbrain and express *TH*, *DCX* and *MAPT* (**Fig 5C**). Similar spatial segregation was observed in MLOs at all differentiation time points, with the progenitors localizing in the more internal areas of the organoids. Immunofluorescence analysis confirmed differential expression of SOX2 and MAP2 in progenitors and post-mitotic cells respectively in both fetal midbrain and MLOs (**Fig 5D**).

To better investigate the spatial distribution of cell types in the developing midbrain and MLOs, we performed cell type deconvolution using our scRNA-seq dataset. We observed agreement between deconvoluted cell types and anatomical regions, with the scRNA-seq-defined cell types distributed along the dorso-ventral axis of the developing human midbrain (**Fig 5E; Supp Fig 10; Supp Fig 11A,B**). Eight immature clusters were in proximity of the VZ and cerebral aqueduct both at 7 and 11 PCW respectively. We identified all hRgl clusters (1-4) and progenitor clusters (hProgFPM, hMidPre, hNPro, hProgM) in the VZ. Neuronal precursors (hPreDA, hNbDA, hNbGABA) localized in the IZ and mantel zone (MZ) of the midbrain at 7 PCW. At later stages of development (11 and 17 PCW) hPreDA were mostly localized at the ventral midline of the midbrain in proximity to the VZ. A more diffuse distribution was observed for hNbDA and hNbGABA, with the latter mostly localizing in the more dorsal section of the developing midbrain at 11 PCW. hDA1a neurons were present in similar areas to hNbDA, although at distinct spots. hDA1b and hDA2 showed different spatiotemporal localizations during development. While at 7 PCW both populations were in the most ventral area of the midbrain where mDA neurons are localized, at 11 and 17 PCW a more scattered distribution was observed, with cells distributed at the ventral midline and the alar section of the tissue.

Ontogeny of mDA neurons is defined by the spatial localization of the neurons in the tissue and follows specific migratory stages. To further investigate developmental cellular trajectories across the tissue and MLOs, we identified genes that are differentially expressed along the path of maturation. For each sample, we manually annotated the maturation path and then used it as a covariate in the “cell type-specific inference of differential expression” (C-SIDE) algorithm, which controls for the influence of different cell types at different spots (i.e. differentially expressed genes are identified for each cell type) (Cable et al. 2022). We identified a total of 83 and 52 genes as differentially expressed along the paths of maturation in fetal midbrain tissue and organoids, respectively (**Table S10**). For all samples we then plotted the moving average expression along the maturation path of genes reported to be involved in mDA neurons maturation (**Fig 5F**). Similar changes in the average expressions of these genes were mostly observed between 7 PCW fetal samples and day 70 MLOs (**Fig 5G**). We detected increased expression along the maturation pathways of the dopaminergic neurons related genes *TH*, *CAMK2N1* and *LMO4* (La Manno et al. 2016, Fiorenzano et al. 2021, Zagare et al. 2022). Similar expression was observed across all samples for the extracellular-matrix-related gene *SPOCK1*, coding for SPARC/osteonectin, cwcv and kazal-like domains proteoglycan 1 (SPOCK1) protein, which plays a role in brain development and has been indicated to been specifically involved in murine mDA neurons specifications (Roll et al. 2006, Vadasz et al. 2007). The mitochondrial-related genes *MT-ND5* and *MTRNR2L12* - reported to be involved in neuronal energy regulation (Maffezzini et al. 2020, Traxler et al. 2021, Novak et al. 2022) and neuronal survival respectively (Elkjaer et al. 2019) - exhibited increased expression patterns over maturation both in the 7 PCW tissue and in the day 70 and 120 MLOs.

We further investigated ligand-receptor interactions in neighboring spots and highlighted communication pathways that are highly active during ventral midbrain development and investigated if they are maintained in MLOs. We used COMMOT (Cang et al. 2023) to compute the expression of pathways at each spot, and then used the Moran’s I non-parametric test for spatial autocorrelation to identify those with a large spatial component to their expression. As a result, we identified highly active communication pathways at 11 PCW fetal midbrain (**Fig 6A; Supp Fig 12**). Among the 20 most highly activated ligand-receptor pathways in this tissue, we identified an important role for heparin binding growth factors, involved in neuronal maturation. In 11 PCW fetal midbrain, the ligands, *PNT* (Jung et al. 2004, Mourlevat et al. 2005) and *MDK* (Bloch et al. 1992, Winkler and Yao 2014), showed expression in the VZ and IZ, throughout the ventral medial axes and the dorsal tectum. Both growth factors have a common receptor, the protein tyrosine phosphatase receptor type Z1 (*PTPRZ1*) (Fujikawa et al. 2019), which was expressed along the progenitor area and ventral medial line. We did not detect *PTPRZ1* in the dorsal tectum of the midbrain (**Fig 6A**). A small number of neighboring ligand-receptors pairs were identified in MLOs (**Fig 6B**). We then confirmed expression of PTPRZ1 in the ventral area of the midbrain, whereas it was absent in the dorsal area of the midbrain (**Fig 6C**). This highlights PTPRZ1’s role in the specification of ventral progenitor development and fate decision. Strong co-localization was also found for *MDK* and another of its receptors *LPR1* (Faissner 2023), mostly expressed in both the ventral and lateral IZ of the developing midbrain at 11 PCW. High expression of *MDK/LPR1* ligand-receptor pair was also observed in the MLOs across differentiation time points (**Supp Fig 12**).

Finally, we used the resolution provided by spatial transcriptomics to quantify the spatial similarity between fetal tissue and organoid samples to further elucidate temporal dynamics. First, we labelled each spot according to the cell type that was estimated in the deconvolution to be most abundant (**Fig 6D**). Then, for each sample, we quantified the spatial relationships between these labelled cell types (**Fig 6E**). We found the organoids to share high spatial similarities that align along maturation. The d40 organoid displayed some similarity to the 7 and 11 PCW tissues, and reduced similarity to the 17 PCW. The d70 and d120 organoids showed very low similarity to 7 PCW, low similarity to 11 PCW, and medium similarity to 17 PCW. The d40 organoid showed low similarity to d70 and d120, whilst these later timepoints exhibited medium similarity to one another. The tissues showed decreasing similarity as the differences in their timepoints increased. We estimated the temporal alignment of organoids and tissue timepoints and found that the d40 organoid corresponded to a tissue timepoint of slightly over 11 PCW whilst the d70 and d120 organoids both corresponded to a tissue timepoint of approximately 15 PCW (**Fig 6F**). Taken together, these data highlight the parallel developments of similar spatial distributions of developing midbrain dopaminergic neuronal populations in fetal and MLOs across different time points.

### A human-derived mDA neuronal in vitro model allows temporal and cell-type specific analysis of dynamics in neurological disease

Our previous observations provide further evidence of temporal variances in MLO and fetal midbrain development. This poses the question of whether the relative maturity of MLOs may pose potential limitations in their use for disease modeling. To address this paradigm, we investigated dopaminergic-specific temporal disease progression in an MLO model of the neurodegenerative disease, Dopamine Transporter Deficiency Syndrome (DTDS). DTDS is due to biallelic mutations in the *SLC6A3* gene, leading to impaired dopamine transporter (DAT) activity (Kurian et al. 2011). Using a 2D patient-derived mDA culture system, we have previously shown that DTDS is characterized by apoptotic neurodegeneration associated with TNFα-mediated inflammation, and dopamine toxicity (Ng et al. 2021). We differentiated our previously characterized DTDS patient iPSC lines together with two age-matched controls lines and isogenic CRISPR-corrected line (Ng et al. 2021) into MLOs (n=4 donors) (**Fig 7A**). After 20 days of differentiation, both the control and patient lines showed expression of midbrain precursors genes and proteins (**Supp Fig 13A,B**). Derived MLOs were further matured over 40, 70 and 120 days in culture, and showed gene and protein expression profile of mature mDA neurons (**Supp Fig 13A; Supp Fig 14-16**). We detected a GABA/TH double positive neuronal population, as similarly observed in the human fetal midbrain tissue. We could also detect GFAP positive cells at later stage of maturation (120 days of differentiation).

Temporal analysis of DTDS patient-derived MLOs (day 40, 70, 120) revealed a significant decrease in total neuron count in patient-derived MLOs, compared to both control and isogenic control lines (**Fig 7B-D**) after 70 days in culture. This neuronal loss was particularly evident in TH+ neurons, at both 70 and 120 days of differentiation (**Supp Fig 17A-C**). No neuronal loss was observed in both control and patient-derived MLOs at day 40. We also observed reduced total TH protein in DTDS MLOs at 70 days of differentiation despite expression of DAT, as previously reported in 2D mDA neuronal cultures (**Supp Fig 17D,E**) (Ng et al. 2021). To elucidate the mechanisms associated with dopaminergic neuronal loss in patient MLOs, we analyzed apoptotic processes via cleaved Caspase3 (c-CASP3) immunofluorescence. We observed increased c-CASP3 positive cells in the patient MLOs and specifically in TH-positive neurons, confirming our previous study in 2D mDA patient-derived model (**Supp Fig 18A,B**) (Ng et al., 2021). Notably, this analysis excluded the necrotic core of MLOs.

Using our single cell dataset, we computed the differentially expressed genes (DEGs) between control and patient lines within each cell population identified in the MLOs, using Seurat (**Fig 7E, Table S11**). Our analysis focused into the dopaminergic neuron lineage (hDA1a, hDA1b, hDA2) and glial cells (Rgl1). Gene enrichment analysis showed a downregulation of the cellular components related to the mitochondrial membrane and mitochondrial respirasome in hDA1a and hPreDA populations from patient MLOs at both 40 and 70 days of differentiation (**Fig 7F; Suppl Fig 18D**). Those components were mainly associated with the mitochondrial electron transfer chain complex I and IV for both hPreDA and hDA1a (*MT-ND2*, *MT-ND3*, *MT-ND5*, *MT-CO1*, *MT-CO2*, & *MT-CO3*) (**Fig 7H; Suppl Fig 18C,D**).

Interestingly, upregulated DEGs in hDA1a at day 120 are enriched in mitochondria components, and mainly associated to mitochondrial inner membrane (*MPC1, ATP5F1B, TIMM17B, TIMM17A, ATP5MC1*), cytochrome c (*UQCRFS1, CYC1*, *CYCS*) and cytoprotection (*PRELID1*), while ribosomal components are upregulated in hDA2 at day 70. (**Fig 7F-H**). In the hRgl1 population, downregulated DEGs were mainly associated with the extracellular space and endoplasmic reticulum across all MLO differentiation stages, while upregulated genes were linked to the ribosome (**Fig 7G-I**). Overall, these data show the temporal dynamics of DTDS disease progression, characterized by mitochondrial dysfunction specific to dopaminergic neurons prior to their neurodegeneration.

## Discussion

In this study, we provide a detailed molecular and cellular profiling of the first two trimesters of human fetal midbrain development from 6 to 22 PCW. Due to the rarity of these late-stage embryonic samples, our study offers the opportunity to provide both transcriptomics (single cell and spatial transcriptomes) and immunofluorescence images as a valuable resource for the research community. Our analysis, complemented by existing human datasets of midbrain development (La Manno et al. 2016, Birtele et al. 2022, Braun et al. 2023), reveals the structural and cellular complexity of the human midbrain, which emerges mostly during the second gestational trimester.

Indeed, during the second trimester of human fetal midbrain development, we observed the spatiotemporal emergence of major ventral midbrain nuclei, including the red nucleus (RN), substantia nigra (SN), and ventral tegmental area (VTA) (Caminero 2022), along with the appearance of markers indicative of cellular maturity. Cellular maturation of mDA neurons occurs throughout radial migration of neuronal precursors from the ventricular zone to the mantel zone and then a tangential migration towards the specific nuclei location (Almqvist et al. 1996, Nelander et al. 2009). Our observations suggest that while radial migration of mDA neurons has occurred by 13 PCW, significant tangential migration, neuronal morphological maturity, and neuronal network complexity develop after this stage, and mostly between 19 and 22 PCW. This reflects a late specification of dopaminergic nuclei similar to what is observed in mice, where the tangential migration and maturation of mDA neurons occur between embryonic days E13.5 and E18 (Bodea et al. 2014, La Manno et al. 2016, Tiklova et al. 2019, Vaswani et al. 2019). In agreement, histological analysis of second trimester fetal midbrain already reveals structural complexity similar to that of adult tissue (Johns 2014, Gupta 2017). In contrast, our transcriptomic analysis showed a discrepancy between second trimester and adult samples. It is likely that major changes in midbrain maturation happen later in human development which, given the limitations in acquiring human fetal samples, have not been captured in our analysis (Braun et al. 2023).

We also investigated the similarity of later stage human midbrain development to human pluripotent stem cell-derived midbrain dopaminergic neuronal models, including 3D models (Fiorenzano et al. 2021, Birtele et al. 2022, Zagare et al. 2022). To our knowledge, there is limited understanding as to whether human pluripotent stem cell-derived mDA models truly replicate cell type location, proportion, gene expression, cell-to-cell communication and differentiation along developmental time and space of the human midbrain. Therefore, understanding the extent of biological recapitulation in these models is essential to justify their use as biological proxies. We therefore profiled fetal samples, 2D mDA neuronal cultures and 3D MLOs samples and integrated them with other previously profiled fetal midbrain, post-mortem and MLO samples (La Manno et al. 2016, Agarwal et al. 2020, Fiorenzano et al. 2021, Birtele et al. 2022, Zagare et al. 2022) to create a comprehensive map of the midbrain throughout development. This allowed us to provide an in-depth single cell comparison between human fetal samples and in vitro models. Our analysis revealed that the transcriptomic profiles of late-stage MLOs (120 days) closely resembled those of 8-12 PCW fetal samples. This finding indicates that, although MLOs represent a more advanced culture system than 2D cultures, their transcriptomic profile aligns to a less mature human midbrain developmental stage. This discrepancy underscores the difficulty to fully recapitulate developmental progression in vitro, raising questions regarding the utility of organoids in modeling later-stage diseases such as juvenile-onset, adult-onset and sporadic forms of parkinsonism.

To further investigate this, we specifically examined the dopaminergic cell population. In particular, scRNA-seq comparisons between the hDA2 clusters in vivo and in vitro demonstrated higher expression of synaptic gene markers in fetal samples compared to in vitro models, suggesting that MLOs lack appropriate synaptic targets or sufficient maturity. Surprisingly, immunohistochemistry analysis showed GIRK2 (a potassium channel expressed in mature neurons) was uniquely localized to neurites in both late-stage MLOs (120 days) and late fetal samples (22 PCW). This suggests that although the transcriptomes of our MLOs align to earlier fetal stages, certain features of MLO neurons may achieve a level of maturity similar to later stages of fetal midbrain development.

We hypothesized that some discrepancies in maturity between fetal midbrain and MLOs may be attributed to the loss of spatial and cell-to-cell interactions inherent in single cell approaches, which decontextualize cell behavior. Spatial transcriptomics addresses this limitation by spatially aligning the RNA content of cells, enabling the comparison of in vitro models and tissue samples based on spatial organization and cellular interactions (Williams et al. 2022). Our spatial analysis uncovers developmental pathways in both in vitro and tissue samples, allowing us to identify conserved DEGs along these maturation pathways. Additionally, our spatial analyses revealed important cell-to-cell communication networks within the tissue. We identified active ligand-receptor pathways, notably involving heparin-binding growth factors PNT and MDK and their receptor PTPRZ1, conserved in both in vivo and in vitro systems. The spatial information provided by this approach allowed us to estimate the temporal alignment of our MLO system with fetal brain samples based on spatial characteristics. Notably, this revealed an increased similarity between the late-stage MLOs and later stages of fetal midbrain, aligning day 70 and 120 MLOs closer to fetal midbrain samples from the second trimester of gestation, both peaking at approximately 15 PCW in maturity. This improves our single cell analysis of MLO maturity by about 5 PCW, while highlighting the importance of spatial context when considering the in vitro recapitulation of developmental progress. Our description of later stage midbrain cell populations may enable future refinement of in vitro culture systems, better reflecting human dopaminergic and glial populations across late developmental stages, leading to improved modeling.

Although iPSC-derived models may not fully replicate the exact maturation stage of their in vivo counterparts, their usefulness may largely rely on the specific cell type and phenotype under study. Indeed, despite certain imperfections, human-derived in vitro models offer the possibility to investigate disease progression over time in a patient relevant context. We leveraged this by developing MLOs from DTDS patient-derived iPSCs and investigating neurodegenerative progression over time at molecular and single cell transcriptomic levels. Interestingly, we observed specific dopaminergic neurons loss, which was not detected in our previous study in 2D mDA cultures (Ng et al. 2021). This might reflect the ability of MLOs to better recapitulate the pathological cellular interactions and microenvironment of the midbrain in a diseased state (Quadrato et al. 2017, Tekin et al. 2018). Temporal scRNA-seq transcriptomic analysis in DTDS MLOs indicates mitochondrial dopaminergic-specific dysfunction at an early stage of maturation, before dopaminergic neuron loss occurs. Notably, DTDS MLOs at a late stage of maturation showed dysregulation of cytochrome-related genes and increased expression of *PRELID1*, which might indicate a progression of mitochondrial dysfunction and neurodegeneration over time (Green 2005, Gillen et al. 2017, Ng et al. 2021). Overall, our analysis suggests that DTDS primarily affects the dopaminergic neurons, at least in the early stages of the disease, with a process primarily impacting mitochondria. Understanding these neurodegenerative mechanisms, particularly the role of different cellular subtypes in the progression of DTDS, will aid in the development of precision disease-specific therapies, as well as broader spectrum agents that may more generically target neurodegeneration.

In conclusion, our analysis provides a detailed characterization of the main cell types and tissue organization across the first two trimesters of human midbrain development, including the first reported profile of second trimester midbrain. Moreover, this study importantly highlights key similarities and differences between the human fetus and human-derived midbrain models, revealing the limits of in vitro models at late stages. Ultimately, our description of human midbrain cellular, transcriptomic and spatial characteristics will enable further refinement of in vitro culture systems for improved modelling.

## Methods

### Human embryonic tissue collection

The human embryonic and fetal material were provided by the Joint MRC/Wellcome Trust (grant MR/R006237/1) Human Developmental Biology Resource (www.hdbr.org). Human fetal tissues were collected from 6 – 22 PCW from legally terminated embryos and gestational age of each embryonic and fetal was determined by either crown rum length (mm), foot length and/or knee-heel length (mm). Samples from HDBR Newcastle were shipped overnight on ice in Hibernate media (ThermoFisher Scientific), while samples from HDBR London were handed in DMEM medium and processed on the same day.

### Generation of iPSC-derived 2D mDA neurons and 3D MLO

2D mDA neurons and 3D MLOs were derived from iPSCs of two healthy individuals, two patient lines with missense mutations in *SLC6A3* (patient 1: c.1103T>A, p.L368Q; patient 2: c.1184C>T, p.P395L) and a CRISPR-Cas9 corrected isogenic line from patient 2 (Ng et al. 2021). IPSCs were differentiated into 2D mDA neuronal cultures as previously described (Ng et al. 2021, Rossignoli et al. 2021). For MLO generation, iPSCs at day 0 were dissociated to a single cell suspension, resuspended in embryoid body medium (EBM) and plated at a density of 10,000 cells/well in a 96 low-attachment U-shape wells plate (Corning). EBM consisted of 1:1 DMEM/F12:Neurobasal medium, 1:100 N2, 1:50 B27 w/o vitamin A supplement (Invitrogen), 1% GlutaMax, 1% minimum essential media-nonessential amino acid (MEMNEAA), 100 U/ml penicillin G and 100 μg/ml streptomycin (Gibco), and ROCK-inhibitor (Thiazovivin) for the first two days. EBM was supplemented with: 0.1% β-mercaptoethanol (Gibco), 1μg/ml heparin (Sigma-Aldrich), 10 μM SB431542 (Cambridge Bioscience), 100 nM LDN193189 (Generon), 0.8 µM CHIR99021 (Tocris Bioscience) and 100 ng/ml hSHH-C24-II (R&D Systems), and on day two, 0.5 μM purmorphamine (Cambridge Bioscience) was added. At day 7, MLOs were embedded in 10 ul Matrigel and SB431542 withdrawn from medium. The following day, embedded MLOs were transferred to a low attachment plate and cultured in final differentiation medium, consisting of Neurobasal medium, 1:100 N2 supplement, 1:50 B27 w/o vitamin A supplement, 1% GlutaMax, 1% MEMNEAA, 1% Pen/Strep, 10.1% β-mercaptoethanol (Gibco), 0.2 mM ascorbic acid (AA) (Sigma), 20 ng/ml BDNF (Miltenyi Biotech), 0.5 mM dibutyryl c-AMP (Sigma-Aldrich) and 20 ng/ml GDNF (Miltenyi Biotech), 200 ng/ml laminin (Sigma-Aldrich). From day 9 to day 14, medium was supplemented with 100 ng/mL FGF8 (Miltenyi Biotech), and with γ-secretase inhibitor DAPT (10 μM, Tocris) from day 30 until the end of differentiation. The organoids were cultured under static conditions and medium changed every other day.

### Immunostaining

Fetal midbrain samples were washed in ice-cold PBS, and the meninges removed using tweezers. Samples were fixed in ice-cold 4% paraformaldehyde (PFA) for 10 min (2D mDA neurons), 30 min (MLOs) or overnight (fetal midbrains), washed with PBS, and fetal and MLO samples immersed in 30% sucrose in PBS at 4°C until saturated. Fetal midbrain and MLO samples were embedded in cryomolds using optimal temperature cutting (OCT) compound and frozen at −80°C. 2D mDA neurons, cultured on LabTek slides, were stored in 1 X PBS at 4°C. All embedded samples were sectioned at 12 μm onto SuperFrost Plus microscope slides. All slides were washed with 1 X PBS and blocked using 5% bovine serum albumin (BSA), 0.3% Triton in 1 X PBS for 1h at room temperature. Primary antibodies (**Table S12**) were diluted in blocking solution and incubated overnight at 4°C, followed by 3 washes in 1 X PBS. Secondary antibodies were diluted in blocking solution and incubated 1h at RT, followed by 3 washes in 1 X PBS. All secondary antibodies used were AlexaFluor used at a dilution of 1:400.

### Reverse transcription and qPCR

Total RNA was extracted following RNeasy kit guidance (QIAGEN). RNA purification and cDNA preparation were performed according to manufacturers’ guidance (Invitrogen). Samples cDNA were mixed in a qRT-PCR plate with MESA BLUE qPCR 2X MasterMix Plus for SYBR Assay (Eurogentec) that contained the appropriate target primer mix (ratio: 1 µl primers/10 µl MasterMix) (FOXA2, LMX1A, LMX1B, EN1, EN2, OTX2, NURR1, TH, PITX3, DAT, SNCA and GAPDH, **Table S13**). Each target was plotted in triplicates for each biological sample. GAPDH was used as a housekeeping gene for normalization purposes. qRT-PCR was performed on the StepOnePlusTM Real-Time PCR System (Applied Biosystems) with the following protocol: initiation at 95°C for 5 min, 40 cycles (denaturation 95°C for 15 s, annealing/elongation at 60°C for 1 min$ elongation 72°C. Gene expression was analyzed using the ΔΔCT method, with an age-matched control mDA line as control.

### Single cell isolation and 10X sequencing

Fetal, 3D and 2D iPSC-derived samples were mechanically dissociated to generate single cell suspensions using the Papain Dissociation System (Worthington). Following the supplier’s recommendations, the EBSS was equilibrated with O_2_:CO_2_ to improve cell survival. Samples were weighted and minced into small pieces using open-wide tips coated in BSA 7.5%. After three ice-cold PBS washes, 200 uL of the dissociation mix (18.85U/mL Papain, 50ug/mL DNase I, 1:4 Accutase/Accumax, 4ug/ml Actinomycin D) for 30 min at 37°C on a thermoshaker (750 rpm). Samples were triturated 10 X with a narrow pipette tip every 10 min for a total of 40 minutes. Cells were then pelleted at 300g for 4 minutes and resuspended in the inhibitor mix (148.25uL EBSS, 50ug/mL DNase I, 17.5uL Ovomucoid for 175uL). 90 uL of Ovocumoid and 90 uL of EBSS were added on the top and gently mixed by tube flickering. Cells were pelleted at 300g for 4 minutes. After discarding the supernatant, the pellet was resuspended in 10X buffer and filtered (40um cell strainer, membrane humidified with 10X buffer). Alive cell count target was >90% (LUNA cell counter, acridine orange/propidium iodide stain). Barcode and library preparation was performed using the 10X Genomics Chromium platform (v.2) and following the manufacturer’s instructions. Briefly single cell suspensions were prepared and processed using the Chromium Single Cell 3’ Reagent Kits v3 (10x Genomics, CG000183 Rev A). Cells were encapsulated into Gel Bead-In-EMulsions (GEMs), followed by reverse transcription, cDNA amplification, and library construction. Samples with different genetic backgrounds were randomly pooled across 10x libraries (ie GEM wells), as indicated in **Table S14**. Nine libraries (2D and 3D iPSC-derived samples) were sequenced with Illumina-HTP HiSeq 4000 paired-end sequencing. Four libraries corresponding to the fetal samples were sequenced in a separate batch using Illumina-HTP NovaSeq 6000 paired-end, across 4 lanes. To discard any potential sequencing lane effect, each 10x library was sequenced in all 4 lanes. Specifically, each fetal library was further subdivided into four samples, with each sample sequenced in one of the 4 lanes of the NovaSeq600 paired-end sequencing.

### Data preprocessing (single-cell RNA-seq)

We processed the thirteen pooled 10x libraries (ie. GEM well) using CellRanger (version 3.1.0) (**Table S2**). Using the “*cellranger count*” pipeline and the sequencing data (FASTQ files) as input, we performed alignment to GRCh38 reference genome, barcode counting and UMI counting to generate filtered feature-barcode matrices for each library. The genes counts were quantified using the Ensembl 93 reference gene annotation (N=33,538 genes). For each 10x library, we used 32 CPU cores, 200 GB of memory and a maximum of 64 jobs to be run simultaneously. Only for fetal samples, the “*-fastqs”* argument was used to specify the paths to the multiple FASTQ files generated from the library subdivisions in the NovaSeq600 sequencing.

The matrices of each 10x library went through a quality control step based on the distribution of i) expressed genes per cell, and ii) the mitochondrial count fraction, so as to remove dying cells or those with broken membranes. For 2D and 3D-iPSC derived samples, cells expressing <1,500 or >6,000 genes were discarded, as well those with an excess of mitochondrial count fraction (>10%). For fetal samples, with a different distribution for both parameters, cells were removed when expressing <1,500 or >5,000 genes, and when the mitochondrial count exceeded 25%.

To remove doublets/multiplets in the single-cell libraries, we performed cell deconvolution with demuxlet (Kang et al. 2018), using existing genetic variation from the eight donors previously genotyped with SNP arrays (VCF file, **Data S1,S2**). Demuxlet was run using a default prior doublet rate of 0.5 and only those cells unambiguously linked to a donor (singletons) were retained. All pooled libraries presented an acceptable singleton rate (>60%), and all expected donors were deconvoluted.

### Normalisation, dimensional reduction, clustering and projection (single-cell RNA-seq)

After preprocessing, all 10x libraries were merged and all genes that were not expressed in at least 0.1% of the total cells were removed. The gene counts were initially normalized to a scale factor of 10,000 and log-transformed (log1p). We then selected the 2,000 highly variable genes (HVG) based on the variance stabilizing transformation and scaled the data using default Seurat parameters (scale.max=10). Based on this selection, we then calculated the first 50 principal components (PCs) and batch-corrected them with Harmony (v.0.1.0) (Korsunsky et al. 2019), treating each 10x library as a different batch (default parameters, tau=0, max.iter.harmony=10, max.iter.cluster=20). We used the batch-transformed PCs to compute a neighborhood graph (neighbors=10), visualize the graph using UMAP and performed cell clustering using the Louvain algorithm (resolution=0.75), identifying 24 different clusters (**Data S3**). We used the Seurat R package (v.4.1.3) (Hao et al. 2024) for all the processing steps.

### Cell type annotation (single-cell RNA-seq)

Cell type annotation was performed using a manually curated set of markers (**Table S3)** established in previous studies (La Manno et al. 2016, Fiorenzano et al. 2021). Initially, we iteratively performed differential expression (DE) for each unannotated Seurat cluster against all other clusters to generate a list of specific cell type markers. We identified cell types by intersecting the DE list with the curated markers, while also considering the expression level of each marker. We confidently identified 23 out of the 24 clusters, with one unknown cluster likely having a connective tissue identity. Based on those markers, we also calculated gene module scores per cell type to evaluate the specificity of the annotation using the “*AddModuleScore*()” function from the *Seurat* package (ie. Figure 3B).

### Differential abundance analysis (single-cell RNA-seq)

To assess cell type composition differences between models, we used the *miloR* package (v.1.2.0) (Dann et al. 2022) that performs differential abundance testing. This statistical framework assigns cells to neighborhoods based on a k-Nearest Neighbor (kNN) graph, being particularly well-suited for continuous trajectories observed in neurodevelopment or in differentiation.

We tested three different comparisons:

a. In vitro models: 2D vs 3D (accounting for differentiation day as a covariate)
b. In vitro 3D model vs in vivo fetal
c. Post-conceptional weeks of fetal gestation (early vs late foetal)

For each comparison, the kNN graph was built using the *buildGraph*() function, using 30 nearest-neighbors (k=30) and 30 Harmony batch-corrected dimensions (d=30). Then, we defined neighborhoods on a graph with *makeNhoods*() function using the same parameters (k=30, d=30) and randomly sampling 30% of the graph vertices. We only used 20% of the vertices in the fetal comparison. Cells were then counted for each neighborhood and experimental sample (for “a-b” comparisons, the experimental sample is defined by combining “donorDimensions” and “timePoint” metadata columns from **Data S3**; and for the “c” comparison, is defined by combining “midBrainId” and “timePoint” columns). After that, distances between neighborhoods were calculated using the *calcNhoodDistance*() function using again 30 Harmony dimensions (d=30). We performed differential neighborhood abundance testing with the following design for each comparison:

a. ∼ Time + originDimensions (discrete: 2D vs 3D)
b. ∼ originDimensions (discrete: 3D vs foetal)
c. ∼ timePoint (continuous: post-conceptional weeks 10, 12, 16, 20)

Finally, neighborhoods were annotated with a given cell type when the most frequent identity among the cells from each neighborhood was >70% (>75% in “c” comparison). Otherwise, non-homogenous neighborhoods were annotated as “Mixed”. Results were visualized with a custom beeswarm plot using default FDR significance level (<0.1). The log-fold change values of differentially abundant cell type-specific neighborhoods indicates the following enrichments between conditions:

a. Enriched in 3D (logFC>0), enriched in 2D (logFC<0)
b. Enriched in fetal (logFC>0), enriched in 3D (logFC<0)
c. Enriched in late fetal (logFC>0), enriched in early fetal (logFC<0)

### Differential expression analysis (single-cell RNA-seq)

We performed differential gene expression (DGE) analysis between different conditions and experimental contexts:

1. Comparison between 3D and 2D in vitro models within differentially abundant dopaminergic populations (hDA1a, hDA1b, hDA2). To do so, we first aggregated neighborhoods in groups based on their cell type identity. We then ran differential expression using the *testDiffExp*() function of the miloR package for each dopaminergic population using the input design (∼*Time + originDimensions)*. Differentially expressed genes were selected based on significance level (adjusted P-value<0.05) and fold-change (upregulated: FC>1.75; downregulated: FC<1/1.75).
2. Comparison between fetal hDA2 and 3D hDA2 dopaminergic population. We subsetted our integrated dataset by only considering hDA2 cells from this publication and then filtering out 2D samples. Then, we set the object’s identity class to the metadata column “system”, and we ran differential expression by using the *FindMarkers*() function from Seurat, being the two comparing classes “Foetal” and “Organoid”. Significance level was set at an adjusted P-value<0.05 and a fold-change difference on expression of 1.5.
3. Patient versus controls (2×2 donor comparison in 3D models): For each timepoint (days 40, 70 and 120) and cell type with a minimum of 10 cells for both conditions, we ran differential expression analysis using the *FindMarkers*() function from Seurat. Significance level was set at an adjusted P-value<0.05 and a fold-change difference on expression of either 1.5 (upregulation) or 1/1.5 (downregulation).

### Gene ontology enrichment analysis (single-cell RNA-seq)

Based on the differential expression analysis, we performed gene ontology enrichment analysis on the following results:

1. Comparison between 3D and 2D in vitro models within differentially abundant dopaminergic populations (hDA1a, hDA1b, hDA2). The gene universe was defined by the 22,032 genes from our single-cell data that passed QC on preprocessing. Then for each dopaminergic population, we run the functional enrichment on the list of DE genes using the *gost*() function from the *gprofiler2* R package (Kolberg et al. 2023). We set the significance threshold at an adjusted P-value<0.05 using the g:SCS method for multiple-test correction. We defined the gene universe in the “custom_bg” and used the functional annotation from the Cellular Component category of Gene Ontology (GO:CC) (Gene Ontology et al. 2023). Based on the term size, the difference between the effective domain size and the term size, the query size and the intersection size, we computed the odds ratio for each term. In Fig 3H, only the GO:CC enriched terms in dopaminergic populations hDA1a, hDA1b and hDA2 are shown.
2. Comparison between fetal hDA2 and 3D hDA2 dopaminergic population. The gene universe was defined by the 22,032 genes from our single-cell data that passed QC during preprocessing. Before the GO analysis, we annotated each gene with their corresponding Entrez gene identifiers using the package *org.Hs.eg.db* from R/Bioconductor. We filtered out those genes without correspondence or showing duplicate identifiers. We then run the hypergeometric test for GO term overrepresentation of biological processes (GO:BP) conditional to the hierarchical GO structure (package *GOstats* from R) (Falcon and Gentleman 2007). The test was run separately for upregulated and downregulated genes. In each case, we used the following thresholds for significance: minimum gene set size = 25, maximum gene set size=500, adjusted P-value<0.05, minimum number of counts per gene set = 10. In Fig 3I and Supp Fig 7C, only the top-5 upregulated biological processes and top-10 are shown, respectively. In Supp Fig 7D, only the top-10 downregulated biological processes are shown.
3. Patient versus control (3D models): We performed GO enrichment analysis on the list of differentially expressed genes for the three dopaminergic populations (hDA1a, hDA1b, hDA2), dopaminergic precursors (hPreDA), dopaminergic neuroblasts (hNbDA), as well midbrain precursors (hMidPre) and radial glia type 1 (hRgl1), separately for upregulated and downregulated genes. We used the same parameters to run the *gost*() function and to compute the odds ratio as detailed in the first comparison. Results are illustrated in Fig 7F-G. We then evaluated which genes contributed the most to the hDA1a enrichment on cellular components, either in downregulation (days 40/70) or in upregulation (days 120) (Fig 7H). Additionally, we checked which genes consistently contributed to cellular components that were enriched across all timepoints in hRgl1 (Fig 7I), either upregulated (cytosolic ribosome) or downregulated (endoplasmic reticulum, extracellular region). Finally, we also checked which genes contributed to the downregulation of the mitochondrial components at day 120 in midbrain precursors (hMidPre) (Supp Fig 18D).

### Integration analysis

We integrated our single-cell dataset with six published midbrain development datasets (total of 194,521 cells) that either contained cells from in vivo fetal development or from iPSC-derived in vitro models (**Data S4**). Here we shortly describe each dataset and where the data was obtained from:

- **This work** (2D, 3D, Fetal): It encompasses 49,284 cells, containing four developmental stages of fetal development (one donor sampled at post-conceptional weeks 10, 12, 16 and 20, respectively) and iPSC-derived 2D and 3D models. 2D models have been profiled at two differentiation timepoints (days 40 and 70; n=2 controls), and 3D models at three timepoints (days 40, 70, 120; n=2 controls, n=2 patients, n=1 patient-corrected line). After annotation, 24 cell types were identified, with 19% of the cells being dopaminergic neurons. To download our data, check the specific data availability section.
- **La Manno et al. 2017** (Fetal) (La Manno et al. 2016): It encompasses 1,977 cells sampled at six developmental stages of fetal development (gestational weeks 6, 7, 8, 9, 10 and 11). 26 cell types were identified in annotation, with 3 dopaminergic populations accounting for 6.2% of total cells. Raw counts were downloaded from the Gene Expression Omnibus (GEO) using the accession identifier GSE76381 at https://www.ncbi.nlm.nih.gov/geo/query/acc.cgi?acc=GSE76381. From the list of available files, we only processed the embryo molecule counts file.
- **Braun et al. 2023** (Fetal) (Braun et al. 2023): It encompasses 16,800 midbrain cells from 2 donors sampled at post-conceptional weeks 8 and 14, respectively. The selected cells are a subset from a larger atlas of the first trimester developing human brain. In the midbrain, 11 cell classes were identified, of which 41.4% of the cells were neurons. Raw counts were downloaded from the complete processed dataset “*HumanFetalBrainPool.h5*”, previously deposited in the *Github* repository: https://github.com/linnarsson-lab/developing-human-brain. We subset the original matrix of counts by selecting only “Ventral midbrain” cells from the metadata “Tissue” column, and by selecting only the valid genes, as defined by the authors.
- **Birtele et al. 2022** (Fetal) (Birtele et al. 2022): It encompasses 23,483 cells from four fetal ventral midbrain samples, sampled at three developmental timepoints: post-conceptional weeks 6 (MP06-hVM-6wks), 8 (MP03-hVM-8wks) and 11 (MP02-hVM-10-5wks and MP01-hVM-11-5wks, technical replicates). In those samples, 10 cell types were identified, of which 16.3% were dopaminergic neurons. Raw counts were downloaded from the Gene Expression Omnibus (GEO) using the accession identifier GSE192405 at https://www.ncbi.nlm.nih.gov/geo/query/acc.cgi?acc=GSE192405.
- **Fiorenzano et al. 2021** (3D) (Fiorenzano et al. 2021): It encompasses 91,034 organoid cells sampled at four differentiation timepoints: days 15 (n=2 organoids), 30 (n=5), 60 (n=5), 90 (n=2) and 120 (n=6). In those 20 samples, 8 cell types were identified, of which 16% of the cells were dopaminergic neurons. Raw counts were downloaded from the Gene Expression Omnibus (GEO) using the accession identifier GSE168323 at https://www.ncbi.nlm.nih.gov/geo/query/acc.cgi?acc=GSE168323. Only the 20 standard organoid samples (“*standardorgday*”) were merged for further analysis. Metadata containing cell type annotation was shared by the authors after request.
- **Zagare et al. 2022** (3D) (Zagare et al. 2022): It contains 6,000 organoid cells mimicking the human embryo ventral midbrain, sampled at two differentiation timepoints (days 35 and 70). Raw counts were downloaded from the Gene Expression Omnibus (GEO) using the accession identifier GSE133894 at https://www.ncbi.nlm.nih.gov/geo/query/acc.cgi?acc=GSE133894. The cell type annotation presented in the original publication was not available at the time of this integration. Only 2 wild-type samples were included for downstream analysis.
- **Agarwal et al. 2020** (PostMortem) (Agarwal et al. 2020): It contains 5,943 cells from 7 post-mortem substantia nigra samples, of which two are technical replicates (n=5 donors). 10 cell types were identified, of which only 1.2% were dopaminergic neurons. Raw counts were downloaded from the Gene Expression Omnibus (GEO) using the accession identifier GSE140231 at https://www.ncbi.nlm.nih.gov/geo/query/acc.cgi?acc=GSE140231. We merged the 7 samples profiling substania nigra (N). Metadata was downloaded from Supplementary Data 2 of the original publication.

For the integration, we used a later version of the Seurat package (v.5.0.0). We initially merged the raw counts of the seven datasets and normalized the data. Then, we found the highly variable features for each layer separately, using default parameters. After that, we scaled the data and calculated the first 50 principal components (PCs) and ran a fast implementation of the mutual nearest neighbors, FastMNN integration (package *SeuratWrappers, v.0.3.19*), using the *IntegrateLayers*() function and the previous dimensional reduction. We computed a neighbourhood graph, using 30 of the MNN embeddings, and performed cell clustering using the Louvain algorithm (resolution=0.75). We visualized the dimensional reduction with a UMAP projection using the same 30 MNN embeddings from the integration. After this, we harmonized the columns from the different datasets to have a coherent metadata. Linked to cell type annotation, we created two columns:

- “**annotation_mixed**”: This column mixed all the original annotations per publication in one column. Below you can find the columns names of the original metadata:

- Our reference: “seurat_clusters_24_Annot”
- La Manno et al.: “CellType”
- Braun et al: “CellClass”
- Birtele et al: “AnnotType”
- Fiorenzano et al: “NamedClusters”
- Agarwal et al: “Level_2_cell_type”
- “**annotation_unified**”: To compare cell type composition differences, we established a unified cell type annotation based on the published annotations of the datasets. You can find each cell type and the corresponding unified definition in **Table S6**.

### Pseudo-bulk transformation and correlation analysis

Leveraging the final integrated object, we excluded all those cells that did not correspond to a wild-type control (i.e. patient iPSC lines or edited lines). After that, we selected the 26 samples for pseudobulk analysis, generated by combining each dataset with its corresponding developmental timepoints (for fetal) or differentiation days (for 2D and 3D). For each sample, we computed the average expression per gene, considering only the 2,000 highly variable genes, and using the batch-corrected values from the FastMNN integration. We then computed the Pearson correlation of the pseudobulk expression between all pairwise combinations of in vitro (2D, 3D) and in vivo (fetal or postmortem) samples. The results from this analysis can be visualized in Fig 3B heatmap, that presents an additional hierarchical clustering of rows and columns to highlight the transcriptional similarity between samples. For that visualization, we used the ComplexHeatmap R package (v2.18.0).

### Pseudotime analysis

We performed pseudotime inference using three different R packages. Initially, we filtered-out cells belonging to rare cell types (n<50) based on the unified annotation of the integrated dataset. Additionally, cells from patient lines or edited lines were removed.

- **Slingshot (v.2.10.0)** (Street et al. 2018): Trajectory inference was performed using the *slingshot*() function, leveraging the MNN dimensional reduction from the integration, with the starting point set at floor-plate progenitors (FPP). We then extracted the pseudotime values corresponding to the maximum weight for each cell and lineage, ranking these pseudotime values into cell positions across the trajectory to ensure comparability with other pseudotime inferences.
- **Destiny (v.3.16.0)** (Angerer et al. 2016): The integrated Seurat object was converted to a SingleCellExperiment object for compatibility. We then created a diffusion map of the cells using the MNN dimensional reduction to model differentiation-like dynamics with the *DiffusionMap*() function. The map includes the diffusion components (eigenvectors) and their importance (eigenvalues of the diffusion distance matrix). The first diffusion component was used as a proxy for pseudotime and cells were ranked for comparability. Additionally, we computed the diffusion pseudotime with the *DPT*() function, but found the temporal ordering was not better resolved and therefore not used in downstream analysis.
- **Monocle3 (v.1.2.9)** (Cao et al. 2019): The integrated Seurat object was converted to a Monocle3 object using the *new_cell_data_set*() function to transfer the expression matrices, metadata and feature attributes. Additionally, the UMAP embeddingsgenerated with the MNN dimensional reduction during the integration were also transferred. Cells were clustered with a resolution of 10^-4^ using the UMAP dimensionality reduction to define the partitions. We then learned the principal graph trajectory that cells follow through the UMAP space within each partition. Afterward, cells were ordered according to pseudotime on the learned trajectory, with the start node set within the population of floor plate progenitors (FPP). This pseudotime was later ranked for comparability with the other tools. Following pseudotime inference, we tested genes for differential expression across the pseudotime and selected those that significantly changed (q-val<0.05, Moran’s I test). We then used the list of pseudotime-dependent genes to cluster them into modules that are co-expressed across cells. Using a random seed (123) and a Louvain resolution list (10^-6^ to 0.1), we obtained 50 gene modules of varying gene size.

Then, we compared the pseudotime distribution of neurons (as defined by the unified annotation) among the developmental timepoints and the differentiation days of each dataset (Wilcoxon test). Further, we also compared the pseudotime distribution of the dopaminergic subtypes presenting at least 50 cells as defined in our annotation for all the existing differentiation days or developmental timepoints.

We also computed the module activation scores (aggregated expression of log-normalized values) for each cell type to identify those modules whose expression was specific or enriched for a given cell type. We used the *aggregate_gene_expression*() function and the cell types defined in the unified annotation for that purpose. After selecting the five most highly activated modules in neurons (modules 6, 12, 18, 21 and 46), we performed gene ontology enrichment analysis to identify the potential biological processes driven by the genes in those modules. Using the *gost*() function from the *gprofiler2* R package again, we set the significance threshold at an adjusted P-value<0.05 using the g:SCS method for multiple-test correction. We defined the gene universe in the “custom_bg” and used the functional annotation from the Cellular Component category of Gene Ontology (GO:CC) (Gene Ontology et al. 2023). Based on the term size, the difference between the effective domain size and the term size, the query size and the intersection size, we computed the odds ratio for each term.

Finally, we also computed the module activation scores for each cell to compare the expression profile of the neuron-enriched modules in the neurons profiled in our data versus the neurons from other datasets (unified annotation).

- This work (Fetal) vs Birtele et al. 2022 (Fetal)
- This work (Fetal) vs Braun et al. 2023 (Fetal)
- This work (Fetal) vs La Manno et al. 2016 (Fetal)
- This work (in vitro) vs This work (Fetal)
- This work (in vitro) vs Fiorenzano et al. 2021 (3D)

Given the differences in neuron cell number between the datasets, we performed a permutation test (*oneway_test*() function) resampling from the smaller group in each case (n=100 times, mean p-value reported). For each comparison, we set the significance threshold at an adjusted P-value<0.05 using the Benjamini & Hochberg multiple-test correction. The significance levels are indicated as follows: pAdj<0.001 (***), pAdj<0.01 (**) or pAdj<0.05 (*). Additionally, to capture the dynamics of gene module expression in neurons throughout midbrain development, as well to detect batch differences between datasets, we calculated the neuron-enriched module activation scores (mean ± standard deviation) for each of the developmental timepoints or differentiation days per dataset.

### Spatial transcriptomics

Snap frozen, OCT embedded tissue blocks were cryosectioned on a Bright OTF5000 cryostat. 10um sections were loaded onto Visium Spatial Gene Expression slides (product code: 1000184, 10X Genomics, Pleasanton, CA) as per 10X Genomics demonstrated protocol: CG000240. Triplicate sections of each of four gestational ages (7, 11, 12 and 17 PCWs) of brain tissue were randomized across the top three capture areas of 4 slides to avoid any potential batch effect. The fourth capture area of each slide was loaded with sections of OCT blocks, each containing six organoids randomized by donor, time point, differentiation and duplicate. Loaded Visium slides were subsequently methanol fixed, stained with Haematoxylin and Eosin (H&E) and imaged as per 10X Genomics demonstrated protocol: GC000160 on a Motic EasyScanOne slide scanner, at 40X magnification. Following 10X Genomics user guide: CG000239, sections were then subjected to permeabilization for mRNA capture, reverse transcription and second strand synthesis to generate full length cDNA, followed by library preparation. The permeabilization time of 12 minutes for both brain tissue and organoids was established as appropriate using the Visium Spatial Tissue Optimisation kit (product code: 1000193, 10X Genomics, Pleasanton, CA) following 10X Genomics demonstrated protocols and user guide: CG000240 for sectioning and slide loading, GC000160 for H&E staining and brightfield imaging and CG000238 for mRNA capture, reverse transcription, second strand synthesis and fluorescent imaging. The resulting libraries were sequenced on an Illumina Novaseq 6000 sequencer, using a Novaseq 6000 S2 (100 cycle) v2 sequencing kit.

### Spatial Transcriptomics Computational analysis

We used spaceranger (v2.1.0)(Cable et al. 2022) from 10X Genomics to process the bcl files obtained from the sequencer following spatial transcriptomics (ST) sequencing. We generated FASTQ files and gene expression matrices with the mkfastq and count commands with the ENSEMBL homo sapiens reference genome build GRCh38. For each sample, the FASTQ files, gene expression matrix and spaceranger output directory are available at the Gene Expression Omnibus (GEO data will be made public after publication).

We conducted the subsequent analysis in Seurat (v4.4.0) (Hao et al. 2021), writing the code as an Rmarkdown notebook. All analysis was processed within Docker, and the images and code used are available on Dockerhub (will be made public after publication). The analysis can be replicated on any computer where Docker is available by downloading the above data, code, and Docker images.

We removed the 12 PCW sections from the dataset as the tissue had been damaged during processing. We also removed spots that were disconnected from the main slice of tissue/organoid. We identified progenitor and post-mitotic cell populations by clustering the data using Seurat’s FindClusters function with sample-specific resolutions that generated two clusters. These two clusters were then annotated using marker genes and anatomical knowledge.

Since each spot is likely to contain multiple cells (possibly of different cell types), we ran cell type deconvolution using Cell2Location (v0.1.3) (Kleshchevnikov et al. 2022) in conjunction with the annotated single cell reference that we developed in this paper.

To identify genes that are differentially expressed with respect to maturation, we first manually annotated the path of maturation on each sample and then used this as a covariate in CSIDE (from spacexr package v2.2.1) (Cable et al. 2022). We provided CSIDE with our cell type deconvolutions, which allowed us to identify maturation-related differentially expressed genes for each cell type. We calculated the running weighted average of several genes of interest by averaging their expression along the path of maturation using a window size of 50 spots.

We profiled cell-cell interaction using COMMOT (v0.0.3) (Cang et al. 2023), which identifies the most-active spatially co-located ligand-receptor pairs, where the pairs were taken from the CellChat database (v1) (Jin et al. 2021). We used Moran’s I (Moran 1950) to identify the ligand-receptor pairs with the highest spatial autocorrelation (i.e. where their position appears to explain their expression).

To quantify the spatial similarity between ST samples, we first labelled each spot as the cell type for which the deconvolution estimates it to have in the greatest abundance. We then counted the number of neighboring spots for each pair of cell types. We normalized these counts and used them to compute (Euclidean) distances between samples. Inverting and normalizing these distances to the range [0, 1] allowed us to treat them as percentages of similarity.

We estimated the temporal alignment of each organoid by computing a weighted average of the timepoints of the tissue samples, where weights were taken with respect to the organoid’s spatial similarity to each timepoint.

### Immunoblotting

Proteins were extracted from cells in ice-cold RIPA lysis and extraction buffer (Sigma-Aldrich) supplemented with protease inhibitor (Roche). Protein concentration was measured with Pierce™ BCA Protein Assay kit (Thermo Scientific): 10 µg of protein was denatured with Laemmli buffer (Bio-Rad Laboratories LTD) with dithiothreitol (DTT). Proteins were separated with Mini-PROTEAN TGX Stain Free Gels (Bio-Rad Laboratories LTD) and transferred to a Trans-Blot Turbo Transfer membrane (Bio-Rad Laboratories LTD). After blocking in 5% milk, 1x PBS, 0.1% Tween for 1h at room temperature, membranes were incubated with primary antibodies (Table S12) at 4°C overnight. Membranes were then incubated with the secondary anti-rabbit horseradish peroxidase-conjugated antibody at a dilution of 1:3000 (Cell Signalling). Immunoreactive proteins were visualized with Chemidoc MP (Bio-Rad Laboratories). Equal loading was evaluated with membranes reprobing for GAPDH, after clearance with Restore Western Blot Stripping Buffer (Thermo Scientific). The intensity of immunoreactive bands was analyzed using ImageJ software (National Institutes of Health). The density of the bands was normalized to GAPDH. Results are reported as means SEM of independent experiments, the number of which is stated for each experiment in the respective figure legend.

## Statistical analyses

For the statistical analysis of iPSCs derived data, two-tailed Student’s t-test was applied for dual analysis comparison. Results are reported as mean ± standard error or the mean (SEM) from at least three independent biological replicates. Significant levels were determined by P-value. Analysis was performed using GraphPad Prism.

## Ethical approval

The national ethical commission has approved the work with induced pluripotent stem cells under the REC reference number 13/LO/0171, IRAS project ID: 100318.

Fetal samples have been collected under the ethical approval to HDBR: REC reference number 23/NE/0135, IRAS project ID: 330783 and REC reference number 23/LO/0312, IRAS project ID: 326492.

## Data availability

Raw FASTQ files for both single-cell and spatial transcriptomics will be deposited in Gene Expression Omnibus (GEO). Four supplemental data files will be made available in a Zenodo repository, detailed as follows:

**Data S1-S2:** VCF files with donor genotypes to deconvolute pooled scRNA-seq data.

**Data S3:** Single-cell data generated in our study (RDS object: Seurat object with counts and metadata).

**Data S4:** Integrated datasets (RDS object: Seurat object with counts and metadata).

## Code availability

Two GitHub repositories will be made available upon publication: one for single-cell transcriptomics and another for spatial transcriptomics. These repositories will include scripts for processing steps, data analysis and figure generation. Additionally, the Docker image used for spatial transcriptomics analysis will be made available on Dockerhub following the publication.

## Acknowledgements

We sincerely thank our patients and their families for participating in this study. S.B. is supported by the Great Ormond Street Hospital Children’s Charity. P.P.C and H.K. are currently funded by the Sigrid Jusélius Foundation and previously by the NIHR Great Ormond Street Hospital Biomedical Research Centre. G.T.H. is funded by the NIHR Great Ormond Street Hospital Biomedical Research Centre. J.J. was supported by a postdoctoral fellowship from OpenTargets. We thank UCL Genomics (UCL GOS Institute of Child Health) and the Wellcome Sanger Institute for undertaking sc-RNA and spatial sequencing. We thank the Human Developmental Biology Resource (HDBR) for providing human midbrain fetal tissue. We thank the UCL Imaging Facility and the facility manager Dr Dale Moulding for the support with immunofluorescence analysis. This research was supported by the NIHR Great Ormond Street Hospital Biomedical Research Centre. The views expressed are those of the author(s) and not necessarily those of the NHS, the NIHR or the Department of Health.

## Author contributions

D.B., S.B. and M.A.K conceived and designed the study. S.B. designed and performed experiments and data analysis for the in vitro and in vivo study. D.B. designed and performed experiments and data analysis for the in vitro and in vivo study. P.P.C. designed and performed data analysis for the scRNA-seq study. G.T.H. designed and performed data analysis for the spatial transcriptomic study. C.R. performed experiments for the in vitro study. X.T. designed and performed experiments for the scRNA-seq and spatial transcriptomic studies. E.M. performed experiments for the spatial transcriptomics study. J.J. performed experiments for the scRNA-seq study. A.D.D. performed experiments for the in vitro study. S.C. supervised scRNA-seq and spatial transcriptomic studies. D.B., P.P.C., G.T.H., S.C., S.B., and M.A.K. drafted the manuscript. C.R. and X.T. contributed to written sections of the manuscript. All authors reviewed the manuscript prior to submission.

## Additional information

All supplementary tables have been merged in one excel file named “Budinger et al_supplementaryTables”, which will be made available upon request.

---

**Supplementary Fig 1.**
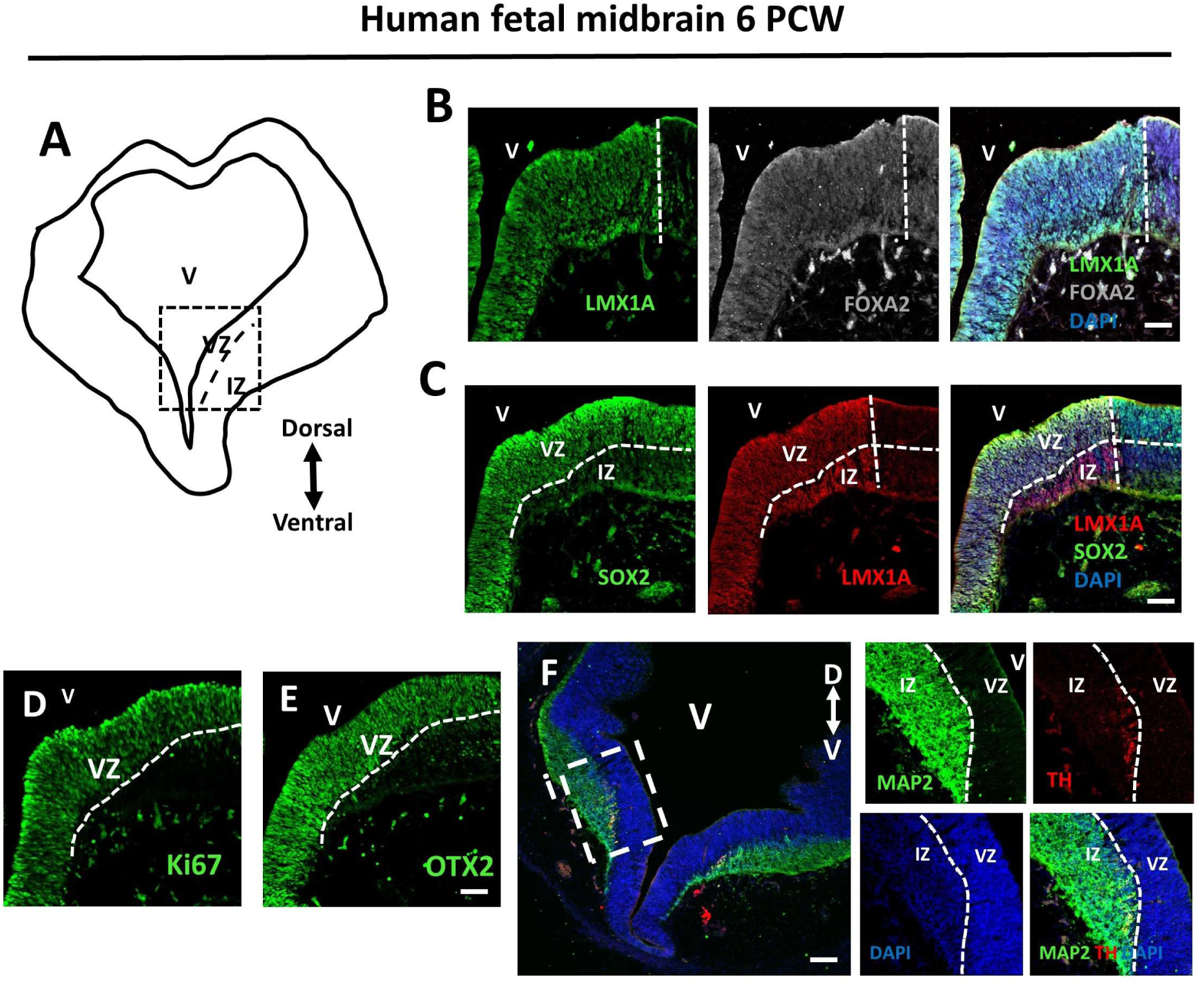
Human ventral midbrain at early stage of development. **A**, graphical representation of a coronal section of the human midbrain at 6 pcw. **B**, immunofluorescence for LMX1A and FOXA2 in the midbrain ventral ventricular zone (VZ). **C**, immunofluorescence staining for SOX2 and LMX1A in the ventral midbrain. **D-E**, staining for Ki67 and OTX2 of the human midbrain at 6 pcw. **F**, immunofluorescence analysis of a coronal section of the human ventral midbrain at 6 pcw for TH and MAP2. Nuclei are stained for DAPI. Lines demarcate the ventral midbrain region and the VZ to IZ boundary Scale bars= 100 μm (b, c, d and e); 200 μm (f). Ventricular zone (VZ), intermediate zone (IZ).

**Supplementary Fig 2.**
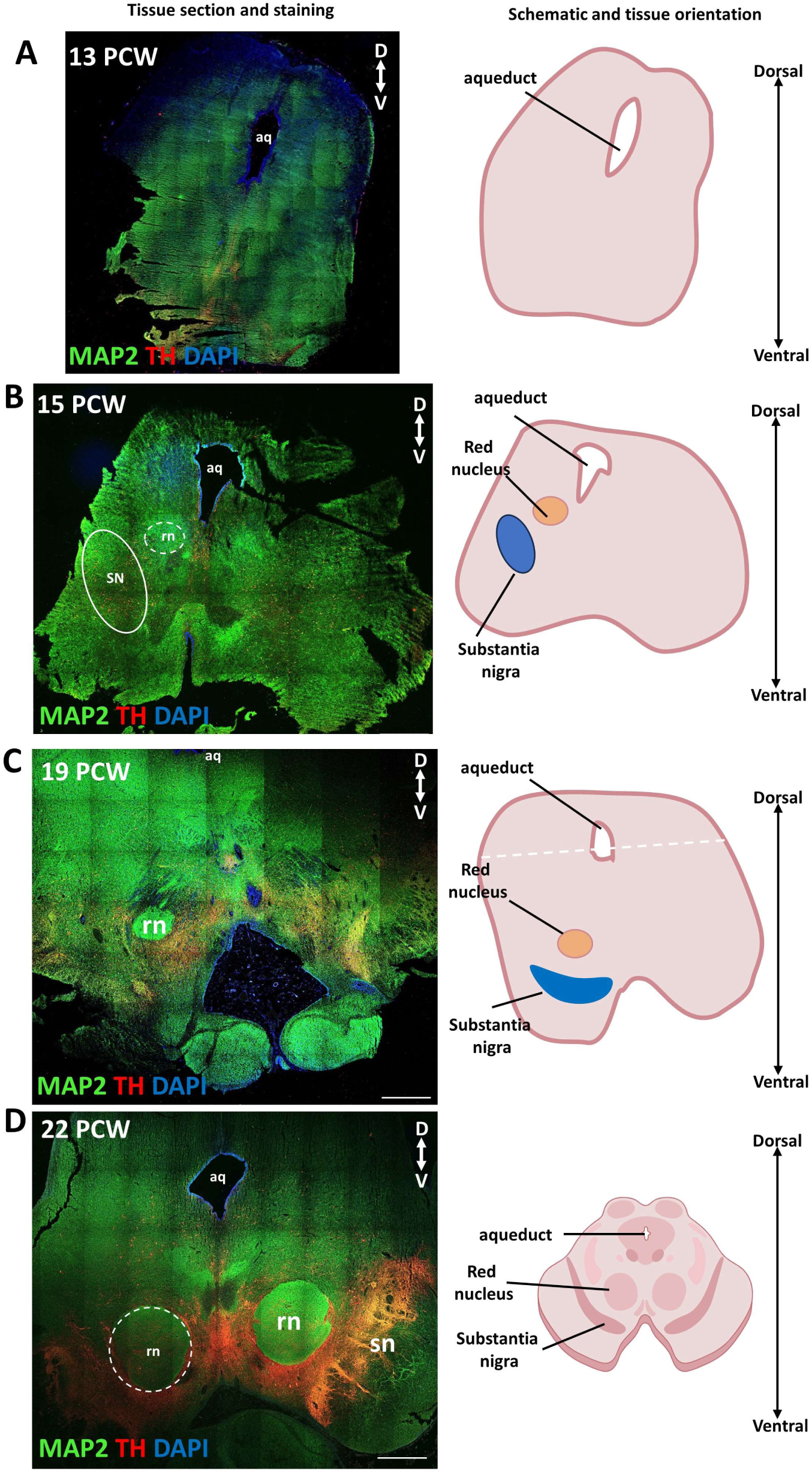
Human midbrain maturation during the second trimester. **A**, immunofluorescence representative images of human midbrain coronal sections at 13, 15, 19, 22 pcw stained for TH and MAP2 (*left panel*) and graphical representation of the section (*right panel*). Scale bar= 500 μm.

**Supplementary Fig 3.**
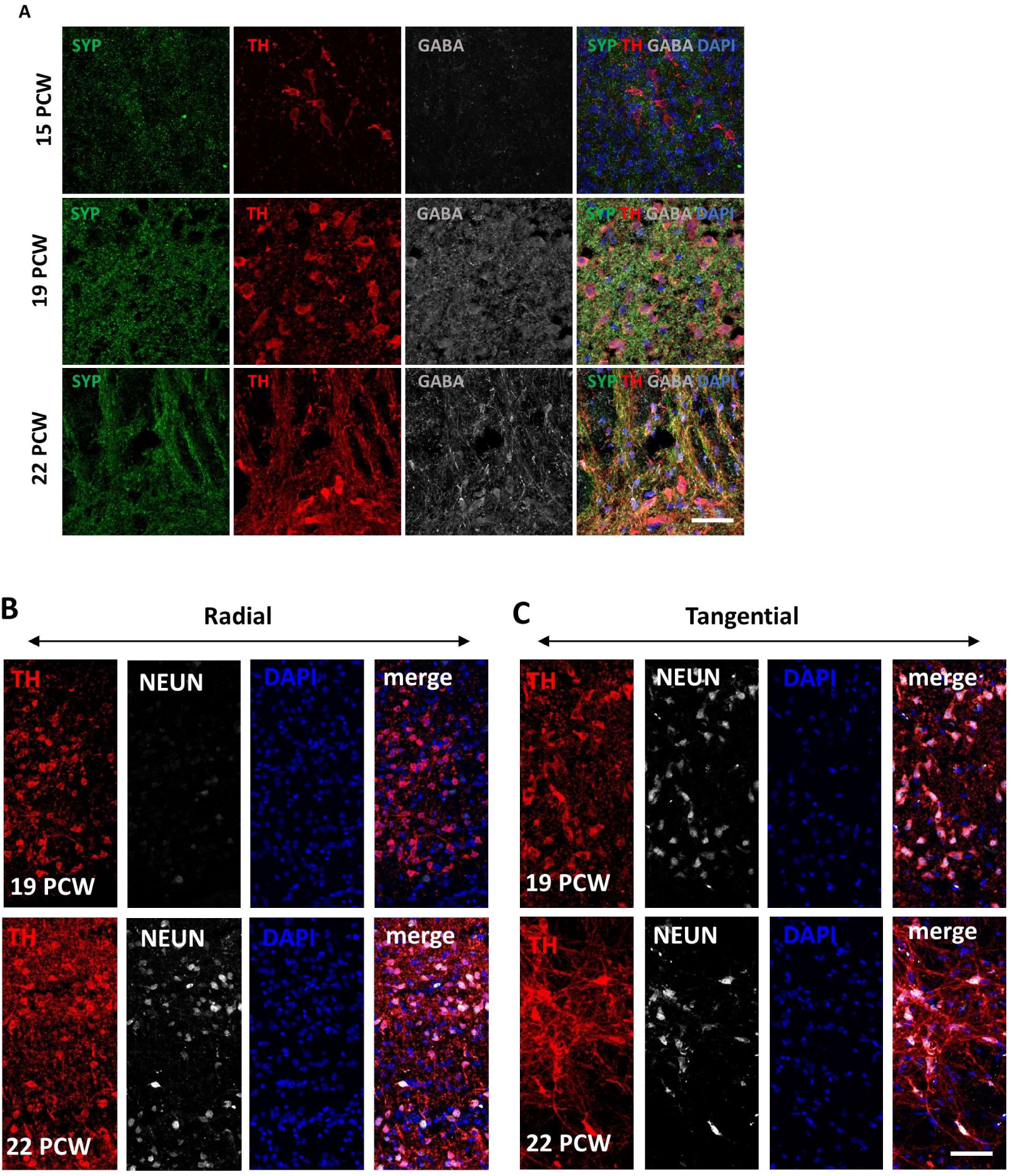
Human midbrain dopaminergic neurons subtypes and maturation stage. **A**, immunofluorescence analysis for Synaptophisin (SYP), TH and GABA of fetal samples at 15, 19 and 22 pcw. **B-C**, representative immunofluorescence images of radial and tangential migrating dopaminergic neurons (TH+) and maturation stage as indicated by NEUN positivity. Nuclei are stained for DAPI. Scale bar= 50 μm.

**Supplementary Fig 4.**
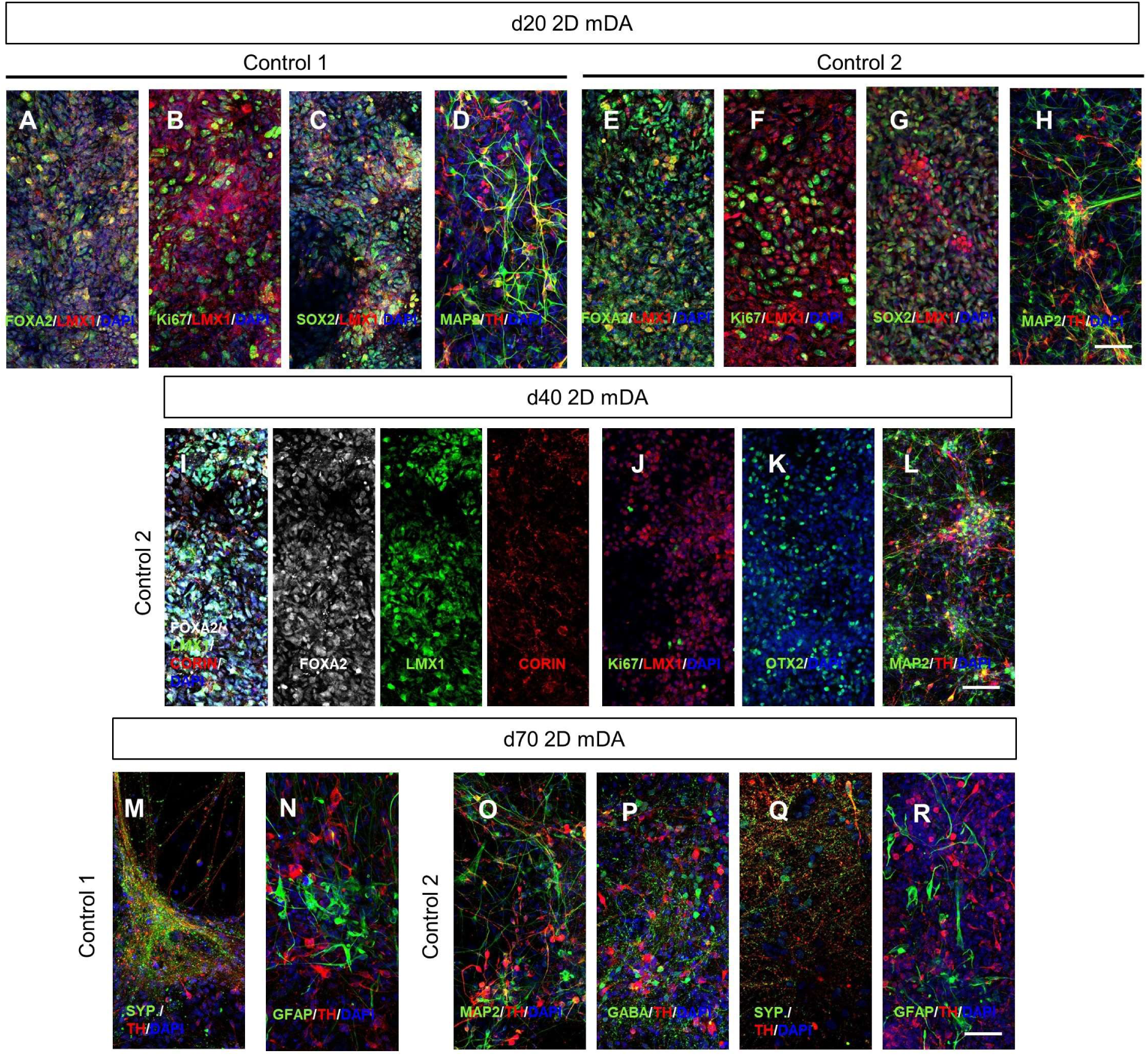
Characterization of iPSC-derived 2D mDA neuronal cultures. **A-R**, representative images of immunofluorescence analysis for midbrain and neuronal-related proteins FOXA2, LMX1, CORIN, SOX2, MAP2, TH, OTX2, GABA, SYP, the glial cell marker GFAP and the cell cycle protein Ki67 in control1 and control 2-derived 2D mDA neural cultures at 20, 40 and 70 days of differentiation. Nuclei are staining with DAPI. Scale bar= 50 μm.

**Supplementary Fig 5.**
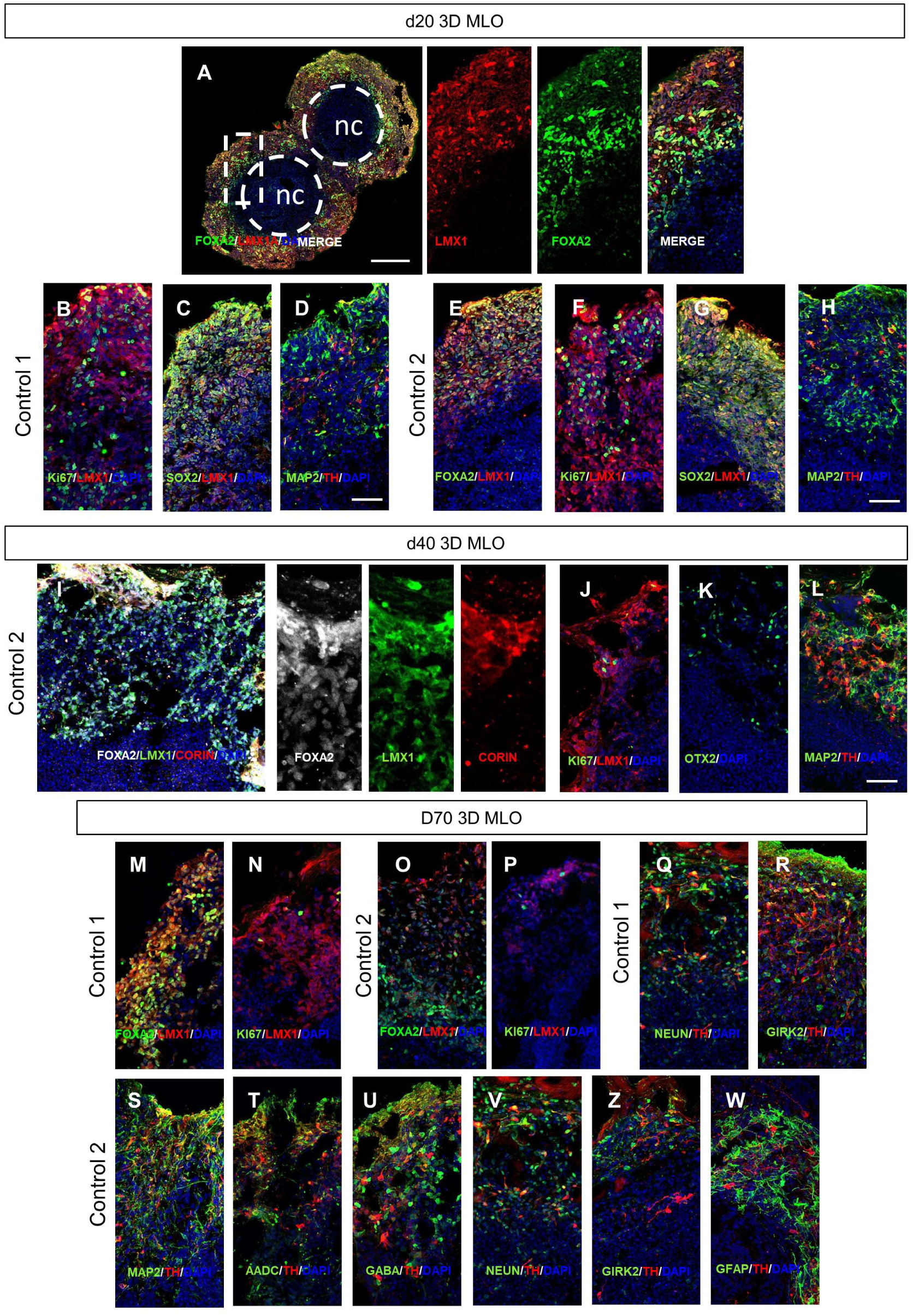
Characterization of iPSC-derived MLOs. **A-W**, representative images of immunofluorescence analysis for midbrain and neuronal-related proteins FOXA2, LMX1, CORIN, SOX2, MAP2, TH, OTX2, GABA, NEUN, GIRK2, the glial cell marker GFAP and the cell cycle protein Ki67 in control1 and control 2-derived MLO at 20, 40 and 70 days of differentiation. Nuclei are staining with DAPI. Scale bar= 50 μm.

**Supplementary Fig 6.**
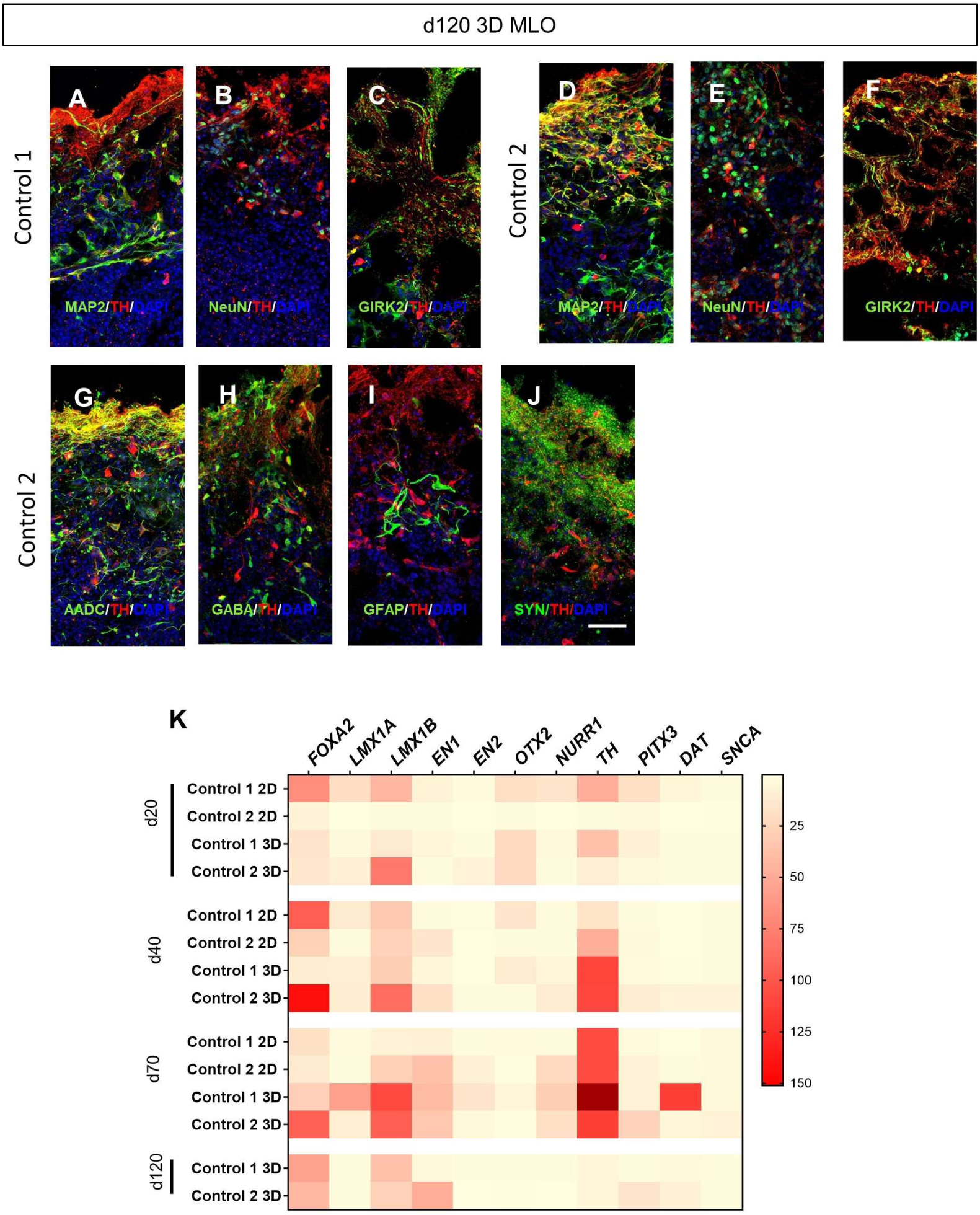
Characterization of iPSC-derived MLO at late stage of maturation. **A-J**, immunofluorescence images of control1 and 2-derived MLO at 120 days of differentiation showing positive cells for TH, MAP2, NEUN, GIRK2, GFAP and SYP. Nuclei are staining with DAPI. Scale bar= 50 μm. **K**, quantitative real time PCR (q-RT-PCR) for midbrain related genes in Control 1 and 2-drived 2D mDA (2D) cultures and MLOs at 20, 40, 70 and 120 days of differentiation. Genes expression for *FOXA2*, *LMX1A*, *LMX1B*, *EN1*, *EN2*, *OTX2*, *NURR1*, *PITX3*, *DAT*, *SNCA* is relative to housekeeping gene (GAPDH) and normalized to their respective iPSCs (n=1 for each line).

**Supplementary Figure 7.**
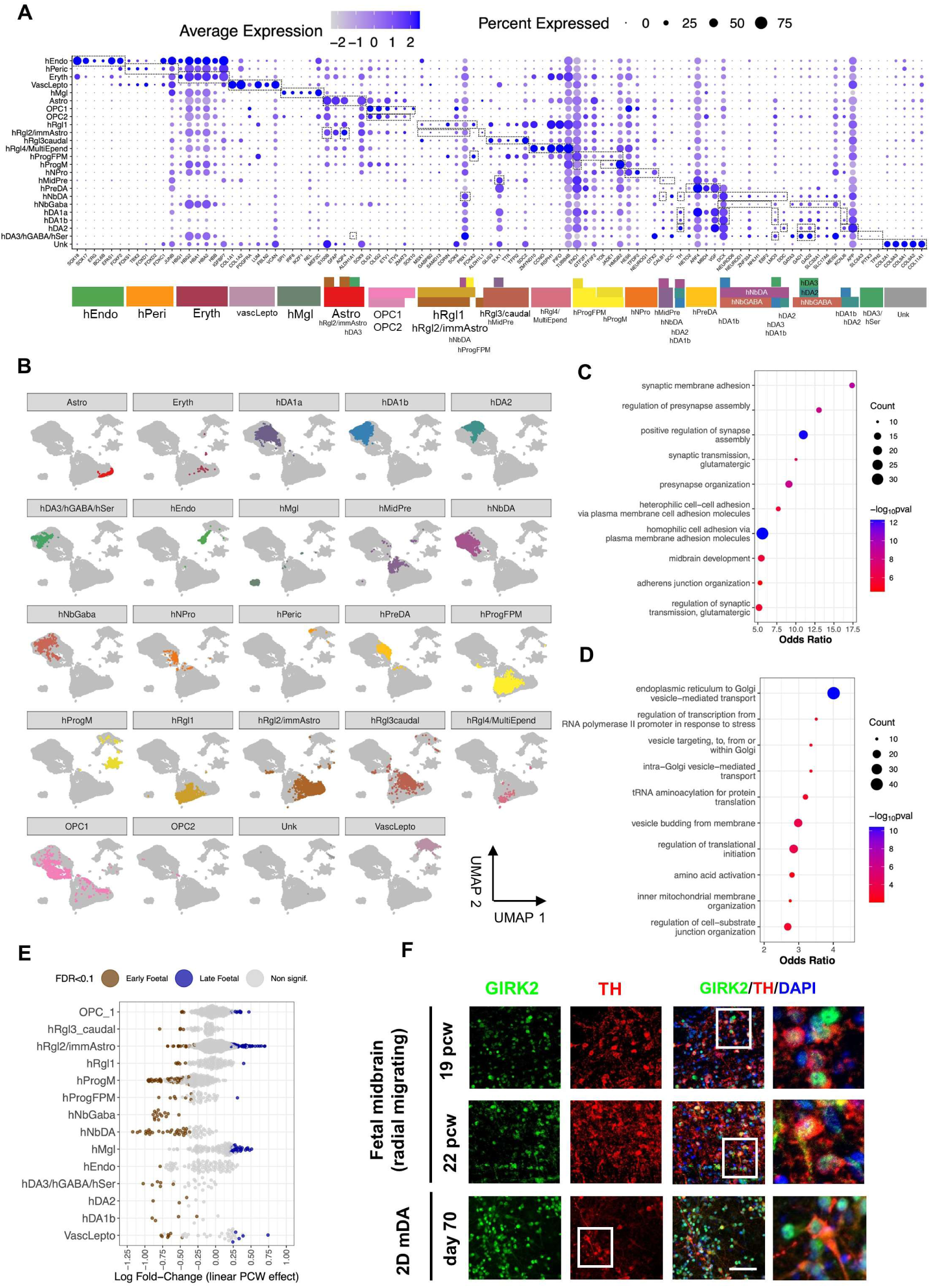
Single cell RNA-seq profiling of in vitro and in vivo foetal samples. **A,** Gene markers used for cell type annotation. The blue gradient indicates the level of averaged scale expression per cell type, while the dot size indicates the percentage of cells. Cell types are colored as in Fig. 3. For further information on markers and abbreviations, check (**Table 2**). **B**, UMAP visualization per cell type based on gene expression. Each facet only depicts one cell type at a time: astrocytes (Astro), erythrocytes (Eryth), dopaminergic neurons (hDA1a, hDA1b, hDA2), mixed group of dopaminergic, GABAergic-related and serotonergic neurons (hDA3/hGABA/hSer), endothelial cells (hEndo), microglia (hMgl), midbrain precursors (hMidPre), dopaminergic neuroblasts (hNbDA), GABAergic neuroblasts (hNbGaba), neuronal progenitors (hNPro), pericytes (hPeric), dopaminergic precursors (hPreDA), progenitors medial floorplate (hProgFPM), progenitor midline (hProgM), radial glia 1 (hRgl1), radial glia 2 / immature astrocytes (hRgl2/immAstro), radial glia 3 (hRgl3caudal), radial glia 4 / ependymal cells (hRgl4/MultiEpend), oligodendrocytes progenitor cells (OPC1, OPC2), vascular leptomeningeal cells (vascLepto) and unknown cells (Unk). **C**, Upregulation of synaptic-related pathways in fetal vs 3D hDA2 cells. Only the top-10 upregulated biological processes are shown (gene ontology enrichment parameters: pval<0.05, minSize=25, maxSize=500, minCount=10). **D**, Downregulation of transcription and translation-related pathways in fetal vs 3D hDA2 cells. Only the top-10 downregulated biological processes are shown (gene ontology enrichment parameters: pval<0.05, minSize=25, maxSize=500, minCount=10). **E,** Differential abundance analysis between early and late foetal cell types. As expected, microglia is enriched in groups of cells from the second trimester of fetal gestation, while neuroblasts (hNbGaba, hNbDA) and dopaminergic neurons (hDA2, hDA1b) are more abundant in the first trimester. Some cell types like OPC1 and hRgl2/immAstro show a dual enrichment within the same cell type, indicating large transitional cell states between the two trimesters.

**Supplementary Figure 8.**
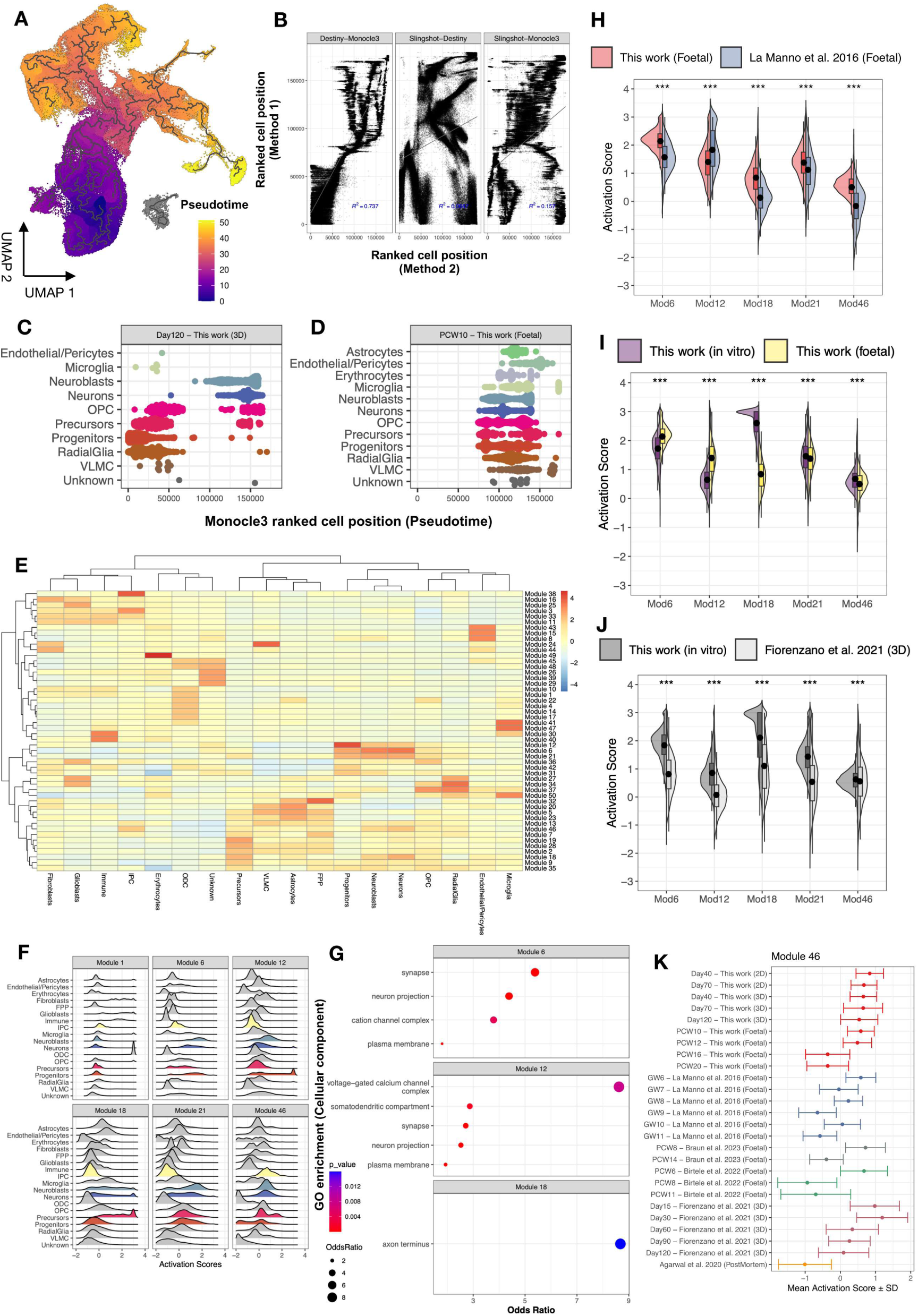
Meta-integration of multiple dopaminergic scRNA-seq datasets and analysis of pseudotemporal dynamics. **A,** Pseudotime trajectory inferred by Monocle3 on the UMAP projection. The gradient ranges from 0 (dark blue, rooted at floor plate progenitors, FPP) to 50 (yellow, corresponding to oligodendroctyes), spanning the learned trajectory of cells undergoing dopaminergic differentiation. **B**, Comparison of pseudotime trajectories. Pairwise correlation of the ranked cell position from the learned trajectories in Monocle3, Slingshot and Destiny. Only Monocle3 and Destiny show a high degree of agreement (R^2^=0.737). **C-D**, Violin scatter plot depicting pseudotime distribution for MLO and first-trimester fetal development. Ranked cell positions per cell type across the integrated dataset for 3D-dopaminergic model at day 120 (c) and for fetal development at post-conceptional week 10 (d), respectively. **E**, Heatmap displaying the activation scores of pseudotime-dependent gene modules across cell types. Each tile represents the average activation score (aggregated module scores) for each cell type (unified annotation in the columns). The inferred gene modules (rows, n=50) include groups of trajectory-variable genes that are co-regulated. **F**, Distribution of activation scores for a subset of gene modules. Ridge density plot showing the activation scores across cell types for the ODC-specific module (Module 1) and the top-enriched modules in neurons (Modules 6, 12, 18, 21 and 46). Neuron-enriched modules are typically also activated in neuroblasts and other cell types, such as progenitors (Modules 6 and 12), precursors (Module 18), and intermediate progenitor cells (Module 46). **G**, Gene ontology enrichment for neuron-enriched modules. Enriched terms per module (p<0.05, adjusted for multiple testing) highlight module 6 in synapse function, module 12 in voltage-gated calcium channel complexes, and module 18 in axonal structure. Modules 21 and 46 showed no significant enrichment in molecular functions, biological processes, cellular components, or KEGG pathways. Odds ratios were calculated based on term and query sizes, effective domain size and intersection size. **H-J**, Distribution of activation scores for neuron-enriched modules. Split violin plots illustrate the aggregated gene expression for five neuron-enriched modules defined by co-regulated trajectory-dependent genes. The comparisons include: fetal data from this study versus La Manno et al. 2016 (h); in-vitro data from this study versus all fetal datasets (i); and in vitro data from this study versus the 3D-dopaminergic model from Fiorenzano et al. 2021 (j), respectively. Significance level: p<10^-3^ (***, One-way permutation test). **K**, Activation score distribution for the neuron-enriched module 46. The scores are colored by dataset and displayed per timepoint (each row represents a different timepoint). The plot includes mean values with standard deviation error bars.

**Supplementary Figure 9.**
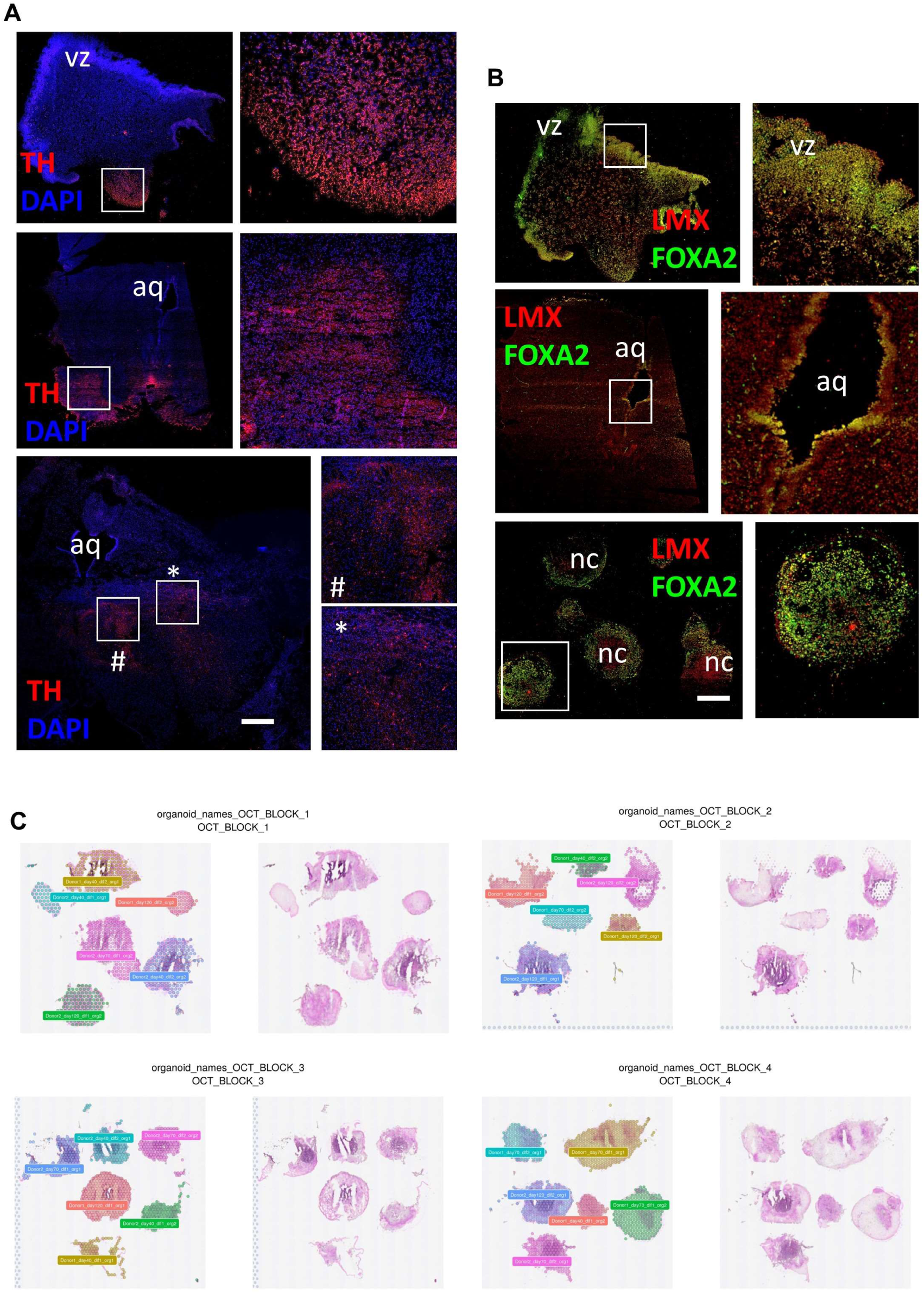
Midbrain fetal and MLO systems. **A**, immunofluorescence analysis for TH in fetal samples at 7, 11 and 17 PCW. Nuclei are stained with DAPI **B**, immunofluorescence analysis for LMX and FOXA2 in fetal samples at 7, 11 and 17 pcw. Scale bar = 500 µm. **C**, arrangement of MLOs in Visium capture areas.

**Supplementary Figure 10.**
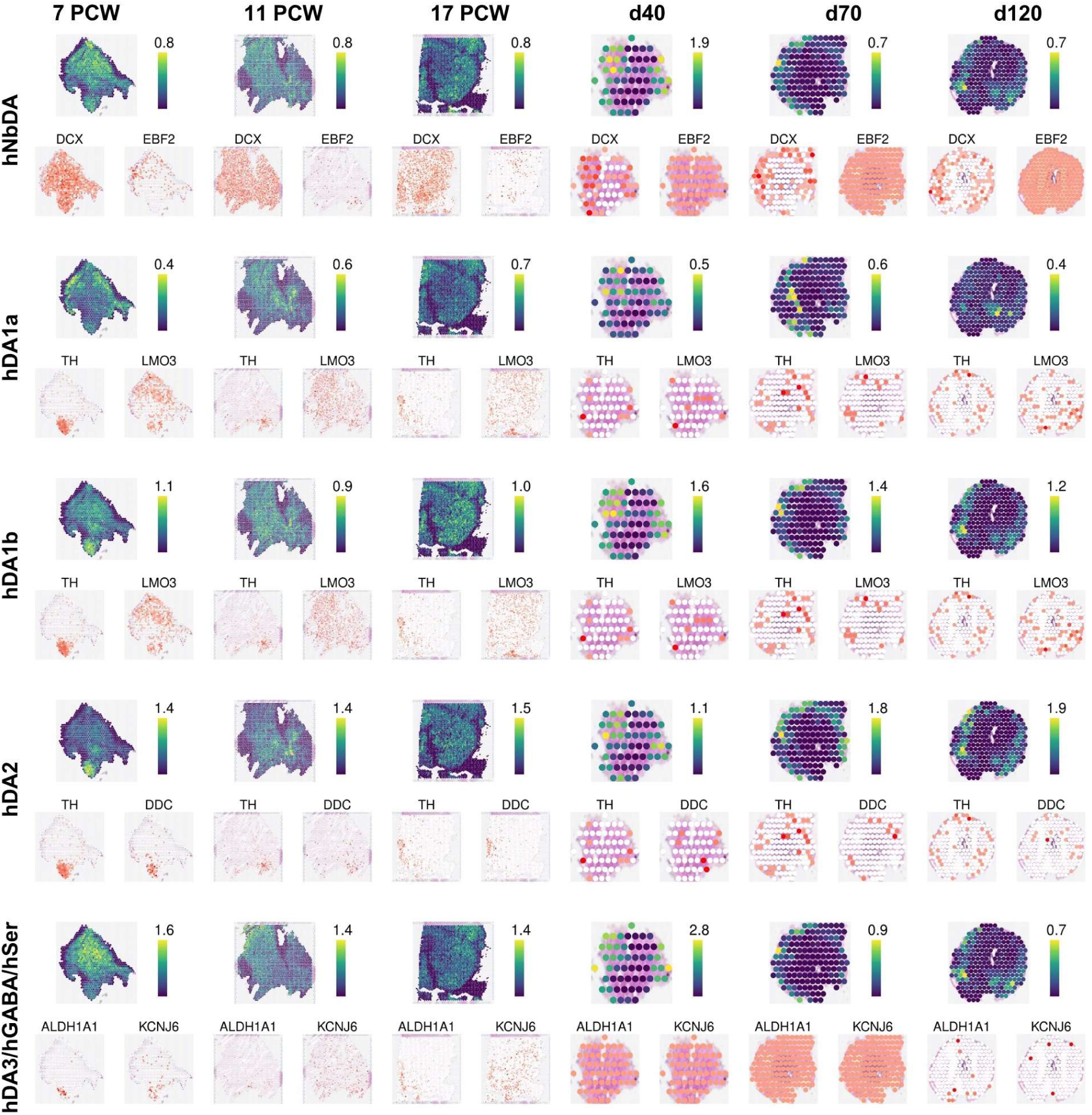
Distribution of dopaminergic populations in fetal tissue and MLOs. **A**, Abundance of dopaminergic lineage cells in tissues and organoids, as estimated by the Cell2Location deconvolution algorithm. Expression of two marker genes for each cell type are displayed below (white = low, red = high).

**Supplementary Figure 11.**
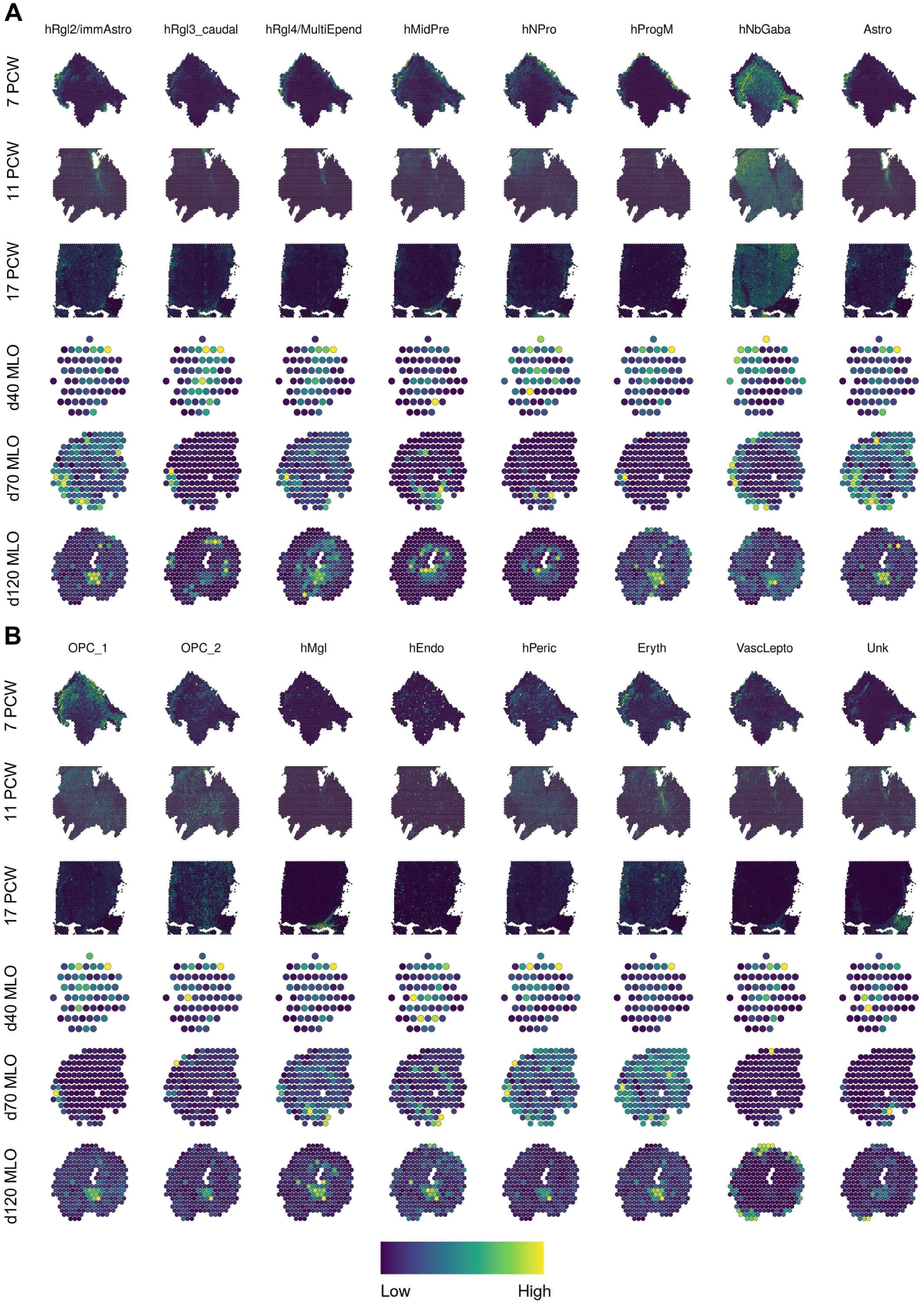
Distribution of cell types in fetal tissue and MLOs. **A**,**B**, Abundance of neuronal and non-neuronal cells in tissues and organoids, estimated by the Cell2Location deconvolution algorithm.

**Supplementary Figure 12.**
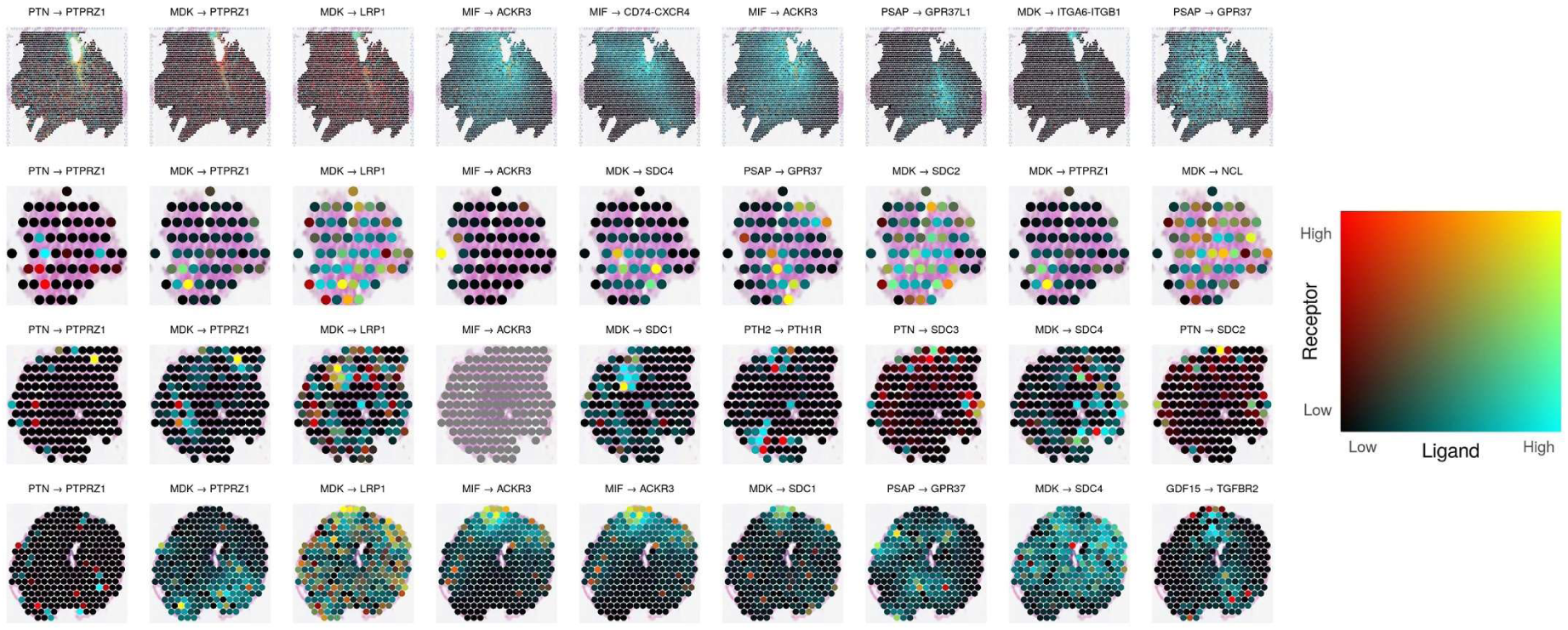
Distribution of cell-cell communication pathways in fetal tissue and MLOs. **A**, expression of pathways at each spot in 11 pcw fetal midbrain and MLOs at 40, 70 and 120 days of differentiation. Computed with COMMOT. Pathways selected according to spatial autocorrelation as determined by the Moran’s I non-parametric test.

**Supplementary Figure 13.**
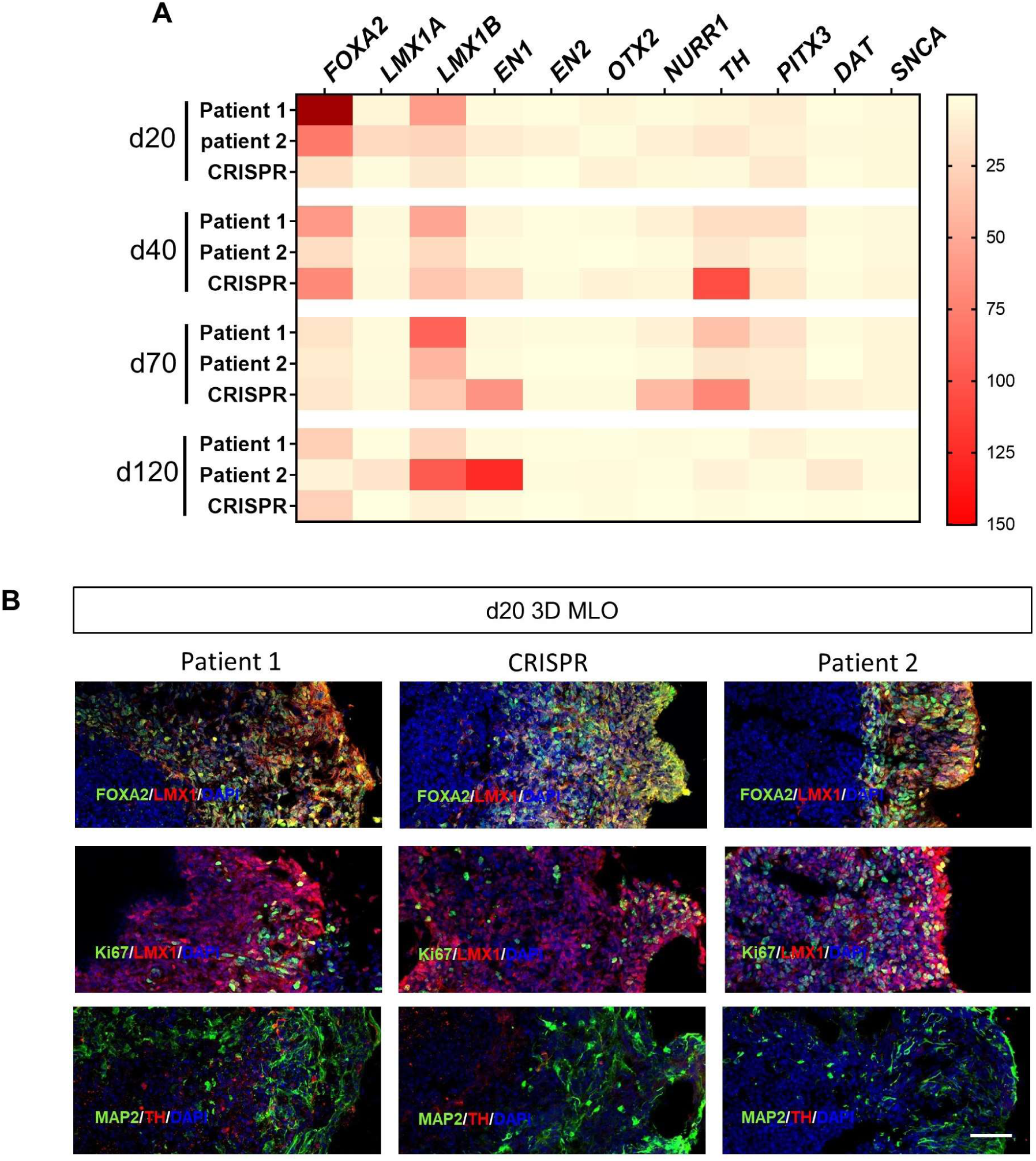
Characterization of DTDS derived MLOs. **A,** qRT-PCR for midbrain related markers *FOXA2*, *LMX1A*, *LMX1B*, *EN1*, *EN2*, *OTX2*, *NURR1*, *PITX3*, *DAT*, *SNCA* relative to housekeeping gene (GAPDH) and normalized to their respective iPSCs (n=1 for each line). **B**, immunofluorescence analysis for midbrain and neuronal-related proteins FOXA2, LMX1, MAP2 and TH, and Ki67 in Patient 1, Patient 2 and CRISPR corrected isogenic line-derived MLO at 20 days of differentiation. Nuclei are staining with DAPI. Scale bar= 50 μm.

**Supplementary Figure 14.**
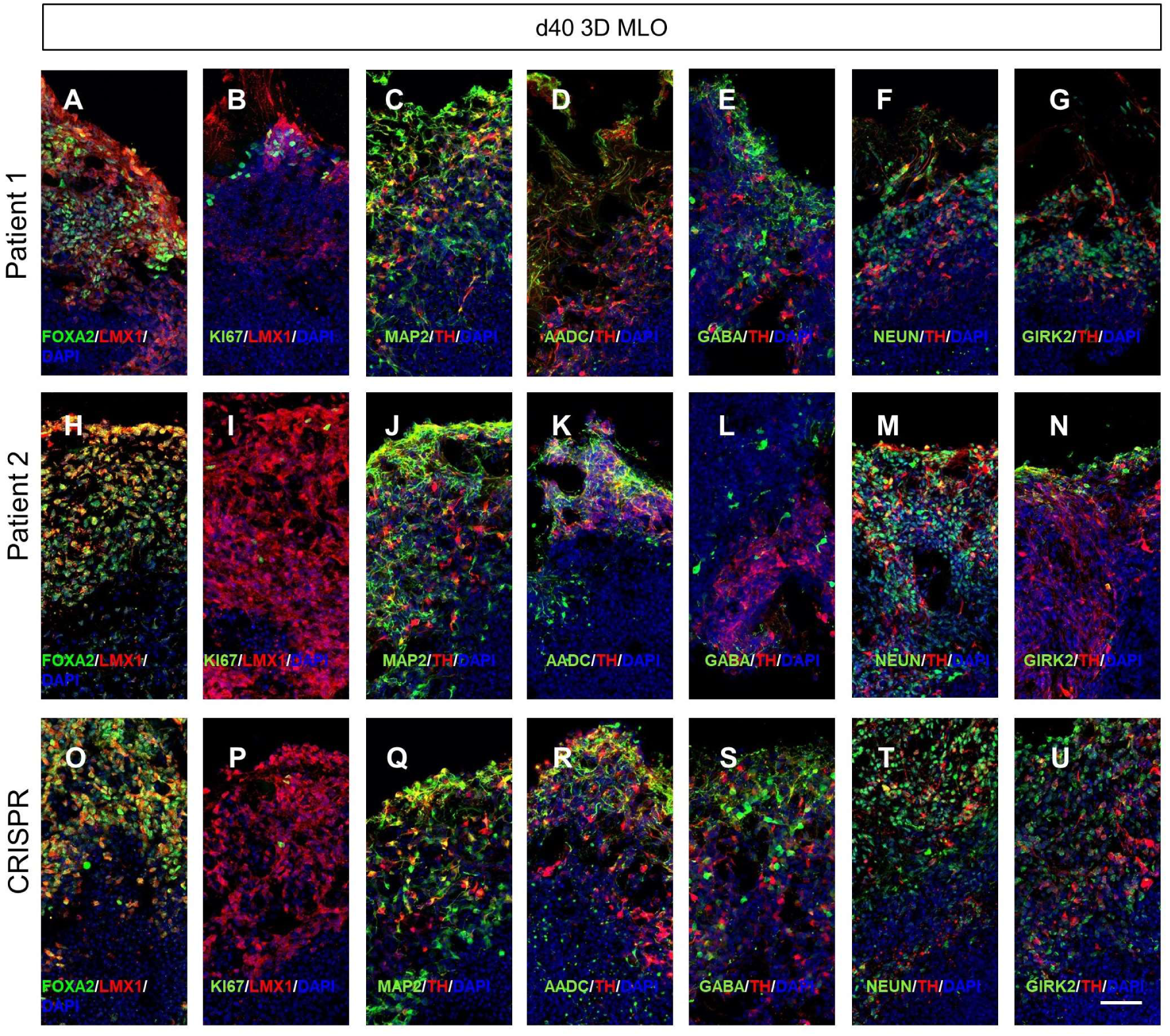
Maturation of mDA neurons in DTDS-derived MLOs at 40 days of differentiation. **A-U**, representative immunofluorescence images of Patient 1, Patient 2 and CRISPR isogenic lines showing expression of midbrain and neuronal-related proteins FOXA2, LMX1, MAP2, TH, AADC, GABA, NEUN and GIRK2 and the proliferative marker Ki67. Nuclei are staining with DAPI. Scale bar= 50 μm.

**Supplementary Figure 15.**
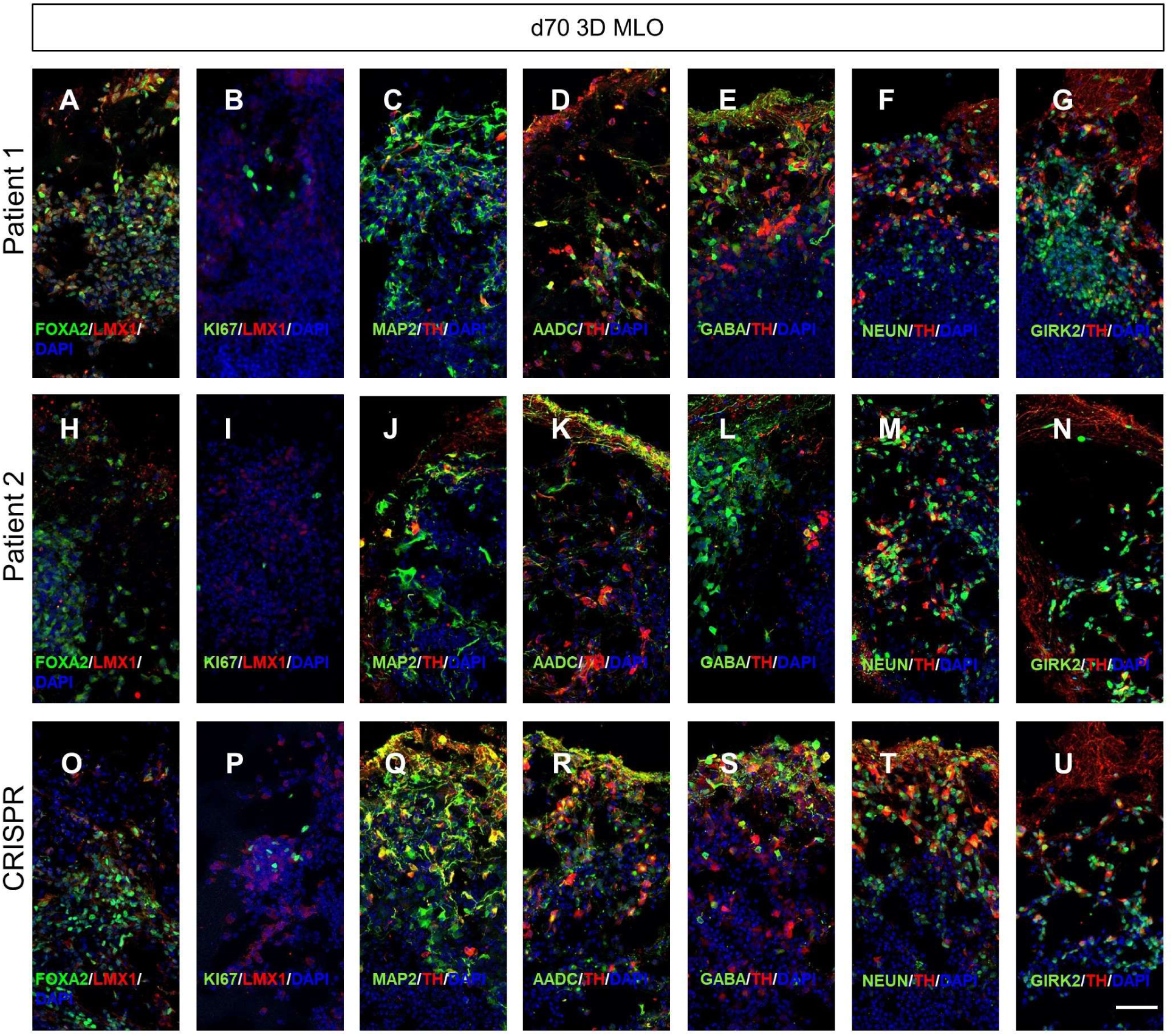
Maturation of mDA neurons in DTDS-derived MLOs at 70 days of differentiation. **A-U**, representative immunofluorescence images of Patient 1, Patient 2 and CRISPR isogenic lines showing expression of midbrain and neuronal-related proteins FOXA2, LMX1, MAP2, TH, AADC, GABA, NEUN and GIRK2 and the proliferative marker Ki67. Nuclei are staining with DAPI. Scale bar= 50 μm.

**Supplementary Figure 16.**
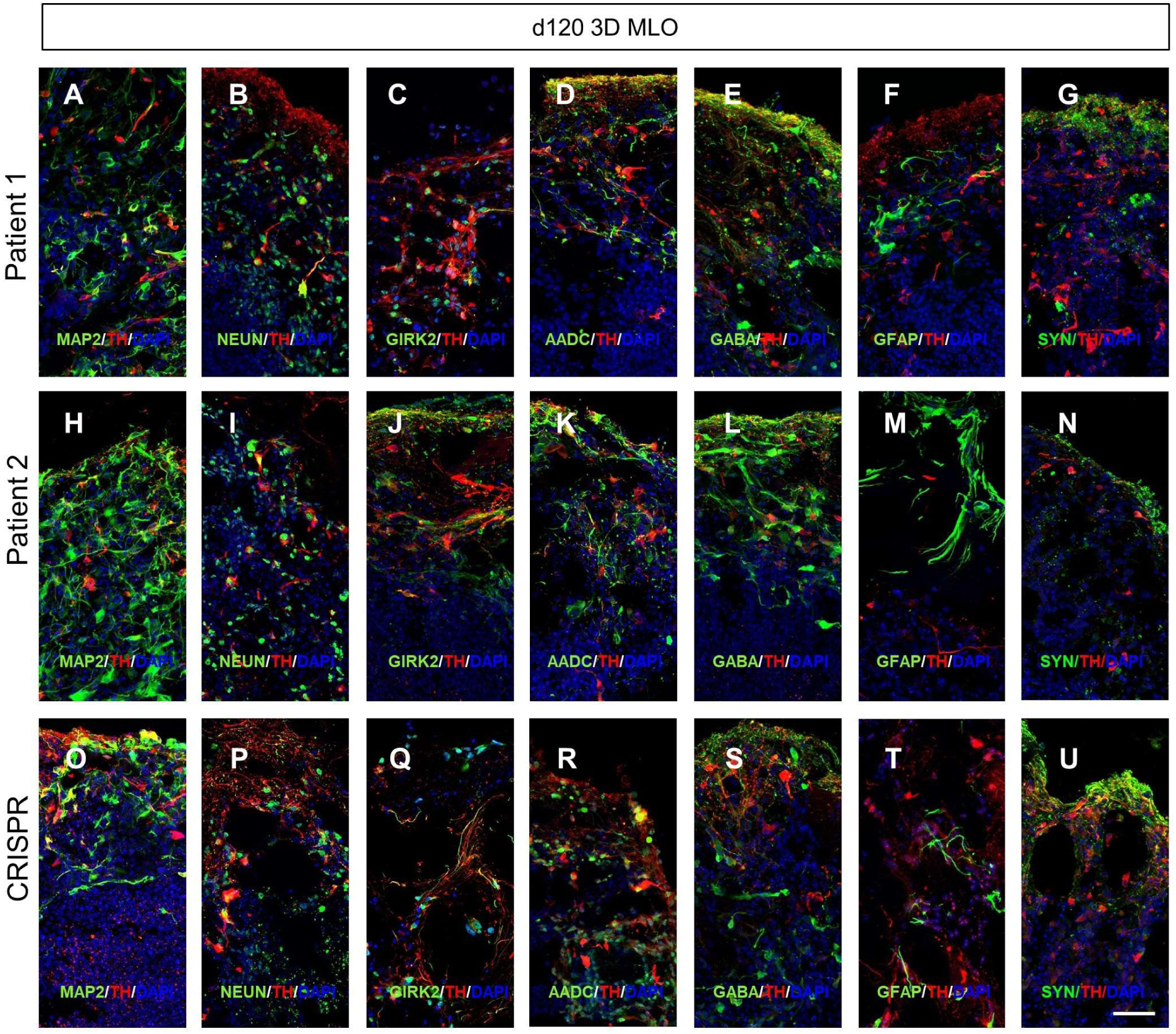
Maturation of mDA neurons in DTDS-derived MLOs at 70 days of differentiation. **A-U**, representative immunofluorescence images of Patient 1, Patient 2 and CRISPR isogenic lines showing expression of mDA neuron-related proteins TH, AADC and GIRK2, mature neuronal makers MAP2, NEUN, SYP, neural cells GABA positive cells and GFAP positive glial cells. Nuclei are staining with DAPI. Scale bar= 50 μm.

**Supplementary Figure 17.**
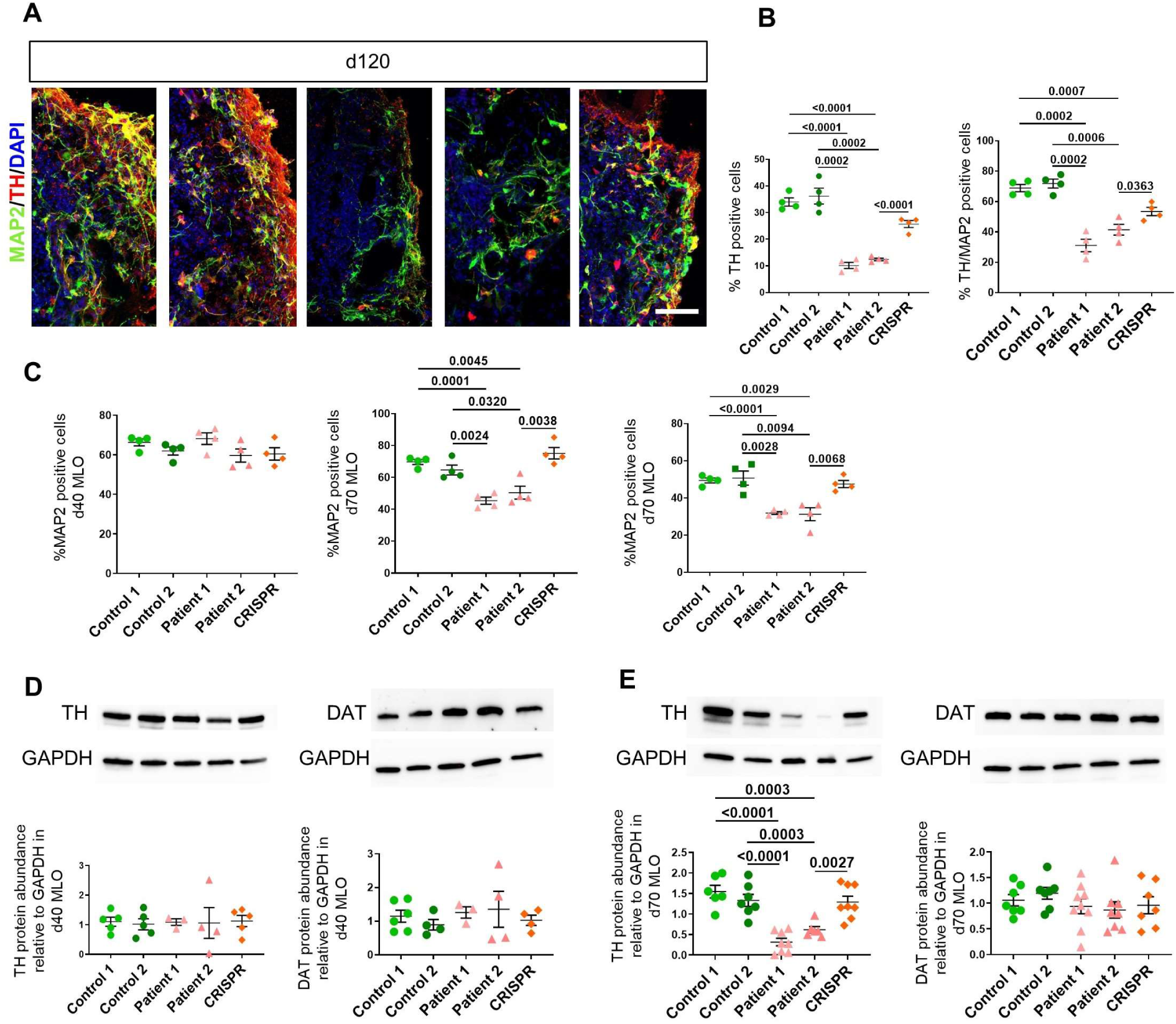
DTDS MLOs show loss of mDA neurons and reduced TH protein content. **A**, immunofluorescence analysis of control and DTDS patient-derived MLOs for TH and MAP2 at 120 days of differentiation. Nuclei are staining with DAPI. Scale bar= 50 μm. **B**, quantification of TH positive and TH/MAP2 double positive neurons for patients and controls lines at 120 days of differentiation. **c**, quantification of MAP2 total positive cells in DTDS control and patient-derived MLOs at 40, 70 and 120 days of differentiation. **D-E**, western blot analysis for TH and DAT and relative protein abundance to GAPDH in DTDS control and patient-derived MLOs at 40 and 70 days of differentiation respectively. Error bars indicate SEM. DTDS lines were independently compared to controls using two-tailed Student’s *t*-test for all analyses. Statistically significant differences indicate comparison of Control 1 or 2 to Patient 1 or 2 and Patient 2 to CRISPR.

**Supplementary Figure 18.**
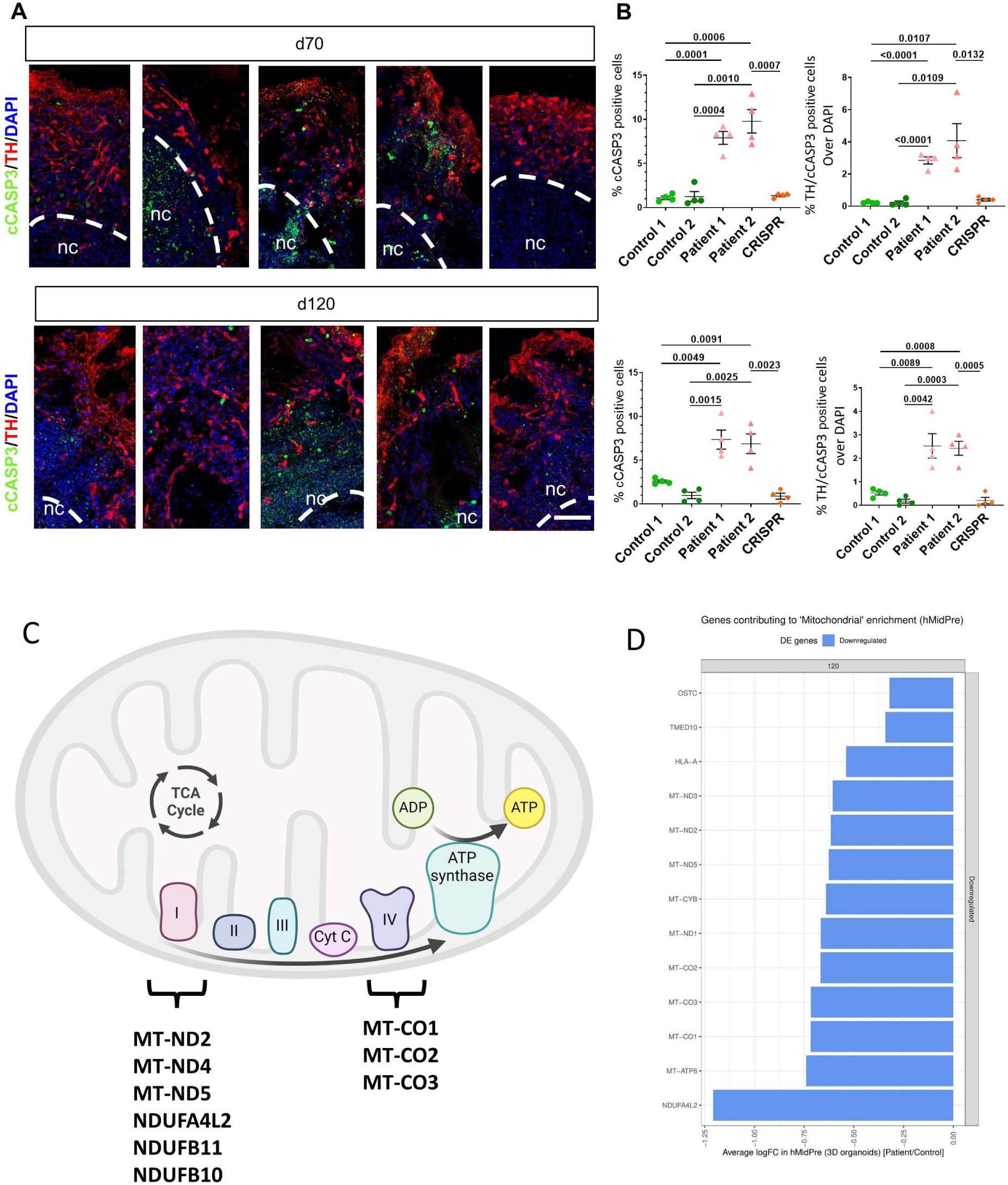
DTDS MLOs show increased levels of cCASP3 positive cells. **A**, immunofluorescence analysis of control and DTDS patient-derived MLOs for cleaved CASP3 (cCASP3) and TH at 40 and 70 days of differentiation. Nuclei are staining with DAPI. Positive cells located in the necrotic core (nc) were excluded from the quantification. Scale bar= 50 μm. **B**, quantification of total cCASP3 positive cells and TH/ cCASP3. Error bars indicate SEM. DTDS lines were independently compared to controls using two-tailed Student’s *t*-test for all analyses. Statistically significant differences indicate comparison of Control 1 or 2 to Patient 1 or 2 and Patient 2 to CRISPR. Necrotic core (nc). **C**, graphic representation of mitochondria genes dysregulated in DTDS mDA neurons and role in mitochondrial respiratory chain. **D**, Average LogFC for mitochondrial genes downregulated in patients MLOs compared to controls for midbrain precursors cell type (hMidPre).

## Reference

Abela, L., L. Gianfrancesco, E. Tagliatti, G. Rossignoli, K. Barwick, C. Zourray, K. M. Reid, D. Budinger, J. Ng, J. Counsell, A. Simpson, T. S. Pearson, S. Edvardson, O. Elpeleg, F. M. Brodsky, G. Lignani, S. Barral and M. A. Kurian (2024). "Neurodevelopmental and synaptic defects in DNAJC6 parkinsonism, amenable to gene therapy." Brain 147(6): 2023–2037.

Agarwal, D., C. Sandor, V. Volpato, T. M. Caffrey, J. Monzon-Sandoval, R. Bowden, J. Alegre-Abarrategui, R. Wade-Martins and C. Webber (2020). "A single-cell atlas of the human substantia nigra reveals cell-specific pathways associated with neurological disorders." Nat Commun 11(1): 4183.

Almqvist, P. M., E. Akesson, L. U. Wahlberg, H. Pschera, A. Seiger and E. Sundstrom (1996). "First trimester development of the human nigrostriatal dopamine system." Exp Neurol 139(2): 227–237.

Andersson, E., U. Tryggvason, Q. Deng, S. Friling, Z. Alekseenko, B. Robert, T. Perlmann and J. Ericson (2006). "Identification of intrinsic determinants of midbrain dopamine neurons." Cell 124(2): 393–405.

Angerer, P., L. Haghverdi, M. Buttner, F. J. Theis, C. Marr and F. Buettner (2016). "destiny: diffusion maps for large-scale single-cell data in R." Bioinformatics 32(8): 1241–1243.

Arenas, E., M. Denham and J. C. Villaescusa (2015). "How to make a midbrain dopaminergic neuron." Development 142(11): 1918–1936.

Asgrimsdottir, E. S. and E. Arenas (2020). "Midbrain Dopaminergic Neuron Development at the Single Cell Level: In vivo and in Stem Cells." Front Cell Dev Biol 8: 463.

Barral, S. and M. A. Kurian (2016). "Utility of Induced Pluripotent Stem Cells for the Study and Treatment of Genetic Diseases: Focus on Childhood Neurological Disorders." Front Mol Neurosci 9: 78.

Billett, H. H. (1990). Clinical Methods: The History, Physical, and Laboratory Examinations. 3rd edition. Boston: Butterworths; 1990. Chapter 151.

Birtele, M., P. Storm, Y. Sharma, J. Kajtez, J. N. Wahlestedt, E. Sozzi, F. Nilsson, S. Stott, X. L. He, B. Mattsson, D. R. Ottosson, R. A. Barker, A. Fiorenzano and M. Parmar (2022). "Single-cell transcriptional and functional analysis of dopaminergic neurons in organoid-like cultures derived from human fetal midbrain." Development 149(23).

Bissonette, G. B. and M. R. Roesch (2016). "Development and function of the midbrain dopamine system: what we know and what we need to." Genes Brain Behav 15(1): 62–73.

Bloch, B., E. Normand, I. Kovesdi and P. Bohlen (1992). "Expression of the HBNF (heparin-binding neurite-promoting factor) gene in the brain of fetal, neonatal and adult rat: an in situ hybridization study." Brain Res Dev Brain Res 70(2): 267–278.

Bodea, G. O., J. H. Spille, P. Abe, A. S. Andersson, A. Acker-Palmer, R. Stumm, U. Kubitscheck and S. Blaess (2014). "Reelin and CXCL12 regulate distinct migratory behaviors during the development of the dopaminergic system." Development 141(3): 661–673.

Braun, E., M. Danan-Gotthold, L. E. Borm, K. W. Lee, E. Vinsland, P. Lonnerberg, L. Hu, X. Li, X. He, Z. Andrusivova, J. Lundeberg, R. A. Barker, E. Arenas, E. Sundstrom and S. Linnarsson (2023). "Comprehensive cell atlas of the first-trimester developing human brain." Science 382(6667): eadf1226.

Cable, D. M., E. Murray, V. Shanmugam, S. Zhang, L. S. Zou, M. Diao, H. Chen, E. Z. Macosko, R. A. Irizarry and F. Chen (2022). "Cell type-specific inference of differential expression in spatial transcriptomics." Nat Methods 19(9): 1076–1087.

Caminero, C. (2022). Neuroanatomy, Mesencephalon Midbrain.

Cang, Z., Y. Zhao, A. A. Almet, A. Stabell, R. Ramos, M. V. Plikus, S. X. Atwood and Q. Nie (2023). "Screening cell-cell communication in spatial transcriptomics via collective optimal transport." Nat Methods 20(2): 218–228.

Cao, J., M. Spielmann, X. Qiu, X. Huang, D. M. Ibrahim, A. J. Hill, F. Zhang, S. Mundlos, L. Christiansen, F. J. Steemers, C. Trapnell and J. Shendure (2019). "The single-cell transcriptional landscape of mammalian organogenesis." Nature 566(7745): 496–502.

Carola, G., D. Malagarriga, C. Calatayud, M. Pons-Espinal, L. Blasco-Agell, Y. Richaud-Patin, I. Fernandez-Carasa, V. Baruffi, S. Beltramone, E. Molina, P. Dell’Era, J. J. Toledo-Aral, E. Tolosa, A. R. Muotri, J. Garcia Ojalvo, J. Soriano, A. Raya and A. Consiglio (2021). "Parkinson’s disease patient-specific neuronal networks carrying the LRRK2 G2019S mutation unveil early functional alterations that predate neurodegeneration." NPJ Parkinsons Dis 7(1): 55.

Dann, E., N. C. Henderson, S. A. Teichmann, M. D. Morgan and J. C. Marioni (2022). "Differential abundance testing on single-cell data using k-nearest neighbor graphs." Nat Biotechnol 40(2): 245–253.

Elkjaer, M. L., T. Frisch, R. Reynolds, T. Kacprowski, M. Burton, T. A. Kruse, M. Thomassen, J. Baumbach and Z. Illes (2019). "Molecular signature of different lesion types in the brain white matter of patients with progressive multiple sclerosis." Acta Neuropathol Commun 7(1): 205.

Eze, U. C., A. Bhaduri, M. Haeussler, T. J. Nowakowski and A. R. Kriegstein (2021). "Single-cell atlas of early human brain development highlights heterogeneity of human neuroepithelial cells and early radial glia." Nat Neurosci 24(4): 584–594.

Faissner, A. (2023). "Low-density lipoprotein receptor-related protein-1 (LRP1) in the glial lineage modulates neuronal excitability." Front Netw Physiol 3: 1190240.

Falcon, S. and R. Gentleman (2007). "Using GOstats to test gene lists for GO term association." Bioinformatics 23(2): 257–258.

Ferri, A. L., W. Lin, Y. E. Mavromatakis, J. C. Wang, H. Sasaki, J. A. Whitsett and S. L. Ang (2007). "Foxa1 and Foxa2 regulate multiple phases of midbrain dopaminergic neuron development in a dosage-dependent manner." Development 134(15): 2761–2769.

Fiorenzano, A., E. Sozzi, M. Birtele, J. Kajtez, J. Giacomoni, F. Nilsson, A. Bruzelius, Y. Sharma, Y. Zhang, B. Mattsson, J. Emneus, D. R. Ottosson, P. Storm and M. Parmar (2021). "Single-cell transcriptomics captures features of human midbrain development and dopamine neuron diversity in brain organoids." Nat Commun 12(1): 7302.

Fujikawa, A., Y. Noda, H. Yamamoto, N. Tanga, G. Sakaguchi, S. Hattori, W. J. Song, I. Sora, T. Nabeshima, G. Katsuura and M. Noda (2019). "Mice deficient in protein tyrosine phosphatase receptor type Z (PTPRZ) show reduced responsivity to methamphetamine despite an enhanced response to novelty." PLoS One 14(8): e0221205.

Funk, M. C., A. N. Bera, T. Menchen, G. Kuales, K. Thriene, S. S. Lienkamp, J. Dengjel, H. Omran, M. Frank and S. J. Arnold (2015). "Cyclin O (Ccno) functions during deuterosome-mediated centriole amplification of multiciliated cells." EMBO J 34(8): 1078–1089.

Gantner, C. W., A. Cota-Coronado, L. H. Thompson and C. L. Parish (2020). "An Optimized Protocol for the Generation of Midbrain Dopamine Neurons under Defined Conditions." STAR Protoc 1(2): 100065.

Garritsen, O., E. Y. van Battum, L. M. Grossouw and R. J. Pasterkamp (2023). "Development, wiring and function of dopamine neuron subtypes." Nat Rev Neurosci 24(3): 134–152.

Gene Ontology, C., S. A. Aleksander, J. Balhoff, S. Carbon, J. M. Cherry, H. J. Drabkin, D. Ebert, M. Feuermann, P. Gaudet, N. L. Harris, D. P. Hill, R. Lee, H. Mi, S. Moxon, C. J. Mungall, A. Muruganugan, T. Mushayahama, P. W. Sternberg, P. D. Thomas, K. Van Auken, J. Ramsey, D. A. Siegele, R. L. Chisholm, P. Fey, M. C. Aspromonte, M. V. Nugnes, F. Quaglia, S. Tosatto, M. Giglio, S. Nadendla, G. Antonazzo, H. Attrill, G. Dos Santos, S. Marygold, V. Strelets, C. J. Tabone, J. Thurmond, P. Zhou, S. H. Ahmed, P. Asanitthong, D. Luna Buitrago, M. N. Erdol, M. C. Gage, M. Ali Kadhum, K. Y. C. Li, M. Long, A. Michalak, A. Pesala, A. Pritazahra, S. C. C. Saverimuttu, R. Su, K. E. Thurlow, R. C. Lovering, C. Logie, S. Oliferenko, J. Blake, K. Christie, L. Corbani, M. E. Dolan, H. J. Drabkin, D. P. Hill, L. Ni, D. Sitnikov, C. Smith, A. Cuzick, J. Seager, L. Cooper, J. Elser, P. Jaiswal, P. Gupta, P. Jaiswal, S. Naithani, M. Lera-Ramirez, K. Rutherford, V. Wood, J. L. De Pons, M. R. Dwinell, G. T. Hayman, M. L. Kaldunski, A. E. Kwitek, S. J. F. Laulederkind, M. A. Tutaj, M. Vedi, S. J. Wang, P. D’Eustachio, L. Aimo, K. Axelsen, A. Bridge, N. Hyka-Nouspikel, A. Morgat, S. A. Aleksander, J. M. Cherry, S. R. Engel, K. Karra, S. R. Miyasato, R. S. Nash, M. S. Skrzypek, S. Weng, E. D. Wong, E. Bakker, T. Z. Berardini, L. Reiser, A. Auchincloss, K. Axelsen, G. Argoud-Puy, M. C. Blatter, E. Boutet, L. Breuza, A. Bridge, C. Casals-Casas, E. Coudert, A. Estreicher, M. Livia Famiglietti, M. Feuermann, A. Gos, N. Gruaz-Gumowski, C. Hulo, N. Hyka-Nouspikel, F. Jungo, P. Le Mercier, D. Lieberherr, P. Masson, A. Morgat, I. Pedruzzi, L. Pourcel, S. Poux, C. Rivoire, S. Sundaram, A. Bateman, E. Bowler-Barnett, A. J. H. Bye, P. Denny, A. Ignatchenko, R. Ishtiaq, A. Lock, Y. Lussi, M. Magrane, M. J. Martin, S. Orchard, P. Raposo, E. Speretta, N. Tyagi, K. Warner, R. Zaru, A. D. Diehl, R. Lee, J. Chan, S. Diamantakis, D. Raciti, M. Zarowiecki, M. Fisher, C. James-Zorn, V. Ponferrada, A. Zorn, S. Ramachandran, L. Ruzicka and M. Westerfield (2023). "The Gene Ontology knowledgebase in 2023." Genetics 224(1).

Gillen, A. E., H. M. Brechbuhl, T. M. Yamamoto, E. Kline, M. M. Pillai, J. R. Hesselberth and P. Kabos (2017). "Alternative Polyadenylation of PRELID1 Regulates Mitochondrial ROS Signaling and Cancer Outcomes." Mol Cancer Res 15(12): 1741–1751.

Green, D. R. (2005). "Apoptotic pathways: ten minutes to dead." Cell 121(5): 671–674.

Gupta, D. (2017). Chapter 1 - Neuroanatomy.

Hanaway, J., J. A. McConnell and M. G. Netsky (1971). "Histogenesis of the substantia nigra, ventral tegmental area of Tsai and interpeduncular nucleus: an autoradiographic study of the mesencephalon in the rat." J Comp Neurol 142(1): 59–73.

Hao, Y., S. Hao, E. Andersen-Nissen, W. M. Mauck, 3rd, S. Zheng, A. Butler, M. J. Lee, A. J. Wilk, C. Darby, M. Zager, P. Hoffman, M. Stoeckius, E. Papalexi, E. P. Mimitou, J. Jain, A. Srivastava, T. Stuart, L. M. Fleming, B. Yeung, A. J. Rogers, J. M. McElrath, C. A. Blish, R. Gottardo, P. Smibert and R. Satija (2021). "Integrated analysis of multimodal single-cell data." Cell 184(13): 3573–3587 e3529.

Hao, Y., T. Stuart, M. H. Kowalski, S. Choudhary, P. Hoffman, A. Hartman, A. Srivastava, G. Molla, S. Madad, C. Fernandez-Granda and R. Satija (2024). "Dictionary learning for integrative, multimodal and scalable single-cell analysis." Nat Biotechnol 42(2): 293–304.

Hegarty, S. V., A. M. Sullivan and G. W. O’Keeffe (2013). "Midbrain dopaminergic neurons: a review of the molecular circuitry that regulates their development." Dev Biol 379(2): 123–138.

Jin, S., C. F. Guerrero-Juarez, L. Zhang, I. Chang, R. Ramos, C. H. Kuan, P. Myung, M. V. Plikus and Q. Nie (2021). "Inference and analysis of cell-cell communication using CellChat." Nat Commun 12(1): 1088.

Jo, J., Y. Xiao, A. X. Sun, E. Cukuroglu, H. D. Tran, J. Goke, Z. Y. Tan, T. Y. Saw, C. P. Tan, H. Lokman, Y. Lee, D. Kim, H. S. Ko, S. O. Kim, J. H. Park, N. J. Cho, T. M. Hyde, J. E. Kleinman, J. H. Shin, D. R. Weinberger, E. K. Tan, H. S. Je and H. H. Ng (2016). "Midbrain-like Organoids from Human Pluripotent Stem Cells Contain Functional Dopaminergic and Neuromelanin-Producing Neurons." Cell Stem Cell 19(2): 248–257.

Johns, P. (2014). Chapter 3 - Functional neuroanatomy.

Jung, C. G., H. Hida, K. Nakahira, K. Ikenaka, H. J. Kim and H. Nishino (2004). "Pleiotrophin mRNA is highly expressed in neural stem (progenitor) cells of mouse ventral mesencephalon and the product promotes production of dopaminergic neurons from embryonic stem cell-derived nestin-positive cells." FASEB J 18(11): 1237–1239.

Kang, H. M., M. Subramaniam, S. Targ, M. Nguyen, L. Maliskova, E. McCarthy, E. Wan, S. Wong, L. Byrnes, C. M. Lanata, R. E. Gate, S. Mostafavi, A. Marson, N. Zaitlen, L. A. Criswell and C. J. Ye (2018). "Multiplexed droplet single-cell RNA-sequencing using natural genetic variation." Nat Biotechnol 36(1): 89–94.

Kawano, H., K. Ohyama, K. Kawamura and I. Nagatsu (1995). "Migration of dopaminergic neurons in the embryonic mesencephalon of mice." Brain Res Dev Brain Res 86(1-2): 101–113.

Kirkeby, A., S. Grealish, D. A. Wolf, J. Nelander, J. Wood, M. Lundblad, O. Lindvall and M. Parmar (2012). "Generation of regionally specified neural progenitors and functional neurons from human embryonic stem cells under defined conditions." Cell Rep 1(6): 703–714.

Kleshchevnikov, V., A. Shmatko, E. Dann, A. Aivazidis, H. W. King, T. Li, R. Elmentaite, A. Lomakin, V. Kedlian, A. Gayoso, M. S. Jain, J. S. Park, L. Ramona, E. Tuck, A. Arutyunyan, R. Vento-Tormo, M. Gerstung, L. James, O. Stegle and O. A. Bayraktar (2022). "Cell2location maps fine-grained cell types in spatial transcriptomics." Nat Biotechnol 40(5): 661–671.

Kolberg, L., U. Raudvere, I. Kuzmin, P. Adler, J. Vilo and H. Peterson (2023). "g:Profiler-interoperable web service for functional enrichment analysis and gene identifier mapping (2023 update)." Nucleic Acids Res 51(W1): W207–W212.

Korsunsky, I., N. Millard, J. Fan, K. Slowikowski, F. Zhang, K. Wei, Y. Baglaenko, M. Brenner, P. R. Loh and S. Raychaudhuri (2019). "Fast, sensitive and accurate integration of single-cell data with Harmony." Nat Methods 16(12): 1289–1296.

Kurian, M. A., P. Gissen, M. Smith, S. Heales, Jr. and P. T. Clayton (2011). "The monoamine neurotransmitter disorders: an expanding range of neurological syndromes." Lancet Neurol 10(8): 721–733.

Kurian, M. A., Y. Li, J. Zhen, E. Meyer, N. Hai, H. J. Christen, G. F. Hoffmann, P. Jardine, A. von Moers, S. R. Mordekar, F. O’Callaghan, E. Wassmer, E. Wraige, C. Dietrich, T. Lewis, K. Hyland, S. Heales, Jr., T. Sanger, P. Gissen, B. E. Assmann, M. E. Reith and E. R. Maher (2011). "Clinical and molecular characterisation of hereditary dopamine transporter deficiency syndrome: an observational cohort and experimental study." Lancet Neurol 10(1): 54–62.

La Manno, G., D. Gyllborg, S. Codeluppi, K. Nishimura, C. Salto, A. Zeisel, L. E. Borm, S. R. W. Stott, E. M. Toledo, J. C. Villaescusa, P. Lonnerberg, J. Ryge, R. A. Barker, E. Arenas and S. Linnarsson (2016). "Molecular Diversity of Midbrain Development in Mouse, Human, and Stem Cells." Cell 167(2): 566–580 e519.

Lancaster, M. A., M. Renner, C. A. Martin, D. Wenzel, L. S. Bicknell, M. E. Hurles, T. Homfray, J. M. Penninger, A. P. Jackson and J. A. Knoblich (2013). "Cerebral organoids model human brain development and microcephaly." Nature 501(7467): 373–379.

Li, Y., Z. Li, C. Wang, M. Yang, Z. He, F. Wang, Y. Zhang, R. Li, Y. Gong, B. Wang, B. Fan, C. Wang, L. Chen, H. Li, P. Shi, N. Wang, Z. Wei, Y. L. Wang, L. Jin, P. Du, J. Dong and J. Jiao (2023). "Spatiotemporal transcriptome atlas reveals the regional specification of the developing human brain." Cell 186(26): 5892–5909 e5822.

Maffezzini, C., J. Calvo-Garrido, A. Wredenberg and C. Freyer (2020). "Metabolic regulation of neurodifferentiation in the adult brain." Cell Mol Life Sci 77(13): 2483–2496.

Marklund, U., Z. Alekseenko, E. Andersson, S. Falci, M. Westgren, T. Perlmann, A. Graham, E. Sundstrom and J. Ericson (2014). "Detailed expression analysis of regulatory genes in the early developing human neural tube." Stem Cells Dev 23(1): 5–15.

Monzel, A. S., L. M. Smits, K. Hemmer, S. Hachi, E. L. Moreno, T. van Wuellen, J. Jarazo, J. Walter, I. Bruggemann, I. Boussaad, E. Berger, R. M. T. Fleming, S. Bolognin and J. C. Schwamborn (2017). "Derivation of Human Midbrain-Specific Organoids from Neuroepithelial Stem Cells." Stem Cell Reports 8(5): 1144–1154.

Moran, P. A. (1950). "Notes on continuous stochastic phenomena." Biometrika 37(1-2): 17–23.

Mourlevat, S., T. Debeir, J. E. Ferrario, J. Delbe, D. Caruelle, O. Lejeune, C. Depienne, J. Courty, R. Raisman-Vozari and M. Ruberg (2005). "Pleiotrophin mediates the neurotrophic effect of cyclic AMP on dopaminergic neurons: analysis of suppression-subtracted cDNA libraries and confirmation in vitro." Exp Neurol 194(1): 243–254.

Nelander, J., J. B. Hebsgaard and M. Parmar (2009). "Organization of the human embryonic ventral mesencephalon." Gene Expr Patterns 9(8): 555–561.

Ng, J., S. Barral, C. De La Fuente Barrigon, G. Lignani, F. A. Erdem, R. Wallings, R. Privolizzi, G. Rossignoli, H. Alrashidi, S. Heasman, E. Meyer, A. Ngoh, S. Pope, R. Karda, D. Perocheau, J. Baruteau, N. Suff, J. Antinao Diaz, S. Schorge, J. Vowles, L. R. Marshall, S. A. Cowley, S. Sucic, M. Freissmuth, J. R. Counsell, R. Wade-Martins, S. J. R. Heales, A. A. Rahim, M. Bencze, S. N. Waddington and M. A. Kurian (2021). "Gene therapy restores dopamine transporter expression and ameliorates pathology in iPSC and mouse models of infantile parkinsonism." Sci Transl Med 13(594).

Ng, M. Y. W., T. Wai and A. Simonsen (2021). "Quality control of the mitochondrion." Dev Cell 56(7): 881–905.

Nickels, S. L., J. Modamio, B. Mendes-Pinheiro, A. S. Monzel, F. Betsou and J. C. Schwamborn (2020). "Reproducible generation of human midbrain organoids for in vitro modeling of Parkinson’s disease." Stem Cell Res 46: 101870.

Novak, G., D. Kyriakis, K. Grzyb, M. Bernini, S. Rodius, G. Dittmar, S. Finkbeiner and A. Skupin (2022). "Single-cell transcriptomics of human iPSC differentiation dynamics reveal a core molecular network of Parkinson’s disease." Commun Biol 5(1): 49.

Parraga, R. G., L. L. Possatti, R. V. Alves, G. C. Ribas, U. Ture and E. de Oliveira (2016). "Microsurgical anatomy and internal architecture of the brainstem in 3D images: surgical considerations." J Neurosurg 124(5): 1377–1395.

Quadrato, G., T. Nguyen, E. Z. Macosko, J. L. Sherwood, S. Min Yang, D. R. Berger, N. Maria, J. Scholvin, M. Goldman, J. P. Kinney, E. S. Boyden, J. W. Lichtman, Z. M. Williams, S. A. McCarroll and P. Arlotta (2017). "Cell diversity and network dynamics in photosensitive human brain organoids." Nature 545(7652): 48–53.

Reyes, S., Y. Fu, K. Double, L. Thompson, D. Kirik, G. Paxinos and G. M. Halliday (2012). "GIRK2 expression in dopamine neurons of the substantia nigra and ventral tegmental area." J Comp Neurol 520(12): 2591–2607.

Ricard-Blum, S. (2011). "The collagen family." Cold Spring Harb Perspect Biol 3(1): a004978.

Roll, S., J. Seul, M. Paulsson and U. Hartmann (2006). "Testican-1 is dispensable for mouse development." Matrix Biol 25(6): 373–381.

Rossignoli, G., K. Kramer, E. Lugara, H. Alrashidi, S. Pope, C. De La Fuente Barrigon, K. Barwick, G. Bisello, J. Ng, J. Counsell, G. Lignani, S. J. R. Heales, M. Bertoldi, S. Barral and M. A. Kurian (2021). "Aromatic l-amino acid decarboxylase deficiency: a patient-derived neuronal model for precision therapies." Brain 144(8): 2443–2456.

Siletti, K., R. Hodge, A. Mossi Albiach, K. W. Lee, S. L. Ding, L. Hu, P. Lonnerberg, T. Bakken, T. Casper, M. Clark, N. Dee, J. Gloe, D. Hirschstein,N. V. Shapovalova, C. D. Keene, J. Nyhus, H. Tung, A. M. Yanny, E. Arenas, E. S. Lein and S. Linnarsson (2023). "Transcriptomic diversity of cell types across the adult human brain." Science 382(6667): eadd7046.

Street, K., D. Risso, R. B. Fletcher, D. Das, J. Ngai, N. Yosef, E. Purdom and S. Dudoit (2018). "Slingshot: cell lineage and pseudotime inference for single-cell transcriptomics." BMC Genomics 19(1): 477.

Tekin, H., S. Simmons, B. Cummings, L. Gao, X. Adiconis, C. C. Hession, A. Ghoshal, D. Dionne, S. R. Choudhury, V. Yesilyurt, N. E. Sanjana, X. Shi, C. Lu, M. Heidenreich, J. Q. Pan, J. Z. Levin and F. Zhang (2018). "Effects of 3D culturing conditions on the transcriptomic profile of stem-cell-derived neurons." Nat Biomed Eng 2(7): 540–554.

Tiklova, K., A. K. Bjorklund, L. Lahti, A. Fiorenzano, S. Nolbrant, L. Gillberg, N. Volakakis, C. Yokota, M. M. Hilscher, T. Hauling, F. Holmstrom, E. Joodmardi, M. Nilsson, M. Parmar and T. Perlmann (2019). "Single-cell RNA sequencing reveals midbrain dopamine neuron diversity emerging during mouse brain development." Nat Commun 10(1): 581.

Trapnell, C., D. Cacchiarelli, J. Grimsby, P. Pokharel, S. Li, M. Morse, N. J. Lennon, K. J. Livak, T. S. Mikkelsen and J. L. Rinn (2014). "The dynamics and regulators of cell fate decisions are revealed by pseudotemporal ordering of single cells." Nat Biotechnol 32(4): 381–386.

Traxler, L., J. Lagerwall, S. Eichhorner, D. Stefanoni, A. D’Alessandro and J. Mertens (2021). "Metabolism navigates neural cell fate in development, aging and neurodegeneration." Dis Model Mech 14(8).

Vadasz, C., J. F. Smiley, K. Figarsky, M. Saito, R. Toth, B. M. Gyetvai, M. Oros, K. K. Kovacs, P. Mohan and R. Wang (2007). "Mesencephalic dopamine neuron number and tyrosine hydroxylase content: Genetic control and candidate genes." Neuroscience 149(3): 561–572.

Vaswani, A. R., B. Weykopf, C. Hagemann, H. U. Fried, O. Brustle and S. Blaess (2019). "Correct setup of the substantia nigra requires Reelin-mediated fast, laterally-directed migration of dopaminergic neurons." Elife 8.

Waldvogel, H. J., M. A. Curtis, K. Baer, M. I. Rees and R. L. Faull (2006). "Immunohistochemical staining of post-mortem adult human brain sections." Nat Protoc 1(6): 2719–2732.

Williams, C. G., H. J. Lee, T. Asatsuma, R. Vento-Tormo and A. Haque (2022). "An introduction to spatial transcriptomics for biomedical research." Genome Med 14(1): 68.

Winkler, C. and S. Yao (2014). "The midkine family of growth factors: diverse roles in nervous system formation and maintenance." Br J Pharmacol 171(4): 905–912.

Yu, Y., Z. Zeng, D. Xie, R. Chen, Y. Sha, S. Huang, W. Cai, W. Chen, W. Li, R. Ke and T. Sun (2021). "Interneuron origin and molecular diversity in the human fetal brain." Nat Neurosci 24(12): 1745–1756.

Zagare, A., K. Barmpa, S. Smajic, L. M. Smits, K. Grzyb, A. Grunewald, A. Skupin, S. L. Nickels and J. C. Schwamborn (2022). "Midbrain organoids mimic early embryonic neurodevelopment and recapitulate LRRK2-p.Gly2019Ser-associated gene expression." Am J Hum Genet 109(2): 311–327.

